# Multi-omic profiling of human antibody-secreting cells reveals diverse subsets sustain durable humoral immunity

**DOI:** 10.64898/2026.04.13.717827

**Authors:** David R. Glass, Elisabeth Dornisch, Hang Yin, Susan A. Ludmann, Ashwin Samudre, Sydney Kuhl, Jocelin Malone, Aishwarya Chander, Saransh N. Kaul, Cole G. Phalen, Vaishnavi Parthasarathy, Medbh A. Dillon, Palak Genge, Tyanna J. Stuckey, Stephanie Anover-Sombke, Peter J. Wittig, Mark-Phillip Pebworth, Ziyuan He, Kathy E. Henderson, Priya Ravisankar, Veronica Hernandez, Blessing Musgrove, Suraj Mishra, Upaasana Krishnan, Zach Thomson, Morgan Weiss, Nina Estep, Lucas Graybuck, Melinda Angus-Hill, Claire E. Gustafson, Mackenzie Kopp, Julian Reading, Xiao-jun Li, Matheus P. Viana, Thomas F. Bumol, Ananda W. Goldrath, Mikael Sigvardsson, Sean C. Bendall, Peter J. Skene, Damian J. Green, Evan W. Newell, Troy Torgerson, Marla C. Glass

## Abstract

Antibody-secreting cells (ASCs) provide humoral immunity that can mediate lifelong protection against pathogens. Current classifications cannot delineate the heterogenous functionalities, tissue residencies, and lifespans of human ASC subsets, impeding clinical translation. We applied multi-omic sequencing, spatial proteomics, and functional assays to discover and characterize human bone marrow (BM) ASC subsets. We identified two peripheral subsets (ASCp) also present in blood and three BM-resident subsets (ASCr), comprising a maturation continuum associated with increased mitochondrial networking, diminished antibody secretion, differential transcription factor motif accessibility, and preferential co-localization in homotypic niches. CD19+9+ASCr and CD19-ASCr exhibited poor recovery years after BM transplantation, indicating a strong dependence on supportive niches. Childhood vaccine antigens were recognized by long-lived ASCr subsets in adults and by immature HLA-DR+ASCp, implying ASCs can differentiate without recent antigen exposure. Our results provide new insights into ASC identity, maturation, and longevity and a generalizable framework for study and manipulation of human ASCs.

## INTRODUCTION

During immune responses, ASCs differentiate from activated B cells and mediate immune protection through immunoglobulin (Ig) secretion and other effector functions^1–3^. Each ASC secretes one of the five different isotypes of Ig heavy chains which modulate global immune responses. Most ASCs survive only hours or days after differentiation, but a fraction persist in the BM and other tissues to constitutively secrete antibodies for months, years, or decades^4–6^. These long-lived ASCs undergo metabolic and transcriptomic adaptations and occupy specialized niches within the BM that provide survival signals to sustain their longevity^7,8^. Long-lived ASCs have been extensively studied in mice, with seminal findings including their dependence on mitochondrial pyruvate import for long-term survival^9^, the progressive development of long-lived ASC transcriptional programs^10^, and localized chemokine cues guiding their retention within supportive niches^11,12^.

Less is known about human ASC biology, owing in part to a lack of markers that segregate human ASC subsets^13^. Most studies discriminate ASC subsets using only CD19 and CD138, which is unreliable after cryopreservation and therefore unsuitable for retrospective human studies^14,15^. Controversy remains about whether CD138+CD19-ASCs exclusively hold long-lived immunity^16^ or whether they merely represent one of multiple long-lived populations^17–19^. Human BM ASCs are transcriptionally diverse, but current classification schemes fail to capture this heterogeneity^20^. This limits the possibilities to identify, ascribe function to, and manipulate distinct human ASC subsets. Bridging this gap is not only critical for advancing fundamental immunology, but also for improving human health, as ASCs mediate durable humoral protection after vaccination but contribute pathogenic antibodies in autoimmunity. Malignant ASCs cause multiple myeloma (MM), and patients routinely exhibit impaired immunity against pathogens owing both to the therapies and the disease itself^21^. Selective induction and targeted therapeutic intervention of ASCs in clinical settings will require an enhanced understanding of human ASC subset identities. To address this problem, we paired a systems immunology approach with a suite of functional assays to discover and comprehensively characterize human ASC subsets. Interactive tools to explore multi-modal datasets can be found at https://apps.allenimmunology.org/aifi/resources/antibody-secreting-cells/

## RESULTS

### Comprehensive multi-omic analysis identifies five human bone marrow ASC subsets

To identify cell surface molecules expressed by human ASCs, we compiled four high-dimensional cytometry datasets from healthy human tissues and manually gated ASCs and analytes (**Fig. 1A, Fig. S1A-B**). We incorporated published flow cytometry screen data run on spleen samples^22^ and mass cytometry data run on BM, lymph node, and tonsil samples^23^ (**Fig. S1C-D**). Additionally, we performed a mass cytometry screen^23^ on BM and ran mass cytometry on donor-matched BM and peripheral blood (PB) samples (**Fig. S1E-G, Table S1**). Across the datasets, we identified 134 unique surface molecules expressed by human ASCs (**Fig. 1B**).

**Figure 1:**
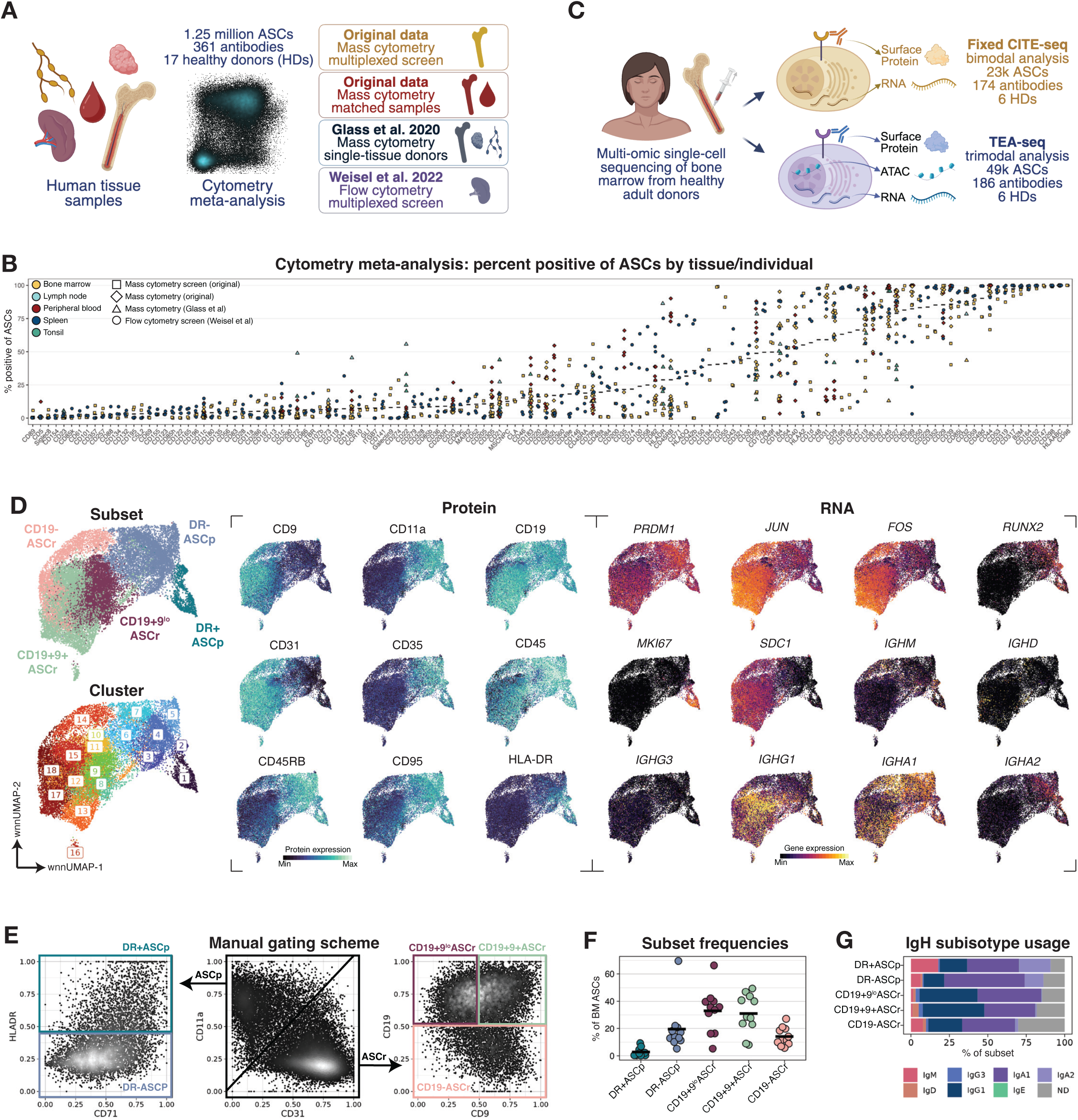
Comprehensive multi-omic analysis identifies five human bone marrow ASC subsets. A) Cytometry meta-analysis diagram. B) Percent positive of each cell surface molecule on total ASCs, by sample (dot), tissue (color) and dataset (shape). Crossbar indicates mean; only analytes in which at least one sample had >5% positivity for ASCs are shown. Data from flow and mass cytometry meta-analysis. C) Single-cell multi-omics diagram. D) wnnUMAPs of ASCs colored by subset (upper left), Seurat cluster (lower left), ADT expression (center), and GEX expression (right). Data from HD BM fixed CITEseq. E) Biaxial plots of ADT manual gating into subsets. Data from HD BM fixed CITEseq. F) Frequency of subset out of total ASCs by individual. Crossbar indicates mean. Data from HD BM fixed CITEseq and HD BM TEAseq. G) Frequency of subisotype usage by subset. ND denotes subisotype not determined. Data from HD BM fixed CITEseq.

Informed by these analyses, we developed large cellular indexing of transcriptomes and epitopes by sequencing (CITEseq) panels^24^ and performed multi-omic single-cell fixed CITEseq^25^ and transcripts, epitopes, and accessibility sequencing (TEAseq)^26^ on healthy human BM aspirate (BMA) samples (**Fig. 1C, Fig. S2A-E, Table S1-2**). As others have noted^27^, we found weak correlations (median r=0.16) between surface protein expression levels and their corresponding mRNA levels (**Fig. S2F-H**), indicating that the RNA and protein modalities capture complementary, non-redundant features of cell identity.

To discover human BM ASC subsets, we generated a weighted nearest-neighbor (wNN) graph informed by both protein and RNA expression^28^ and performed unsupervised clustering of total ASCs. As expected, ASCs expressed *PRDM1*, which encodes the ASC lineage-defining transcription factor (TF) Blimp-1, while other TFs displayed higher variance expression patterns amongst ASCs (**Fig. 1D**). Many surface proteins divided wNN clusters 1-7 and 8-18 into two distinct subtypes, peripheral ASCs (ASCp) and BM-resident ASCs (ASCr; **Fig 1D**, **Fig. S3A**). We coined this nomenclature due to the presence of ASCp in both BM and PB, discussed later.

Dozens of proteins were differentially expressed between subtypes including higher levels of CD11a, CD45RB, and CD35 in ASCp and higher levels of the adhesion molecules CD31, CD54, and CD49f in ASCr (**Fig. S3B**). To formalize a reproducible approach that captures the distinction between ASCp and ASCr and enables more granular dissection of ASC identity, we developed a manual gating scheme (**Fig. 1E**). ASCp are defined as CD11a+CD31- and further stratified by HLA-DR expression as DR+ASCp and DR-ASCp, and ASCr are defined as CD11a-CD31+ and further stratified by CD19 and CD9 expression as CD19+9^lo^ASCr, CD19+9+ASCr, and CD19-ASCr. We manually gated ASCs and each cluster was assigned to the subset in which the plurality of their cells fell into the corresponding gate (**Fig. S3C**). For all sequencing data, we refer to this identity label as “subset,” while manual gating labels are referred to as “gate.” Distinct features of these ASC subsets were illustrated by UMAPs plots with overlays for cellular identity, protein expression, gene expression, and gene set expression (**Fig. S4**).

All ASC subsets were routinely observed in healthy adult BMAs (**Fig. 1F**). A wide range of immunoglobulin heavy chain (IgH) subisotypes were used by all ASC subsets, though IgA2 was enriched in ASCp compared to ASCr (**Fig. 1G**). CD138, encoded by *SDC1*, has historically been used to identify more mature ASC subsets^29^ but poorly discriminates single-cell RNA-sequencing (scRNAseq) clusters^20^ and is unreliable after cryopreservation^14,15^. *SDC1* gene (CITEseq data) and CD138 protein (flow cytometry data) expression were enriched among ASCr subsets but did not completely delineate any subset (**Fig. S3D-F**). Therefore, subset identities in this study align with descriptions of CD138 as a maturation marker but delineate ASCs into populations that are not recapitulated using CD138 expression. In summary, we identified five human BM ASC subsets with distinctive proteomic and transcriptional identities.

### Ig heavy chain subisotype captures a facet of ASC identity

Each ASC uses one heavy chain (sub)isotype and one light chain isotype for secreted and surface-bound Ig. Light chain isotype usage was similar between subsets (**Fig. S5A**). All ASCs exhibited at least some surface Ig expression, as indicated by the presence of separate surface Ig kappa (IgK)^lo^ and surface Ig lambda (IgL)^lo^ populations rather than a single IgK-IgL-population (**Fig. S5B**). Surface Ig expression (quantified using IgK and IgL staining) was more strongly associated with ASC heavy chain subisotype usage rather than cell subset identity, as we previously observed for human B cells^23^ (**Fig. S5C-D**).

ASC subsets used a wide range of IgH subisotypes (**Fig. 1G**). For most HDs, the plurality of ASCs used the IgA1 subisotype, though appreciable fractions (>2% mean) of ASCs used IgG1, IgA2, IgM, and even IgD (**Fig. S5E**). IgG2 and IgG4 RNA probes were not present in the fixed CITEseq experiment and likely contributed to the fraction of subisotype identities not determined (ND; **Fig. S2B**). We identified thirteen IgE ASCs in the fixed CITEseq dataset and 81 in the TEAseq dataset, representing mean 0.06% and 0.16% of BM ASCs, respectively. When delineating ASCs by subisotype usage, we found that most subisotypes were primarily comprised of ASCr phenotypes, except IgM and IgA2 which were majority ASCp phenotypes (**Fig. S5F**). IgM, IgD, and IgA2 subisotypes had the greatest quantity of DEPs (**Fig. S5G-H**).

Notably, IgD ASCs did not co-express IgM, suggesting these are non-canonically class-switched ‘IgD-only’ ASCs^30,31^. As previously observed on type 2-polarized memory B cells that hold allergic memory^32,33^, IgE ASCs expressed CD23, the low-affinity IgE receptor, and high levels of CD40. These cells also expressed high levels of the inhibitory receptor CD200/OX-2. IgM, IgD, and IgA2 ASCs had the greatest quantity of DEGs (**Fig. S5I**), with several overlapping genes significantly upregulated in IgM and IgA2, including *JCHAIN*, *CCR9*, and *TNFRSF1B* (TNFR2; **Fig. S5J**). Collectively, IgA1 was the most prevalently used subisotype, IgM and IgA2 ASCs had distinctive expression patterns, and IgD and rare IgE ASCs were observed in multiple HDs.

### ASCs alter adhesion molecule expression and transcriptional regulators during bone marrow maturation

To interrogate global differences between subsets, we calculated the geodesic distance between cells using a weighted shared nearest neighbor (wSNN) graph, which incorporates both protein and RNA expression information. ASCp subsets were more similar to other ASCp subsets and ASCr subsets were more similar to other ASCr subsets (**Fig. 2A**). The subsets largely formed a phenotypic continuum starting from DR+ASCp, through DR-ASCp, to CD19+9^lo^ASCr, and both CD19+9+ASCr and CD19-ASCr seemed to represent terminal maturation states, with similar distances to ASCp subsets (**Fig. S6A**).

**Figure 2:**
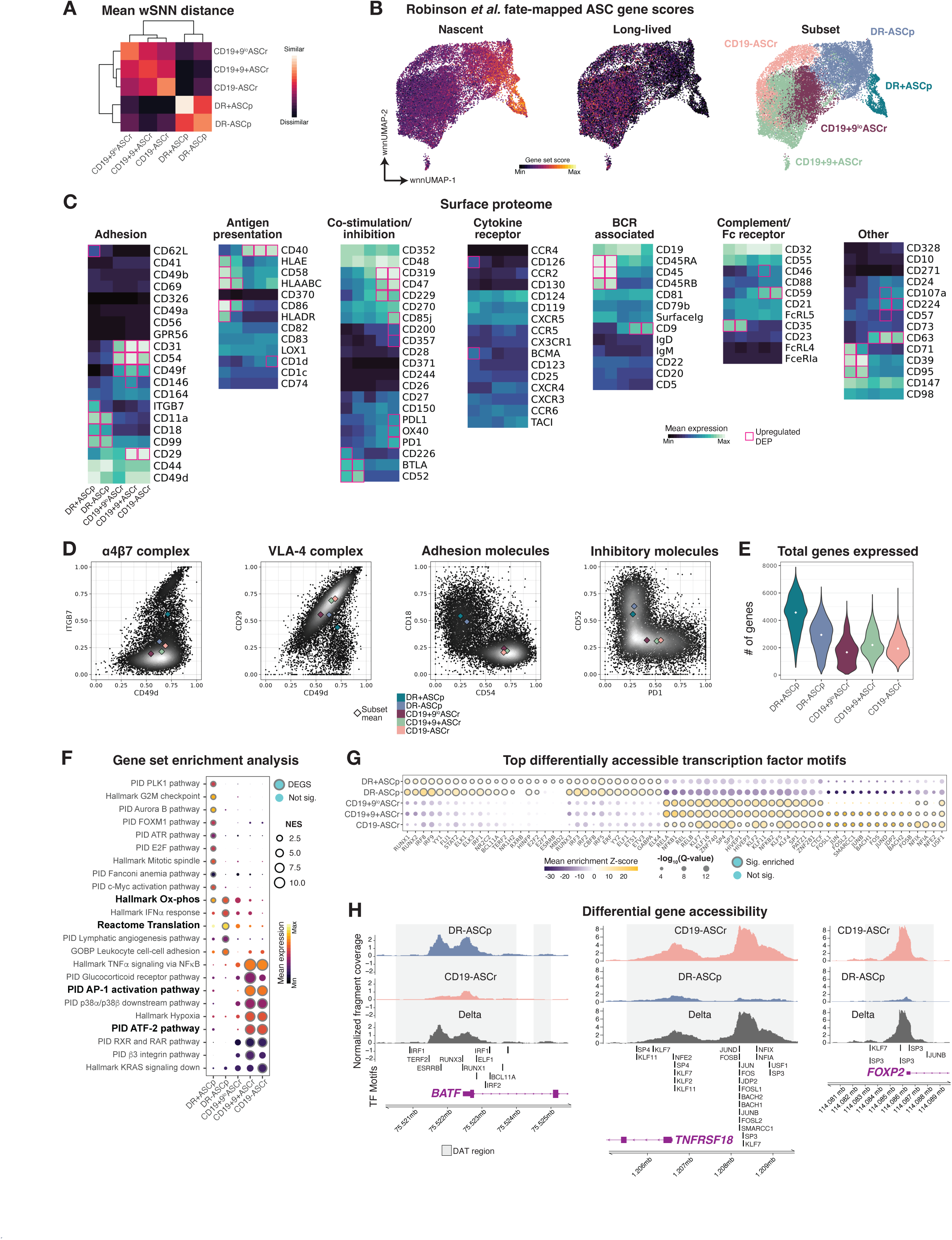
ASCs alter adhesion molecule expression and transcriptional regulators during bone marrow maturation. A) Mean of mean pairwise geodesic distances between all cells through the weighted shared nearest neighbor graph, by subset. Data from HD BM fixed CITEseq. B) wnnUMAPs of ASCs colored by gene set scores derived from DEGs between nascent (<60 days old) and long-lived (>60 days old) ASCs in a fate-mapped murine model^10^ (left/center) or subset (right). Data from HD BM fixed CITEseq. C) Mean expression of all ADTs by subset, grouped by protein function. Boxed values indicated significantly upregulated DEPs. Data from HD BM fixed CITEseq. D) Biaxial plot of indicated ADTs. Diamonds indicate subset mean, color indicates subset. Data from HD BM fixed CITEseq. E) The number of genes expressed by each cell, by subset. White dot indicates median. Data from HD BM fixed CITEseq. F) Mean expression of DEGSs (color) by subset. Outline indicates statistical significance; size indicates normalized enrichment score (NES). Bolded gene sets are highlighted in Fig. S6F. Data from HD BM fixed CITEseq G) Mean enrichment z-scores (color) by subset. Outline indicates statistical significance; size indicates -log_10_(Q-value). Data from HD BM TEAseq. H) Normalized fragment coverage (y-axis) of the indicated subsets and their delta by gene (panels). Grey shading indicates DATs and vertical lines indicate transcription factor motif loci. Gene diagrams’ (colored purple) arrows indicate transcriptional direction, short blocks indicate untranslated regions, and tall blocks indicate exons. Data from HD BM TEAseq.

To relate our ASC subsets to previously described long-lived phenotypes, we leveraged published differentially expressed genes (DEGs) from a murine fate-mapping model that distinguished nascent (<60 days old) from long-lived (>60 days old) BM ASCs^10^. We generated gene sets scores from the DEGs and as expected, the nascent gene set score was highest in DR+ASCp, followed by DR-ASCp, and equally low in all ASCr subsets (**Fig. 2B, S6B**). The long-lived gene set score was highest in CD19+9+ASCr, followed by the two other ASCr subsets, and lowest in the ASCp subsets. For both gene sets scores, the majority of genes were not differentially expressed between subsets, likely due to species level differences (**Fig. S6C**).

To discover features discriminating ASC subsets, we identified and quantified differentially expressed proteins (DEPs), DEGs, and differentially accessible transcription factor (DATF) motifs from fixed CITEseq and TEAseq data (**Fig. S6D, Table S3-5**). DEGs were used to generate gene set scores to identify ASC subsets in other single-cell datasets (**Fig. S4E, Table S6**). As with the distance metric, we observed protein expression often formed a continuum across subsets (**Fig. 2C**). Several proteins associated with antigen presentation were downregulated in ASCr, including the co-stimulatory molecule, CD86, and the nonclassical MHC molecule, HLA-E, while CD40 was upregulated in ASCr. We observed significant differences in adhesion molecule expression, with higher levels of CD11a and CD18 (which form the LFA-1 complex) in ASCp, and higher levels of its ligand, CD54, in ASCr. A subset of DR+ASCp co-expressed CD49d and ITGB7 to form the gut-homing α4β7 complex, whereas the other subsets upregulated CD29 to form the VLA-4 complex with CD49d, enabling retention in the BM^12^ (**Fig. 2D**). We also observed that BTLA and CD52, which restrain BCR signaling^34,35^, were significantly enriched in ASCp, while the checkpoint molecules, PD-L1 and PD-1, were significantly enriched in CD19-ASCr (**Fig. 2C-D**).

DR+ASCp expressed the highest quantity of total genes (**Fig. 2E**), total transcripts (**Fig. S6E**), and DEGs (**Fig. S6D**), suggesting these cells represent recently activated, immature ASCs, while CD19+9^lo^ASCr expressed the lowest quantity of each metric. Gene set enrichment analysis (GSEA)^36^ revealed ASCp subsets were enriched for oxidative phosphorylation and translation genes (**Fig. 2F, S6F, Table S7**), which may enable increased Ig production. CD19+9+ASCr and CD19-ASCr were enriched for genes in the AP-1 and ATF-2 pathways, which are implicated in stress response^37^ and may enable long-lasting constitutive Ig secretion.

To investigate how transcriptional regulators coordinate gene programs, we analyzed DATF motifs from the TEAseq dataset. ASCp were highly enriched for RUNX and IRF family motif accessibility, while ASCr were enriched for NFκB, KLF, and AP-1 family motif accessibility (**Fig. 2G**). Interestingly, these KLF family TFs can both enhance and suppress NFκB activity, suggesting an intrinsic regulatory balance to restrain proinflammatory responses among ASCr^38–40^. Visualization of motif footprints indicated *STAT2* and *IRF1* motifs were more accessible in ASCp with archetypal dips in reads at the TF-specific DNA binding site, indicating TF occupancy^41^ (**Fig. S6G**). *NFIX*, a TF shown to enhance murine hematopoietic stem cell survival^42^, had significantly greater TF motif accessibility in ASCr subsets, particularly CD19-ASCr. Accessible IRF1, IRF2, RUNX1, and RUNX3 motifs were evident proximal to *BATF* in DR-ASCp, but not CD19-ASCr (**Fig. 2H**). Alternatively, in CD19-ASCr but not DR-ASCp, KLF and AP-1 family motifs were accessible proximal to *FOXP2* and *TNFRSF18*, which encodes CD357/GITR. GITR has been shown to restrain proliferation of malignant ASCs in a cell-intrinsic manner^43^.

In summary, published gene sets from a fate-mapping murine model indicate that DR+ASCp represent the most immature subset, whereas CD19+9+ASCr displayed the highest long-lived gene set score. Across all five subsets, we observe a continuum of differences in adhesion molecules, inhibitory receptors, metabolic programs, and transcriptional regulators.

### Only ASCp cycle and their phenotype is stable through cell cycle phases

DR+ASCp were enriched for cycling cell gene programs (**Fig. 2F**), so we investigated if other subsets were actively cycling. We used the G2M checkpoint and E2F pathway gene scores to identify cycling cells, which coincided with *MKI67* expression (**Fig. S7A**). We calculated cell cycle phase labels^44^ and used them to generate continuous cell cycle axes^45^, which organized in the expected order: G1/S, S, G2, G2/M, M/G1 (**Fig. S7B**). We quantified the frequency of cycling cells by subset and found the majority of DR+ASCp, a very small fraction of DR-ASCp, and no ASCr were cycling (**Fig. S7C**). DR+ASCp were comprised of two wNN clusters; the majority of cluster 1 but only a small fraction of cluster 2 were cycling (**Fig. S7D**). Interestingly, cluster 2 instead displayed a mucosal-associated phenotype with high expression of *CCR9* and ITGB7, and an enrichment for the IgM and IgA2 subisotypes (**Fig. S7E-F**). Cycling cells were stable in their cell surface phenotype across cell cycle phases; no proteins were significantly differentially expressed between phases (**Fig. S7G**). In summary, only one cluster of DR+ASCp were highly enriched for cycling cells, which exhibited stable phenotypes across cell cycle phases.

### ASCr subsets have larger and more networked mitochondria

Having observed differences in protein expression, gene expression, and chromatin accessibility between subsets, we sought to establish if subsets differed in morphological features. We derived a sorting strategy replicating the manual gating scheme of our CITEseq data, sorted the five ASC subsets and transitional B cells (TBCs) from three HD BM samples, and imaged cells using differential interference contrast (DIC) microscopy (**Fig. 1E, 3A, S8A-B, Table S1-2**). DR+ASCp had the greatest median cell area, followed by CD19+9^lo^ASCr, DR-ASCp, CD19+9+ASCr, and CD19-ASCr (**Fig. 3B**). Likewise, CD19+9+ASCr, and CD19-ASCr had significantly lower cytoplasm area than other subsets (**Fig. S8C**). We observed cells containing vacuoles, which likely represent autophagosomes supporting sustainable Ig production^16,46^ (**Fig. 3A, S8D**). The mean percentage of cells with at least one vacuole was higher in ASCr subsets (**Fig. S8E**), and CD19+9+ASCr and CD19-ASCr contained significantly more vacuoles than ASCp subsets (**Fig. 3C**).

**Figure 3:**
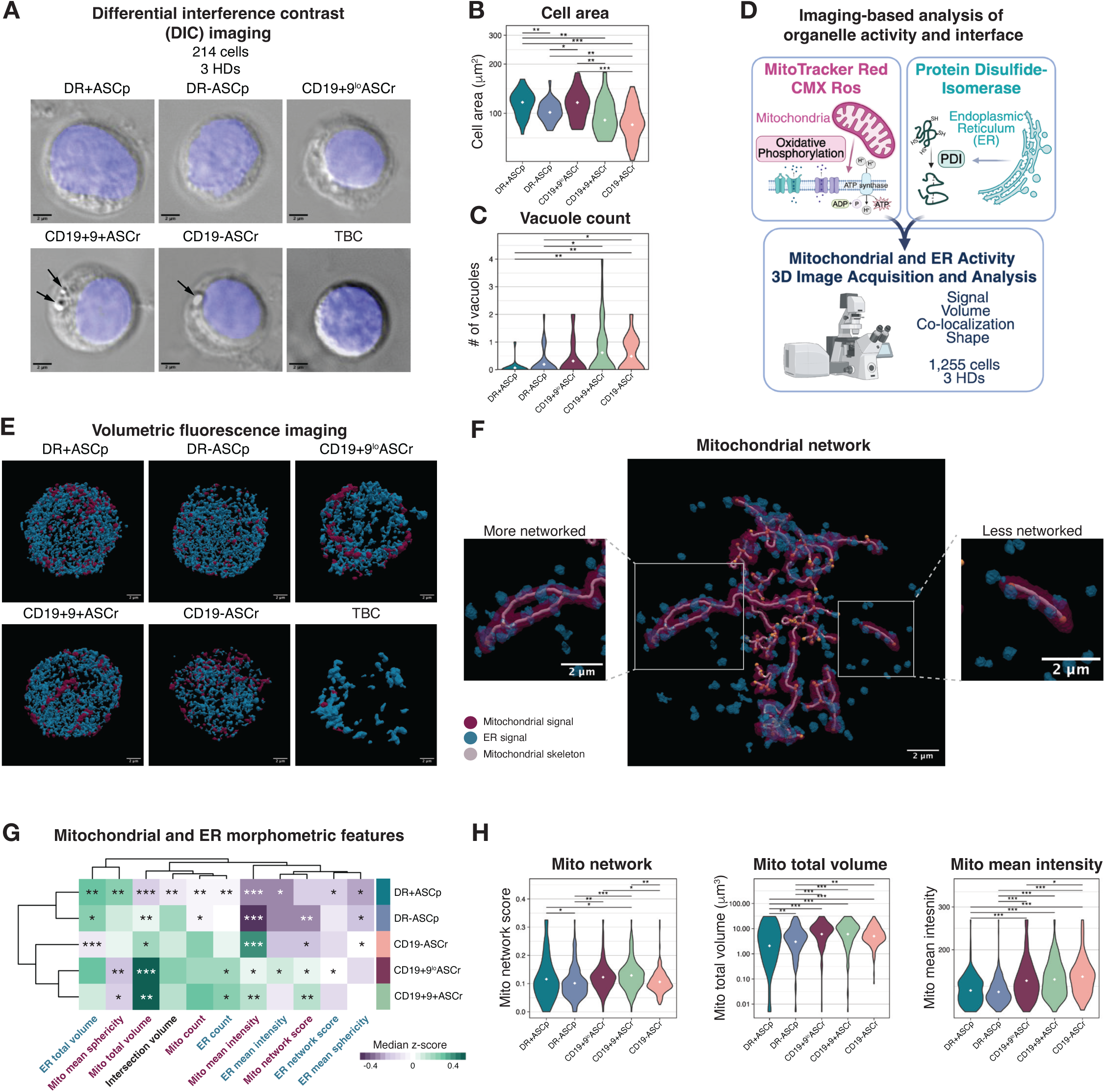
ASCr subsets have larger and more networked mitochondria. A) Representative DIC images (60x magnification) of ASC subsets and TBC sorted from three HD BM samples. B) Cell area by subset. White dot indicates median. Q-values calculated by FDR-corrected Wilcoxon rank sum test. *Q<0.1; **Q<0.01; ***Q<0.001. Data from HD BM DIC imaging. C) Vacuole counts by subset. White dot indicates mean. Q-values calculated by FDR-corrected Wilcoxon rank sum test. *Q<0.1; **Q<0.01. Data from HD BM DIC imaging. D) Experimental diagram E) Representative fluorescence volumetric images of sorted subsets (60x magnification) showing mitochondrial (pink) and endoplasmic reticulum (ER; teal) activity. Data from three HD BM samples. F) Representative images indicating more (left) and less (right) networked mitochondria. G) Median z-scores of morphometric features by subset from fluorescence mitochondrial and ER imaging. Q-values calculated using one-versus-all FDR-corrected linear mixed effects model to account for donor bias. *Q<0.1; **Q<0.01; ***Q<0.001. Data from HD BM volumetric imaging. H) Mitochondrial morphometric features by subset. White dot indicates median. Q-values calculated using pairwise FDR-corrected linear mixed effects model to account for donor bias. *Q<0.1; **Q<0.01; ***Q<0.001. Data from HD BM volumetric imaging.

ASCp subsets were enriched for oxidative phosphorylation and translation gene sets (**Fig. 2F**), so we wanted to establish if these differences manifested in the morphology and organization of mitochondria and endoplasmic reticulum (ER). We sorted ASC subsets and TBCs from HD BM samples (**Fig. S8A-B, Table S1-2**), stained cells with Mitotracker Red CMX Ros, which stains active mitochondria in a membrane potential-dependent manner, and an antibody against protein disulfide isomerase (PDI), which is ubiquitously expressed in ER and responsible for disulfide bond formation, and performed volumetric fluorescence imaging (**Fig. 3D**). As expected, ASCs had substantially higher quantities of mitochondria and ER than TBCs, and differences in ER quantity and mitochondrial morphology were evident between subsets (**Fig. 3E**). Mitochondria can fuse and form networks that enhance their metabolic efficiency and resilience^47^, so we devised a network score to quantify the extent of a cell’s mitochondrial networking (**Fig. 3F**). We also generated other organelle morphometrics to quantify ER and mitochondrial morphological features, focusing on eleven, largely orthogonal morphometric features (**Fig. S8F**). We used these features to quantify mean Euclidean distance between subsets and found the ASCp subsets were more similar to each other than ASCr subsets, and likewise ASCr subsets were more similar to other ASCr subsets than ASCp subsets, mirroring the prior multi-omic findings (**Fig. S8G, 2A**). ASCp subsets had significantly higher total ER volume, suggesting a greater potential for antibody secretion (**Fig. 3G, S8H**). CD19+9^lo^ASCr and CD19+9+ASCr had greater total mitochondrial volume and more networked mitochondria than other subsets (**Fig. 3G-H**). All three ASCr subsets had significantly higher mean mitochondrial intensity than ASCp subsets, particularly CD19-ASCr, despite having less networked mitochondria. We validated these results using flow cytometry with two additional BM HDs and confirmed that unstimulated ASCr subsets exhibited higher mean Mitotracker Red intensity than ASCp subsets (**Fig. S8I-J**). In summary, we found that that DR+ASCp had the largest cell area and total ER volume, while ASCr subsets were enriched for enhanced mitochondrial features.

### DR+ASCp secrete large quantities of antibody and are responsive to stimulation

Having observed high gene set scores for translation and increased ER volumes in ASCp subsets, we asked if these features enabled increased antibody output. We screened Ig secretion from total BM ASCs across thirty culture conditions and selected two conditions for downstream assays: a stimulatory cocktail consisting of BAFF, CpG, IL-2, and IL-21 and a minimalist culture containing only BAFF (**Fig. S9A**). We sorted ASC subsets from HD BM samples (**Fig. S8A-B, Table S1-2**), cultured cells for 72 hours, and quantified the concentration of IgM, IgG, and IgA in culture supernatants (**Fig. 4A**). In parallel, we performed flow cytometry to quantify the frequency of IgH isotype usage for each subset and donor, enabling quantification of each well’s isotypic composition (**Fig. S9B**). No significant differences were observed between subsets in live cell frequencies after culture suggesting neither cell death nor proliferation account for difference in observed Ig secretion (**Fig. S9C**).

**Figure 4:**
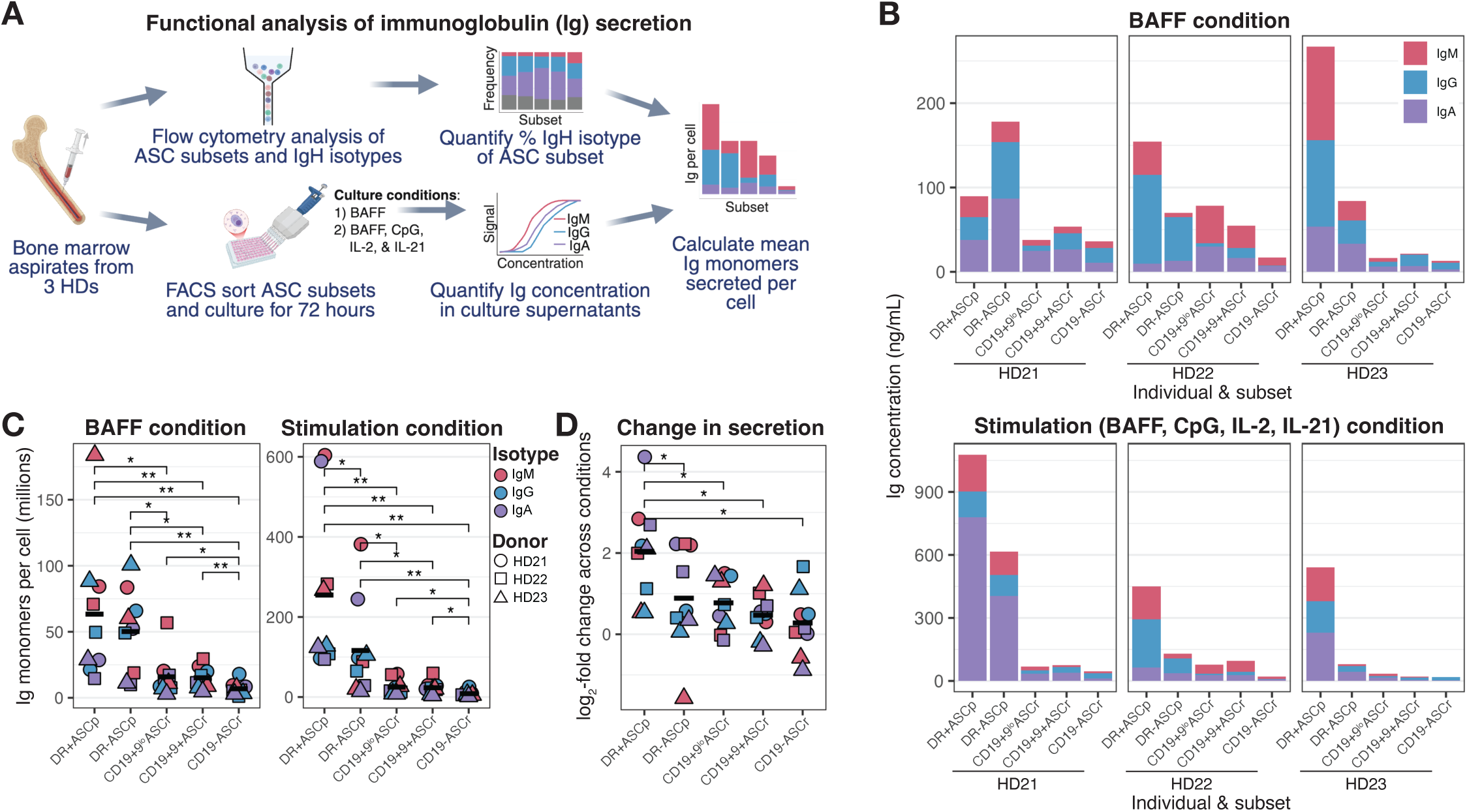
DR+ASCp secrete large quantities of antibody and are responsive to stimulation. A) Experimental diagram. B) Ig concentration from culture supernatants by isotype (color), subset (x-axis), individual (facet), and culture condition (panel). Data from HD BM MSD isotyping panel. C) Mean Ig monomers secreted per cell from culture supernatants by isotype (color), subset (x-axis), individual (shape), and culture condition (panel). Crossbar indicates mean. Q-values calculated by Wilcoxon signed rank test with FDR correction. *Q<0.1; **Q<0.01. Data from HD BM MSD isotyping panel. D) Log_2_-fold change in mean Ig monomers secreted per cell from culture supernatants from BAFF condition to stimulation condition by isotype (color), subset (x-axis), and donor (shape). Crossbar indicates mean. Q-values calculated by Wilcoxon signed rank test with FDR correction. Data from HD BM MSD isotyping panel. *Q<0.1.

In both culture conditions, DR+ASCp and DR-ASCp culture supernatants contained almost uniformly higher Ig concentrations compared to ASCr subsets (**Fig. 4B**). To account for baseline differences in the IgH isotypic composition of wells, we calculated the mean Ig monomers secreted per cell over the 72-hour culture for each IgH isotype, subset, and donor (**Fig. 4C**). In the BAFF culture, DR+ASCp and DR-ASCp had significantly higher levels of antibody secretion than all ASCr subsets, and CD19-ASCr had significantly lower levels of secretion than all other subsets. The same result was replicated in the stimulatory condition and additionally, DR+ASCp secretion levels were significantly higher than DR-ASCp. We quantified the log_2_-fold change in antibody secretion between the BAFF and stimulatory cultures and found DR+ASCp increased antibody secretion to a significantly greater degree than all other subsets (**Fig. 4D**).

We replicated these analyses comparing between IgH isotypes and found that IgM ASCs secreted significantly more antibody than IgG and IgA ASCs in both culture conditions, and IgG ASCs secreted significantly more antibody than IgA ASCs in the BAFF culture (**Fig. S9D**). There were no significant differences between IgH isotypes in log_2_-fold change between culture conditions (**Fig. S9E**). Given the unique responsiveness of DR+ASCp to the stimulatory culture, we assessed if the gene expression levels of the relevant receptors differed between subsets at baseline. We found DR+ASCp expressed significantly higher levels of *IFI16*, *IL2RB*, *LRRFIP1*, and *TNFRSF17*, which encodes BCMA (**Fig. S9F**). No significant differences were observed between IgH isotypes. In summary, we found DR+ASCp and DR-ASCp secreted high quantities of Ig and DR+ASCp more strongly upregulated Ig secretion in response to stimulation.

### Cytokine expression is largely invariant across ASC subsets

To assess if subsets displayed differential cytokine expression, we sorted subsets from HD BM samples (**Fig. S8A-B, Table S1-2**), cultured cells for 72 hours in BAFF or stimulation media and quantified supernatant cytokine expression. In the BAFF culture, only TNF-α, IL-10, and IL-2 were detected in culture supernatants, though IL-10 was detected in only one HD’s DR-ASCp (**Fig. S10A**). IL-2 concentrations were higher than TNF-α concentrations in culture supernatants, but no differences were observed between subsets, though the experiment was underpowered for statistical comparison (**Fig. S10B**). All analytes were detected in the stimulatory culture, though IL-2 was removed from the analysis since it was part of the stimulatory cocktail (**Fig. S10A**). Variance between HDs was low but no substantial differences were observed in cytokine expression between subsets, with the exception of IL-1β, which was noticeably higher in DR-ASCp from all three HDs (**Fig. S10C**). Some significant differences were observed in baseline gene expression levels of cytokines (gene list from Carrasco Pro *et al.*^48^), including significantly higher expression of *IL12A* in DR+ASCp, but this did not translate to increased IL-12 protein concentration after culture (**Fig. S10D**). *TNFSF13* which encodes APRIL, an important ASC survival factor, was significantly enriched in CD19-ASCr. In summary, cytokine expression was largely invariant across ASC subsets.

### ASCp are prevalent in peripheral blood and bone marrow ASCs preferentially colocalize with other ASCs of the same subset

ASCs differentiate in induction sites throughout the body and traverse blood to traffic to BM^13^. To evaluate how the composition of ASCs in PB compares to BM, we analyzed a previously generated fixed CITEseq dataset run on matched PB and BM samples from four HDs (**Fig. 5A**). We labeled subsets and found both ASCp subsets were significantly enriched in PB samples compared to BM samples (**Fig. 5B-C, S11A-B**). We observed a small fraction of ASCr subsets in PB samples, and these cells downregulated lineage-defining ASCp markers such as CD11a, but did not upregulate lineage-defining ASCr markers, such as CD31 (**Fig. S11C**). PB ASCr frequencies were also lower by manual gating (**Fig. S11D**), so these cells might not represent bona fide ASCr subsets. The subisotype composition of ASCs varied between tissues for each donor (**Fig. 5D**), with significantly more IgM and less IgG1 in blood ASCs (**Fig. S11E**). Interestingly, there were IgD ASCs amongst donors and these cells expressed surface IgD, but not IgM, as we previously observed (**Fig. S11F, S5H**).

**Figure 5:**
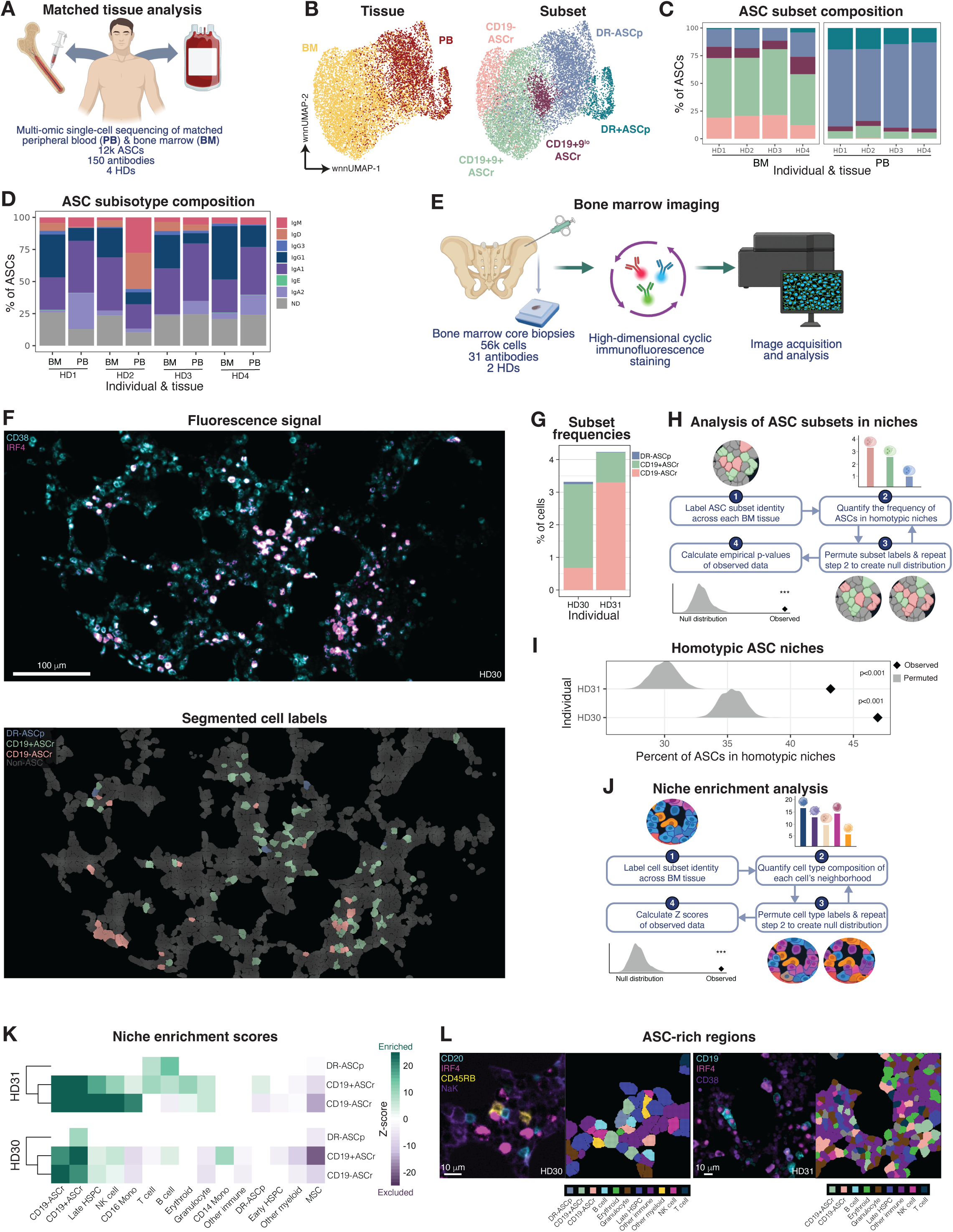
ASCp are prevalent in peripheral blood and bone marrow ASCs preferentially colocalize with other ASCs of the same subset. A) Experimental diagram. B) wnnUMAPs of ASCs colored by tissue (left) or subset (right). Data from HD matched fixed CITEseq. C) Percent subset of total ASCs by sample. Data from HD matched fixed CITEseq. D) Percent subisotype usage by sample. Data from HD matched fixed CITEseq. E) Experimental diagram. F) Representative region of interest from a bone marrow core colored by fluorescence signal (top) or subset labels of segmented cells (bottom). Data from HD BM spatial proteomics. G) Percent ASC subset of total cells by individual. Data from HD BM spatial proteomics. H) Computational diagram. I) Observed (black diamond) and randomly permuted (grey density) percent of ASCs found in homotypic niches (defined as all ASCs in a niche having identical subset identities), by individual. Empirical p-values calculated as the number of permutations with homotypic niche frequencies greater or equal to the observed homotypic niche frequency, divided by the number of permutations plus one. Data from HD BM spatial proteomics. J) Computational diagram. K) Scaled enrichment scores of cell types (x-axis) in ASC subset (y-axis) niches by individual (panel). Q-values<0.1 were colored white. Data from HD BM spatial proteomics. L) Representative images (panels) of ASC-rich regions colored by fluorescence signal (left) or cell type labels of segmented cells (right). Data from HD BM spatial proteomics.

To evaluate spatial patterning and ASC niche composition, we applied spatial proteomics by cyclic immunofluorescence to two normal BM tissue cores (**Fig. 5E, Table S1-2**). We segmented images, clustered cells, and annotated clusters with cell type labels based off their high-dimensional profiles (**Fig. 5F, S12A-B**). We identified DR-ASCp, CD19+ASCr, and CD19-ASCr subsets in BM cores, but were unable to distinguish CD19+9^lo^ASCr and CD19+9+ASCr subsets due to indiscernible CD9 staining. The frequencies of DR-ASCp in BM cores, which are not typically contaminated with blood^49^, were substantially lower than what we observed in BMAs (**Fig. 5G**), indicating blood contamination is a significant, but not exclusive, source of DR-ASCp in BMAs. We did not observe any DR+ASCp, likely due to insufficient cell numbers for detection of this rarer subset within BM cores.

ASC niches in the BM provide extrinsic signals that may impart longevity on ASCs and influence their cellular phenotypes^13^. If true, ASCs should be spatially organized into homotypic niches consisting of a single ASC subset, due to the influence of local factors. To test this hypothesis, we quantified the frequency of ASCs in homotypic, rather than mixed subset niches (**Fig. 5H**). Then, we then randomly permuted cells’ subset labels 1,000 times and recalculated the frequency of ASCs in homotypic niches for each permutation to generate a null distribution. In both HDs, ASCs were significantly more likely to be found in homotypic niches within BM than expected by chance (**Fig. 5I**).

To determine which cell types were enriched in ASC niches, we performed a similar permutation analysis (**Fig. 5J**). ASCr niches were significantly enriched for other ASCr, late hematopoietic stem and progenitor cells (late HSPCs; defined as CD34-CD38+), and NK cells, while mesenchymal stromal cells (MSCs) were significantly excluded (**Fig. 5K-L**). DR-ASCp niches did not have statistically significant enrichments shared between HDs, likely due to low cell counts. In summary, ASCp are prevalent in, but not exclusive to, PB, and BM ASCs preferentially colocalize with other ASCs of the same subset.

### All ASCr subsets and DR+ASCp can harbor long-lived immune memory

There has been controversy in the field concerning the phenotype of long-lived human ASCs^16–18^. To identify which subsets harbor long-lived immune memory, we collected BMAs from older healthy adults that received childhood measles and/or smallpox vaccination, sorted ASC subsets and TBCs, cultured cells for 72 hours in a stimulatory cocktail, and quantified antigen-specific IgG from culture supernatants (**Fig. 6A, S8A-B; Table S1-2**). Measles virus was in low public circulation during sample collection and smallpox was declared globally eradicated in 1980, hence individuals would not have recently been exposed to these viral antigens. Across five HDs, we identified measles-specific ASCs in all three ASCr subsets, but no measles specificity was observed in ASCp subsets, nor in negative control TBC wells (**Fig. 6B, S13A**). Likewise, amongst four HDs, all three ASCr subsets harbored vaccinia virus (VACV) specificity against several vaccine antigens (**Fig. 6C, S13B-C**). No VACV specificity was observed in seronegative individuals (**Fig. S13B**).

**Figure 6:**
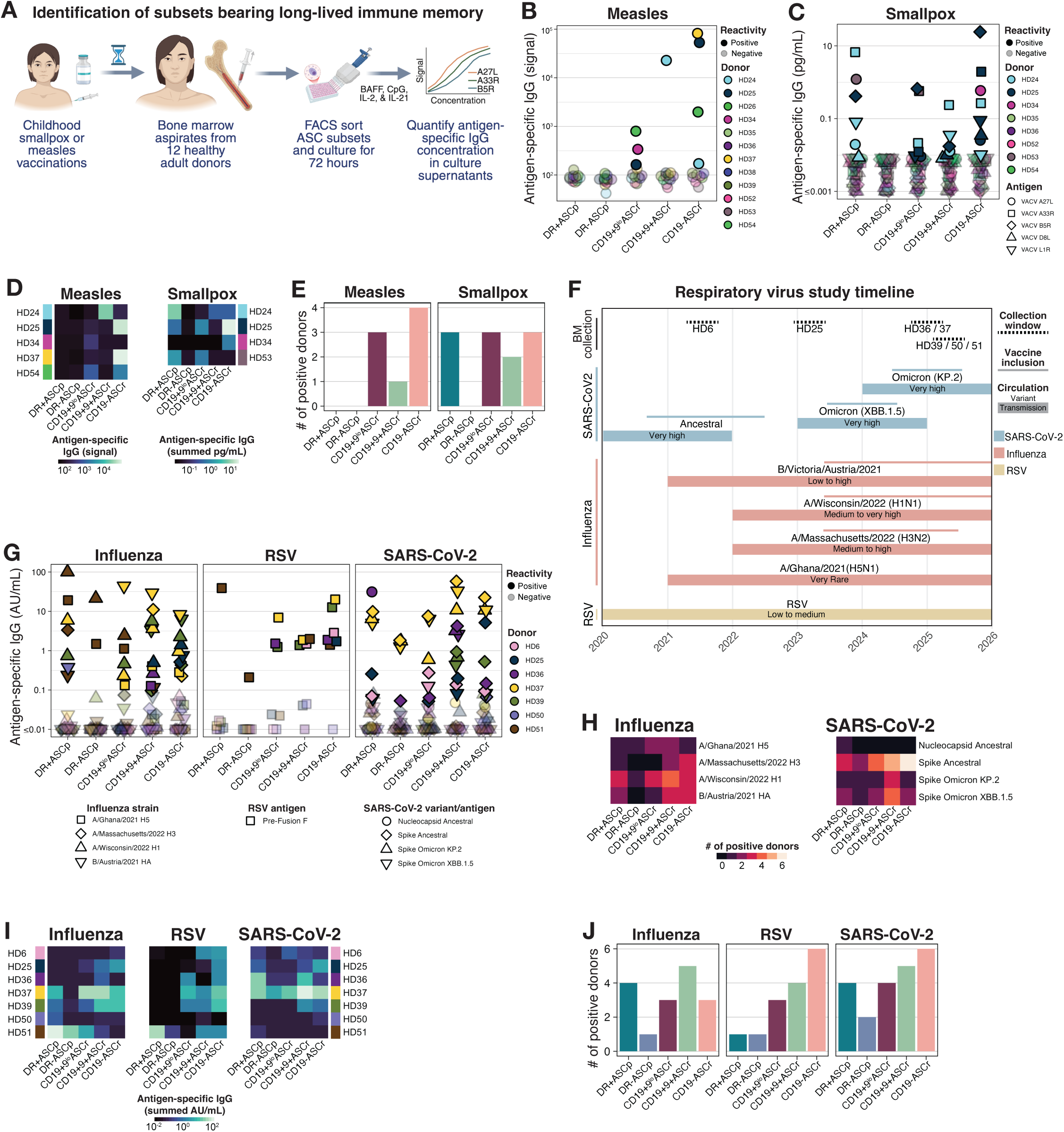
All ASCr subsets and DR+ASCp can harbor long-lived immune memory. A) Experimental diagram. B) Antigen-specific IgG signal from culture supernatants by individual (color) and subset (x-axis). Transparent dots indicate values below positivity threshold. Data from HD BM MSD measles panel. C) Antigen-specific IgG concentration from culture supernatants of non-seronegative donors by vaccinia antigen (shape), individual (color), and subset (x-axis). Transparent dots indicate values below positivity threshold. Data from HD BM MSD orthopoxvirus panel. D) Measles-specific IgG signal (left) and summed VCV-specific IgG concentration (right) from culture supernatants by individual and subset. Data from HD BM MSD measles and orthopoxvirus panels. E) Number of HDs with culture supernatants positive for specificity against any antigen by subset (x-axis) and virus (panels). Data from HD BM MSD measles and orthopoxvirus panels. F) Diagram of study timepoints displaying BM collection ranges (exact timepoints withheld to protect HD privacy), viral/strain/variant circulation in the United States, and viral/strain/variant inclusion in vaccine formulations in the United States. G) Antigen-specific IgG concentration from culture supernatants by individual (color), strain/variant/antigen (shape), subset (x-axis), and virus (panels). Transparent dots indicate values below positivity threshold. Data from HD BM Respiratory panel 7. H) Number of HDs with culture supernatants positive for specificity against viral strains/variants/antigens (y-axis) by subset (x-axis) and virus (panels). Data from HD BM Respiratory panel 7. I) Antigen-specific IgG concentrations from culture supernatants by individual (y-axis) and subset (x-axis), summed across strain/variant/antigen for each virus (panels). Data from HD BM Respiratory panel 7. J) Number of HDs with culture supernatants positive for specificity against any antigen by subset (x-axis) and virus (panels). Data from HD BM Respiratory panel 7.

Unexpectedly, VACV specificity was observed in DR+ASCp in three HDs, against multiple antigens (**Fig. 6C**). DR+ASCp secreted the highest overall quantities of VACV-specific antibodies compared to ASCr subsets in two HDs (**Fig. 6D**). These results establish that all three ASCr subsets (comprised of both CD19+ and CD19-ASCs) are capable of harboring long-lived immune memory (**Fig. 6E**). Further, these results suggest there is a mechanism by which presumably recently generated DR+ASCp can differentiate in the absence of recent antigen exposure.

To contextualize these results, we interrogated the composition of ASC subsets specific to circulating respiratory viruses: respiratory syncytial virus (RSV), four influenza strains (three flu A and one flu B), and three severe acute respiratory syndrome coronavirus 2 (SARS-CoV-2) variants. We applied the same experimental pipeline to HD BM samples collected at indicated timepoints in which viruses were circulating and/or included in vaccine formulations (**Fig. 6F, Table S2**). All five ASC subsets harbored specificity to each of the three viruses in at least one HD (**Fig. 6G, S13D**). Among the influenza strains, most donors held specificity to A/Wisconsin/2022 HA (**Fig. 6H**), which was in high circulation and included in seasonal influenza vaccine formulations^50^. Among SARS-CoV-2 variants, specificity to ancestral spike protein was held by the highest number of HDs for all ASC subsets. Further, six of seven HDs held specificity to ancestral spike in CD19-ASCr. The composition of ASC subsets holding specificity to circulating viral antigens included ASCp subsets but was more enriched for ASCr subsets, with donor variability likely stemming from distinct infection and vaccination histories (**Fig. 6I-J**). Overall, there were not substantial differences in the composition of subsets holding specificities to recently circulating antigens (**Fig. 6J**) compared to non-circulating antigens (**Fig. 6E**). In summary, we found all ASCr subsets harbored long-lived immune memory to childhood vaccine antigens and smallpox vaccine specificity was also maintained in DR+ASCp.

### Non-malignant ASCr frequencies are diminished at least two years after bone marrow transplant in patients with multiple myeloma

To evaluate the utility of our ASC subset paradigm for translational studies, we analyzed CITEseq data from longitudinal BM samples from patients newly diagnosed (ND) with MM, an ASC malignancy presenting in BM (**Fig. 7A, Table S1, S8**). We evaluated ∼3 years of samples starting from pre-treatment (PRETX), through the end of induction therapy (EIND), and post-autologous stem cell transplant at 90 days (TRN90), 1 year (TRNY1), and 2 years (TRNY2). At PRETX, most patients exhibited serum Ig titers above the healthy reference for the isotype of their malignant cells and below the healthy reference for all other isotypes (**Fig. 7B**). At post-treatment timepoints, no patients’ serum titers were above the healthy reference, but the majority of patients had serum antibody titers below the healthy reference for all isotypes, indicative of ongoing humoral dysfunction. Likewise, the total frequency of BM ASCs was significantly higher at PRETX timepoints compared to HDs due to the overabundance of malignant ASCs but dropped after treatment and was significantly lower than HDs at post-transplant timepoints (**Fig. 7C**). We labeled malignant and non-malignant ASCs in a patient-specific manner based on differences in heavy and light chain isotype usage, and transcriptomic and proteomic features (**Fig. S14A**). As expected, the frequency of non-malignant ASCs out of total ASCs increased through therapy as malignant cells were eliminated, which coincided with a recovery of diverse subisotype usage in the BM ASC compartment (**Fig.7D, S14B**).

**Figure 7:**
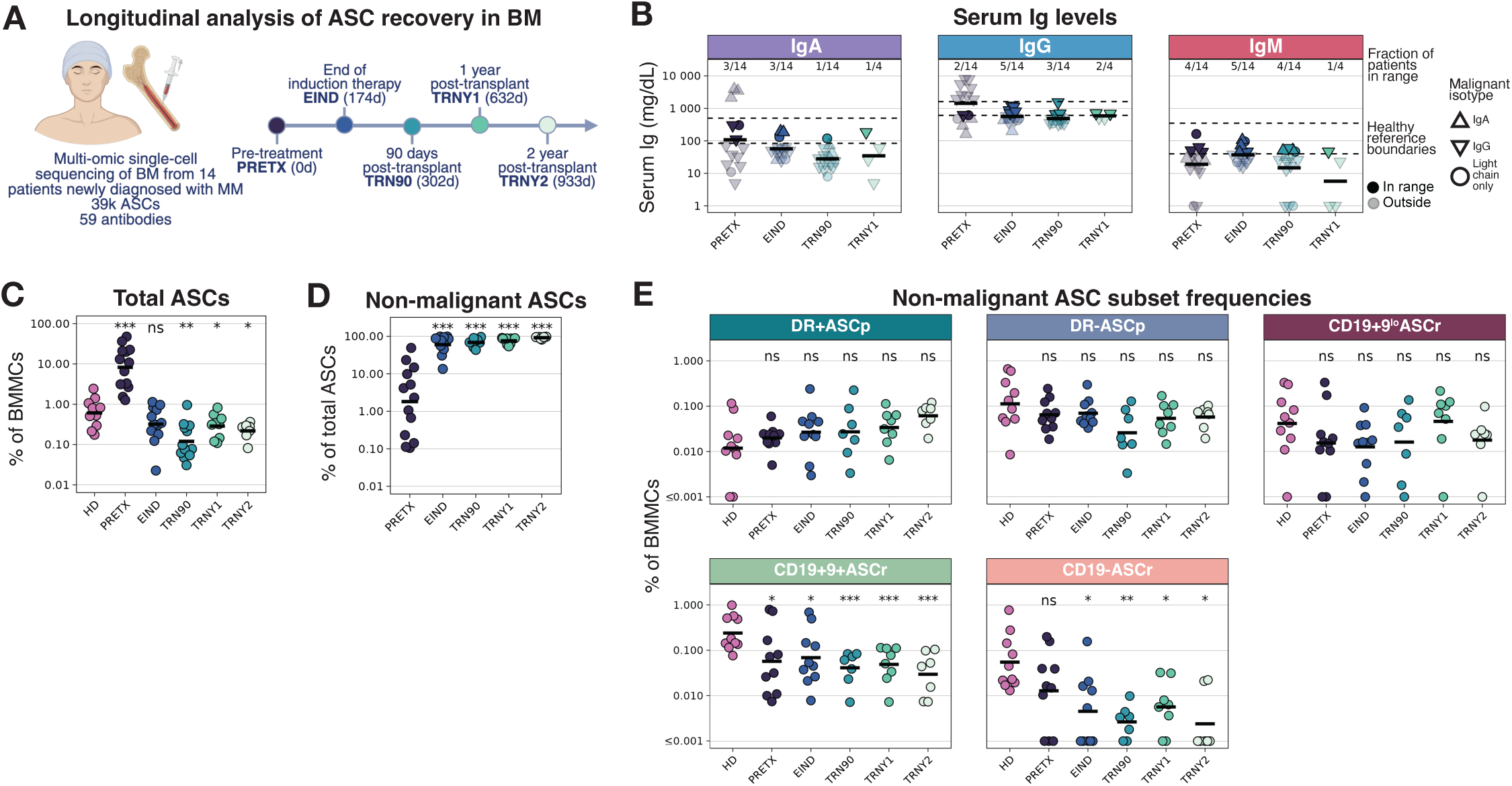
Non-malignant ASCr frequencies are diminished at least two years after bone marrow transplant in patients with multiple myeloma. A) Experimental diagram. B) Serum Ig concentrations of patients with NDMM by timepoint (x-axis), isotype (y-axis), and the isotype of each patient’s malignant cell (shapes). Transparency indicates values outside the healthy reference ranges specified in the clinical reports. Data from clinical quantitative Ig assay. C) Percent ASCs of total BMMCs for HDs (left) and patients with NDMM, by timepoint (x-axis). Crossbar indicates mean. Q-values calculated by Wilcoxon rank sum test with FDR correction, comparing to HDs. Data from MM longitudinal BM CITEseq. *Q<0.1; **Q<0.01; ***Q<0.001 D) Percent non-malignant ASCs of total ASCs for patients with NDMM, by timepoint (x-axis). Crossbar indicates mean. Q-values calculated by Wilcoxon rank sum test with FDR correction, comparing to PRETX. Data from MM longitudinal BM CITEseq. ***Q<0.001 E) Percent non-malignant ASC subset of total BMMCs for HDs (left) and patients with NDMM, by timepoint (x-axis) and subset (panels). Crossbar indicates mean. Q-values calculated by Wilcoxon rank sum test with FDR correction, comparing to HDs. Data from MM longitudinal BM CITEseq. *Q<0.1; **Q<0.01; ***Q<0.001

We labeled subset identities in non-malignant ASCs and compared their frequencies to HDs (**Fig. 7E**). There were no significant differences in the frequencies of DR+ASCp, DR-ASCp, nor CD19+9^lo^ASCr out of total BM mononuclear cells (BMMCs), but CD19+9+ASCr and CD19-ASCr were significantly less frequent than HDs at all post-treatment timepoints. The frequency of DR+ASCp out of total ASCs was significantly increased compared to HDs (**Fig. S14C)**. These findings suggest a compensatory increase in nascent ASC differentiation in these patients. However, ongoing dysregulation in the BM^51^ may impair ASC maturation into or retention of CD19+9+ASCr and CD19-ASCr phenotypes, depressing overall serum antibody titers.

We evaluated clinical features associated with non-malignant ASC recovery post-treatment and found positive correlations with the percent of eosinophils out of total white blood cells, overall serum protein concentrations, serum gamma globulin concentration, and serum IgM and IgA concentrations (**Fig. S14D**). Mean corpuscular volume in PB, an erythrocyte metric, and the concentration of PB monocytes were negatively correlated with non-malignant ASC frequencies. In summary, we found that up to two years after autologous stem cell transplant, patients in remission from MM have diminished frequencies of CD19+9+ASCr and CD19-ASCr in their BM and lower serum antibody titers.

### Malignant ASCs display aberrant expression profiles

To evaluate how malignant ASC profiles differ from non-malignant ASCs, we applied fixed CITEseq to PRETX BMA samples from patients with NDMM and relapsed/refractory MM (RRMM) and labeled malignant ASCs and non-malignant ASC subsets, as before (**Fig. S14A S15A, Table S1, S8**). We found dozens of DEPs and hundreds of DEGs between malignant ASCs and non-malignant subsets, with more differential features compared to ASCp subsets than ASCr subsets (**Fig. S15B, Table S9-10**). Malignant ASCs were mostly CD11a-, CD31+, HLA-DR-, CD19-, and CD9+/-, more closely resembling ASCr than ASCp subsets, though CD9-CD19-cells were rarely observed in non-malignant ASCs (**Fig. S15C**).

To identify potential therapeutic targets uniquely upregulated by malignant ASCs, we visualized surface protein expression by malignant ASCs (labeled with patient IDs), non-malignant ASC subsets, and other BM immune cells (**Fig. S15D**). BCMA was significantly upregulated compared to non-malignant ASC subsets, but the other myeloma therapeutics, CD38 and CD319, were not. CD47, CD117, ITGB7, and PD-L1 were all significantly upregulated in malignant cells and have existing therapeutic antibodies approved for other indications, indicating they may have efficacy in patients with MM (**Fig. S15D-E**). We also identified a number of DEGs that could be mined for therapeutic potential (**Fig. S15F**). In summary, malignant cells more closely resembled ASCr than ASCp and CD47, CD117, ITGB7, and PD-L1 represent potential therapeutic targets for MM.

## DISCUSSION

By integrating over 170 surface protein measurements from a bespoke CITEseq panel with transcriptomic data, we comprehensively profiled human BM ASC identity and discovered five distinct subsets. We report surface proteins that reliably distinguish ASC subsets in low dimension, enabling clinical immune monitoring and cell sorting for *in vitro* manipulation.

We introduced new terminology to differentiate immature ASCp subsets found in PB and BM from mature BM-resident ASCr subsets. Historically, immature ASCs have been called plasmablasts, but this term is misleading because only a fraction of these cells are cycling (blasting)^52^. Further, plasmablasts are described as short-lived^53^, but our results support a model in which ASCs exist along a maturation continuum from DR+ASCp through ASCr subsets, mirroring findings in murine models^10,54^. This implies all long-lived ASCs pass through a so-called short-lived plasmablast stage, so we abandoned this terminology and its associations in favor of a more accurate and modern nomenclature^55^.

Immature ASCp exhibited high translation gene sets scores, contained high volumes of ER, and secreted more antibodies than more mature ASCr. A recent study reported higher antibody secretion rates in mature ASC subsets^56^. These discordant results are likely explained by culture conditions: in that study, ASCs were cultured in a complex media and hypoxic conditions that attempt to mimic the BM microenvironment and induce *in vitro* ASC maturation^57^. These conditions likely favor ASCr subsets already adapted to the BM microenvironment, while the simpler culture conditions used in the current study enable more neutral comparisons. Others have also observed high antibody secretion rates by blood ASCs^58^, which makes intuitive sense as ASCp are generated in high quantity during an active immune response where high antibody titers are needed to neutralize active infectious agents. ASCr are optimized for long-term antibody maintenance rather than maximal short-term output, resembling ultramarathon runners rather than sprinters. Indeed, a key feature in the survival and maturation of ASCp into ASCr may be downregulation of antibody secretion rates, culling of ER, and the increases in mitochondrial volume and networking observed in ASCr. Mitochondrial morphometrics were discordant with the oxidative phosphorylation gene set enrichments observed in ASCp subsets. This discrepancy suggests that rather than being more metabolically fit, ASCp are upregulating this gene program to accommodate unmet metabolic demands, which highlights the importance of collecting functional or proteomic measurements for interpretation of metabolic states^59^.

ASCs preferentially co-localized with ASCs of the same subset identity in BM niches, indicating local niche signals may induce or reinforce ASCr phenotypes. This suggests ASCr subset identity does not solely reflect cellular age/longevity. Late HSPCs and NK cells were enriched in ASC niches, but further studies are warranted to define the key constituents of ASC niches in human tissues. Patients in remission from NDMM exhibited poor recovery of non-malignant CD19+9+ASCr and CD19-ASCr, suggesting these subsets may be particularly dependent on stable and supportive BM niches, which are not fully restored after treatment^51,60^. Incomplete recovery of these subsets persisted at least two years after transplant and may contribute to persistent deficits in humoral immunity following treatment^21^.

While CD19-ASCr can be long-lived, they do not necessarily represent a universal terminal differentiation state nor the optimal phenotype for longevity. We overlayed a long-lived ASC murine gene score^10^ on our data and found the transcriptional profile was highest in CD19+9+ ASCr, not CD19-ASCr. Further, our data agree with prior findings that all ASCr subsets can harbor long-lived pathogen-specific memory, regardless of CD19 expression^17–19^. Consequently, the absence of antigen-specific CD19-ASCr <2 years post-boost does not imply diminished durability of mRNA vaccine responses^61^. In our data, all ASC subsets, including CD19-ASCr, retained SARS-CoV-2 spike specificity, consistent with studies showing that SARS-CoV-2 infection induces long-lived ASCs in human BM^62^ and that mRNA vaccination establishes phenotypically diverse spike-specific ASC populations^63^.

Immature DR+ASCp exhibited smallpox vaccine specificity in multiple donors, so these cells must differentiate from another long-lived population in the absence of antigen, or must be themselves a stable, long-lived population. CD38++CD20-blood ASC frequencies significantly drop after anti-CD20 therapy, suggesting they differentiate from CD20+ B cells and are not a stable population^64^. This transience indicates DR+ASCp either differentiate into ASCr phenotypes or perish. Therefore, we postulate that VACV-specific DR+ASCp are recently differentiated from bystander-activated long-lived memory B cells in the absence of recent antigen exposure. Posited by Lanzavecchia and colleagues^65^, the bystander activation model is supported by the observation that numerous vaccine-responsive antibody clonotypes are not specific to vaccine components^66^. Further, there are substantial clonal connections between plasmablasts and memory B cells in the absence of recent antigen exposure in the blood of HDs^31,67^. We did not observe DR+ASCp specificity to measles, so further studies are required to establish and characterize bystander activation of memory B cells as a bona fide mechanism of humoral replenishment.

In addition to advancing our understanding of ASC biology, the current study provides low-dimensional gating strategies to quantify and prospectively isolate ASC subsets, enabling further functional characterization and immune monitoring in the context of vaccination, infection, cancer, and autoimmunity. Tracking these subsets in diverse immune contexts, particularly when integrated with antigen specificity, spatial localization, and clonal identity, may offer insights into the regulation of durable protective responses to vaccination and infection as well as pathogenic antibody production in diseases such as MM, rheumatoid arthritis, and systemic lupus erythematosus. These insights may reveal new opportunities for targeted induction or depletion in clinical settings.

## DATA & CODE AVAILABILITY

Data and code are available at: https://doi.org/10.5281/zenodo.19499360 Interactive visualization of HD BM fixed CITEseq data available at: https://apps.allenimmunology.org/aifi/resources/antibody-secreting-cells/

## DECLARATION OF INTERESTS

A.W.G is on the science advisory board of Foundry Biosciences. D.J.G. has received research funding, has served as an adviser, and has received royalties from Juno Therapeutics, a BMS company; has served as an adviser, and received research funding from Janssen Biotech and Seattle Genetics; has served as an adviser for GlaxoSmithKline, Celgene, Ensoma, and Legend Biotech; and has received research funding from SpringWorks Therapeutics, Sano, and Cellectar Biosciences. E.W.N. is a cofounder, adviser, and shareholder of ImmunoScape.

## FUNDING

This work was supported by the Allen Institute. D.R.G. was supported by an NIH-NIAID K99 Award: 1K99AI185149-01A1, a Cancer Research Institute Irvington Postdoctoral Fellowship, an American Society of Hematology Fellow Scholar Award, and a Fred Hutchinson Cancer Center Translational Data Science Pilot Award.

## AUTHOR CONTRIBUTIONS

D.R.G. conceptualization, formal analysis, investigation, writing of the original draft, and visualization. E.D. methodology, investigation, formal analysis, writing of the original draft, and visualization. H.Y. methodology, investigation, validation, and data curation. S.A.L. methodology, investigation, and writing (review and editing). A.S. and S.K. formal analysis, software, data curation, and visualization. J.M. data curation, validation, and methodology. A.C., C.G.P., and Z.T. formal analysis and data curation. S.N.K., M.A.D., M.W., and J.R. methodology, investigation, and validation. V.P. methodology, investigation, validation, and supervision. P.G. data curation and investigation. T.J.S. methodology and investigation. S.A.-S. methodology, resources, investigation, and supervision. P.J.W. methodology, investigation, and validation. M.-P.P. resources, software, and supervision. Z.H. data curation and supervision. K.E.H., V.H., and B.M. investigation. P.R. investigation and validation. S.M. methodology and software. U.K. data curation and validation. N.E. investigation. L.G. software and visualization. M.A-H. resources and supervision. C.E.G. writing (review and editing) and resources. M.K. project administration. X.L. writing (review and editing), data curation, and software. M.P.V. methodology, supervision, and software. T.F.B. supervision and funding acquisition. A.W.G. supervision, funding, and writing (review and editing). M.S. supervision, resources, and writing (review and editing). S.C.B. resources and funding acquisition. P.J.S. supervision and resources. D.J.G. resources. E.W.N. and T.R.T. supervision and writing (review and editing). M.C.G. conceptualization, formal analysis, investigation, writing of the original draft, visualization, and supervision.

## ACKNOWLEDGEMENTS

We thank A. Tsai for cytometry experiment contributions and E. Kuan and A. Savage for experiment advice. We also thank A. Ott, M. Ambrose, A. Gogate, and J. Harvey for computational and data infrastructure contributions. We thank L. Shiraiwa, K. Dang, E. Davis, D. Parrish, R. Messinger and Y. Aggoune for laboratory operations and software support. We thank S. Kaech, P. Meijer, E. Coffey, and L.A. Becker for organizational and resource support. We thank C. La France and G. Kong for assistance with data visualization apps. We thank X. Song and C. Crider for patient data administration. We thank K.B. Glass for moral support. Figures were created using BioRender (http://www.biorender.com) and Illustrator (Adobe).

## METHODS

### EXPERIMENTAL AND SUBJECT DETAILS

#### HD cohort information

Deidentified human PB (n=4) and BM (n=39) were obtained from healthy adult donors (StemCell Technologies, AllCells/Discovery Life Sciences). HDs provided informed consent before participation and these studies were approved by the Allen Institute or Stanford University Institutional Review Board (IRB). Deidentified bone marrow core tissue samples were collected from 2 adult patients undergoing core biopsy sampling for screening purposes (Accio Human Biobank). Clinical pathology assessment of the biopsy samples, consisting of tissue staining, flow cytometry, and fluorescence in situ hybridization, indicated normocellular bone marrow with trilineage hematopoiesis. The samples were negative for granulomas, lymphoid aggregates, malignancy, monoclonality, increased blasts and abnormal cytogenetics. Basic demographics for all normal tissue and cell sample donors are provided (**Table S2**).

#### Patients with MM cohort information

A cohort of 24 patients aged 30 to 70 with NDMM and 6 patients aged 64 to 74 with RRMM was enrolled in Seattle, WA (IRB 10265). This study was approved by the IRB of Fred Hutchinson Cancer Center. Amongst patients with RRMM, only PRETX BM samples were assayed. Amongst patients with NDMM, BMAs and PB samples were collected prior to receiving induction therapy (PRETX), post-induction therapy (EIND), and at multiple post-transplant timepoints: 90 days, 1 year, and 2 years (TRN90, TRNY1, TRNY2). Clinical tests performed on patient PB samples at post-treatment timepoints were used for correlative analyses (**Fig. S14D**). Patients with NDMM received induction therapy (consisting of 2- or 3-drug combinations of Lenalidomide, Bortezomib, Dexamethasone, Carfilzomib, Pomalidomide, or Cyclophosphamide), stem cell mobilization, melphalan, autologous stem cell transplant, and maintenance therapy (consisting of 1- to 4-drug combinations (of Lenalidomide, Bortezomib, Dexamethasone, Carfilzomib, Pomalidomide, or Isatuximab). A subset of patients with NDMM in this cohort were previously included in another study^68^. Patient characteristics are reported (**Table S8**).

#### HD sample collection

BMAs were collected from the posterior iliac crest of healthy IRB consenting adult donors via needle aspiration into heparin anticoagulant (AllCells/Discovery Life Sciences; Stemcell Technologies). Either a 25mL or 50mL aspirate was drawn from each hip from a maximum of 2 sites using multiple syringes containing anticoagulants. The aspirate was filtered to remove clots, bone chips, spicules, and pooled across donor draws. Fresh BMAs were then shipped overnight on 4°C gel packs (AllCells/Discovery Life Sciences). Otherwise, BMAs were processed for BMMCs over density gradient and cryopreserved in Cryostore10 (Stemcell Technologies) before being shipped on dry ice and stored in the gaseous phase of liquid nitrogen until use. BM interstitial fluid samples were collected from the cell-free supernatant of fresh BMA samples using centrifugation and then immediately aliquoted and frozen at -80°C. PB samples were collected from healthy IRB consenting adult donors through leukophoresis, in which approximately 2-3x blood volumes were processed through the Spectra Optia Apheresis System with acid-citrate-dextrose solution A anticoagulant (Stemcell Technologies). Leukopak blood samples were processed for PB mononuclear cells (PBMCs) over density gradient, cryopreserved in Cryostore10, and stored in the gaseous phase of liquid nitrogen until use. PB serum or plasma supernatant samples were isolated using centrifugation, aliquoted, and frozen at -80°C.

#### MM cohort sample collection

Fresh whole blood samples were collected from these subjects in heparinized syringes or sodium heparin vacutainers and processed to PBMCs through by density gradient. PBMCs were aliquoted in vials of 5x10^6^ cells/ml each and frozen in cryo-storage solution (90% FBS and 10% Dimethyl sulfoxide (DMSO)) within 4 hours of blood draw using a pre-chilled 4°C CoolCell or Mr. Frosty freezing container stored at -80°C for up to 72 hours before being transferred to liquid nitrogen for long-term storage. Fresh whole blood samples were collected from NDMM subjects in K2 EDTA vacutainers or tubes and plasma supernatant samples were isolated using centrifugation, aliquoted, and frozen at -80°C within 4 hours of blood draw. BMAs were collected from patients with NDMM in heparinized syringes/sodium heparin vacutainers and BMMCs were isolated by density gradient. BMMCs were aliquoted in vials each containing 1x10^6^ cells and frozen in cryo-storage solution using a pre-chilled 4°C CoolCell or Mr. Frosty freezing container stored at -80°C for up to 72 hours before being transferred to liquid nitrogen for long-term storage.

#### Custom virus serology control samples

Cryopreserved plasma samples were collected from measles virus (MeV)-specific Ig seropositive and seronegative adults (Virion-Serion), specifically MeV-specific IgG-positive, IgG-negative and IgM-positive donors. Sample serology status was determined by CE-certified measles virus IgG and IgM SERION ELISA tests (Serion Diagnostics) and the LIAISON Rubeola IgG Assay (Diasorin). MeV-specific IgG-positive and IgG-negative plasma samples were diluted in a range from 1:250 to 1:8x10^6^ and analyzed within a custom MeV HA antigen assay (MSD) to determine positivity thresholds. For the commercially available viral antigen-specific IgG assay kits (smallpox/orthopoxvirus and respiratory viruses; MSD), an antibody calibrator series or reference standard and multiple serology controls were provided with the kit, as described further in *ASC culture supernatant processing and data acquisition*.

#### Cryopreserved healthy BMMC and PBMC sample processing

Cryopreserved PBMCs or BMMCs were rapidly thawed at 37°C and diluted into Thawing Media Solution, consisting of RPMI-1640 media with ATCC modification (Gibco) supplemented with 20% heat inactivated fetal bovine serum (hi-FBS) (Sigma) and 1% Penicillin-Streptomycin (Gibco), and then washed twice in the media solution at 4°C. Washed BMMCs were resuspended in the media solution, filtered twice using 100uM pore size, and stored at 4°C until further processing.

#### Fresh BMA sample processing

Upon arrival, fresh and 4°C chilled BMAs (AllCells/Discovery Life Sciences) were immediately processed for BMMCs over density gradient. Total BMAs were first diluted 1:1 with Dulbecco’s Phosphate Buffered Saline (DPBS), supplemented with 2% hi-FBS, prior to density gradient separation for BMMC collection. The isolated BMMCs were then resuspended in BM Wash Buffer (DPBS containing 1% hi-FBS and 0.01% of 10000x Benzonase (Sigma-Aldrich)) and filtered twice using 100uM pore size. BMMCs were then immediately processed for cell enrichment and/or subsequent assays.

#### Total B cell enrichment

Washed and filtered BMMCs were incubated with Human TruStain FcX Blocking Buffer (BioLegend) following manufacturer’s instructions. BMMCs were subsequently enriched for total B cells by negative selection with the Human Pan-B Cell Enrichment Kit (Stemcell Technologies) following manufacturer’s instructions. A cell aliquot was stained with acridine orange/propidium iodide (AO/PI) viability dye and counted with the Nexcelom Cellaca MX cell counter. For mass cytometry assays, PBMC and BMMCs underwent magnetic lineage depletion using BD Streptavidin Particles Plus and the BD IMag Cell Separation Magnet (BD Biosciences) with a cocktail of biotinylated antibodies consisting of CD3, CD7, CD15, CD33, CD56, CD61, and CD235ab (BioLegend), according to the manufacturer’s instructions. Enriched total B cells were stored in media solution at 4°C until further processing.

#### Total ASC enrichment

Washed and filtered BMMCs were resuspended in 400uL Wash Buffer (Miltenyi Biotec) per 10⁸ cells. The Plasma Cell Isolation Kit II (Miltenyi Biotec) was then used to enrich total ASCs from BMMC samples following manufacturer’s instructions. A cell aliquot was stained using AO/PI viability stain and counted with the Nexcelom Cellaca MX cell counter. Isolated ASCs were then immediately taken into subsequent assays.

#### HD BM ASC subset fluorescence-activated cell sorting (FACS)

Enriched total B cells or ASCs from healthy donor BMMC samples were stained with Fixable Viability Stain 510 (BD Biosciences) and a panel of fluorescently-tagged antibodies to identify ASC subsets (**Table S1**). Cells were stained for 30 minutes at 4°C protected from light, washed twice with cold Wash Buffer (DPBS with 2% hi-FBS), and filtered through a 70um mesh prior to being sorted by FACS on a 4-laser BD FACS Aria Fusion using a 100um nozzle. A sequential gating strategy was used to exclude debris, doublets, dead cells, and lineage-positive cells (**Fig. S8A)**. TBCs were defined as CD10^+^CD38^+^ and total ASCs as CD10^-^CD38^h^. Total ASCs were divided into ASCp, defined as CD31^-^CD11a^+^, and ASCr, defined as CD31^+^CD11a^-^. ASCp subsets were then sorted as HLA-DR+ and HLA-DR-, and ASCr subsets were sorted as CD9^lo^CD19+, CD9+CD19+ and CD9+CD19-(**Fig. S8B**). ASC subsets were collected in sterile 5mL polypropylene tubes containing 1mL cold, pure hi-FBS to preserve viability. Sorted subsets were stored at 4°C and immediately transferred for subsequent culture and analysis.

#### HD BM ASC cultures

Total ASCs or sorted ASC subsets were washed once with media and resuspended in culture media, consisting of RPMI-1640 media (ATCC modification; Gibco) with 20% hi-FBS (Sigma) and 1% Penicillin-Streptomycin (Gibco). Cells were cultured with BAFF (1ug/mL, PeproTech) or a stimulation cocktail containing BAFF (1ug/mL, PeproTech), CpG ODN 2006 (10ug/mL, Invivogen), IL-21 (0.1ug/mL, PeproTech) and IL-2 (0.1ug/mL, Biolegend) in non-treated 96 well U-bottom plates. Cultures were maintained at 37°C and 5% CO₂ for 72 hours. After culturing, cells were immediately pelleted and cell-free supernatants were then collected. For cell counting and viability experiments (n=2 BMA samples), an aliquot of cells from each sorted ASC subset and culture replicate were stained with AO/PI viability dye and counted on a Denovix CellDrop FLi automated cell counter with a 400µm chamber height after 72 hours in culture. For the ASC culture condition screen, total BM ASCs were stimulated for 72 hours with CpG ODN 2006 (10μg/mL, Invivogen) or BCR crosslinker (anti-kappa light chain; 1μg/mL, Southern Biotech) with various combinations of stimulatory reagents, including MEGACD40L (0.1μg/ml, Enzo Life Sciences), BAFF (1μg/mL, PeproTech), CXCL12 (0.1μg/ml, R&D Systems), IL-2 (0.1μg/mL, Biolegend), and IL-21 (0.1μg/mL, PeproTech). In the screen, each culture well was seeded with 3600 cells, while in the downstream experiments, each culture well was seeded with 2000 cells. After culturing, cells were immediately pelleted, and cell-free supernatants were collected. Supernatants were immediately analyzed with the Isotyping Panel 1 Human/NHP Kit (Mesoscale Discovery) per manufacturer’s recommended protocol to quantify secreted antibodies (IgG/A/M).

#### HD BMMC and PBMC CITEseq pipeline

Donor-matched PBMC and BMMC samples were directly taken into sequencing workflows for CITEseq analysis. All other HD BMMC samples were enriched for total B cells, centrifuged, and resuspended in 1X Mojosort Isolation Buffer (Biolegend). To remove dead cells and debris, B cells were processed with the Mojosort Human Dead Cell Removal Kit (Biolegend) following manufacturer’s instructions and resuspended in wash buffer (0.22µm pore-filtered DPBS supplemented with 1% hi-FBS). Cell samples were stained with TotalSeq-C Universal Human cocktail (Biolegend, 399905) and a custom TotalSeq-C antibody cocktail (**Table S1**) and incubated at 4°C for 30 minutes, according to the ‘Cell Surface and Intracellular Protein Labeling for Chromium Fixed RNA Profiling’ protocol (10X Genomics, CG000529, Rev C).

Following staining, BMMC samples were washed once with wash buffer and then fixed according to a modified protocol of the ‘Fixation of Cells and Nuclei for Chromium Fixed RNA Profiling’ protocol (CG000478, Rev C), as previously described^69^. Briefly, samples were fixed in a final concentration of 4% PFA for 1 hour at 20°C in a total volume of 200uL of fixative per sample. After fixation, samples were centrifuged at 850xg for 5 minutes at room temperature (RT), and supernatant was removed using the Integra ViaFlow384. Sample wells were then quenched with 200uL Quenching Buffer at 4°C following the 10X Genomics guidelines (CG000478, Rev C) and centrifuged at 850xg for 5 minutes at 4°C. Supernatant was removed and sample was resuspended in 0.4% BSA (Sigma-Aldrich) in DPBS as an additional wash step. Up to 10^6^ fixed cells per sample were barcoded for multiplexing using the Fixed RNA Feature Barcode Multiplexing Kit (10X Genomics, PN 1000628) with oligo-tagged antibodies (HTOs). Probe hybridization was completed according to the ‘Multiplexed samples with Feature Barcode technology for Protein using Barcode Oligo Capture’ protocol (10x Genomics, CG000673, Rev B) for 16-24 hours at 42°C. After probe hybridization, samples were diluted in Post-Hyb Wash Buffer, and a cell aliquot was stained with AO/PI viability dye and then counted with the Nexcelom Cellaca MX cell counter. Counted samples were then pooled at equivalent cell concentrations (when possible), incubated for 3 rounds of 10 minutes at 42°C, washed in Post-Hyb Wash Buffer, and strained using 30uM strainers (Miltenyi Biotec). The multiplexed pool was loaded on to Chip Q for GEM generation at a loading concentration of the number of barcoded samples multiplied by 16,000 cells/barcode. Library preparation was done in accordance with the ‘Multiplexed Samples with Feature Barcode Technology for Protein using Barcode Oligo Capture’ protocol (10x Genomics, CG000673, Rev B). Library quality control was performed using TapeStation D1000 (Agilent Technologies) and Kapa Library Quantification kit (Roche) following manufacturer’s guidelines. Target library peaks expected at 260 base pairs (bp) per TapeStation assay and quantification was measured prior to sequencing. Final single-cell Gene Expression (GEX) and DNA-barcoded Antibody (ADT) libraries were sequenced using a NovaSeq X 25B PE300 or NovaSeq S4 PE100 flow cell, targeting a minimum read depth of 10,000 reads per cell for the GEX libraries and 5,000 for the ADT libraries. ADT and GEX libraries were sequenced at Northwest Genomic Center at the University of Washington at manufacturer’s recommended sequencing guidelines (Read 1: 28 cycles, Read 2: 90 cycles, Index 1: 10 cycles, Index 2: 10 cycles).

#### TEAseq pipeline

HD BMMC samples were enriched for total ASCs, centrifuged, and resuspended in wash buffer. Cells were stained with TotalSeq-A Universal Human cocktail (Biolegend, 399907), HTO antibodies, and a custom TotalSeq-A antibody cocktail (**Table S1**) and incubated at 4°C for 30 minutes. After staining, ASCs were washed twice with wash buffer and transferred to an Eppendorf 96-well plate. Cell counting was completed as described above. Cells were normalized for counts and pooled into 2 pools with equal sample numbers. For both ASC sample pools, individual ATAC, RNA, HTO and ADT libraries were prepared, sequenced and processed as described previously^26^.

#### NDMM BMMC FACS and CITEseq pipeline

Cryopreserved BMMC samples from the MM patient cohort were processed as previously described^70^. Briefly, BMMC were thawed into pre-warmed AIM V media at 37°C (Gibco) and centrifuged at 400xg for 10 minutes at 4°C. Thawed cells were resuspended in AIM V media at 4°C for cell counting and an additional wash. Individual samples were then resuspended in DPBS (Costar) to 10^6^ cells per mL. Up to 10^6^ BMMCs were blocked with Human TruStain FcX Blocking Buffer (BioLegend) following manufacturer’s instructions. Cells were then stained and incubated for 30 minutes with HTOs and a custom pool of DNA-barcoded antibodies (ADTs) (BioLegend, various PN). Stained cells were washed to remove unbound antibodies and pooled for FACS sorting. To remove dead cells and debris, thawed BMMC samples were sorted by FACS prior to processing via Next Generation Sequencing (NGS) workflows. Cells were resuspended in 300uL DPBS with Fixable Viability Stain 510 (BD Biosciences) and incubated for 30 minutes at 4°C protected from light. Cells were washed once with AIM V medium with 25mM HEPES, passed through a 35um filter and adjusted to 10x10^6^ cells per mL. Cells were sorted on a BD FACS Aria Fusion using a 100um nozzle. A standard viable cell gating scheme was employed; FSC-A vs. SSC-A (to exclude sub-cellular debris), two FSC-A doublet exclusion gates (FSC-A vs. FSC-W followed by FSC-H), followed by a dead cell exclusion gate (BV510 LIVE/DEAD negative). An aliquot of each post-sort population was used to collect 10,000 events to assess post-sort purity. After dead cell depletion via FACS sorting, cells were partitioned and barcoded using the Chromium Controller (10x Genomics) and the Chromium Next GEM Single Cell 3′ GEM, Library & Gel Bead Kit v3.1 (10x Genomics, PN-1000121). Up to 24 10x wells were loaded with total cells loaded per well ranging from 26,000 to 32,000 cells. At cDNA amplification, HTO-cDNA additive primer and ADT-cDNA additive primer were spiked into the cDNA amplification master mix and cDNA was amplified for 11 cycles following 10x Genomics cycling guidelines. Following cDNA amplification, HTO/ADT and cDNA products were separated using a 0.6X SPRI-Select (Beckman Coulter, PN B23319) bead-based cleanup as described in Swanson et al. (2021), before carrying forward into separate library indexing reactions 73. HTO and ADT libraries were amplified an additional 10 or 12 cycles with custom single-end index primers. Gene expression libraries were prepared in accordance with Chromium Next GEM Single Cell 3ʹ Reagent Kits v3.1 (CG000204, Rev D). Libraries were sequenced on a NovaSeq S4 200 cycle flow cell or NovaSeq X 25B 100 cycle flow cell at Northwest Genomic Center at the University of Washington following the manufacturer’s recommended sequencing guidelines (Read 1: 28 cycles, Read 2: 91 cycles, index 1:8 cycles) targeting a minimum of 30,000 reads per cell for gene expression, 2,500 reads per cell for HTO, and 5,000 reads per cell for ADT libraries. Samples were then computationally resolved and quality-checked using in-house pipelines.

#### MM PRETX Fixed CITEseq

Cryopreserved BMMC samples from the NDMM and RRMM patient cohorts collected at pretreatment timepoints were rapidly thawed at 37°C, diluted into thawing media solution, and then washed twice at 4°C. Washed BMMCs were resuspended in the media solution and filtered twice using 100uM pore size. Filtered BMMC were resuspended in 1:10 diluted Human TruStain FcX (Biolegend) in a wash buffer (0.22µm pore-filtered DPBS supplemented with 1% hi-FBS) and incubated for 10 minutes at RT. Cells were then centrifuged and resuspended in 1X Mojosort Isolation Buffer (Biolegend). To remove dead cells and debris, BMMC were processed with the Mojosort Human Dead Cell Removal Kit (Biolegend) following manufacturer’s instructions. BMMCs were briefly washed once with a wash buffer (0.22µm pore-filtered DPBS supplemented with 1% hi-FBS). Cells were then resuspended in wash buffer, stained with TotalSeq-C Universal Human cocktail (Biolegend, 399905) and Custom Total Seq-C Cocktail (**Table S1**) according to the ‘Cell Surface and Intracellular Protein Labeling for Chromium Fixed RNA Profiling’ protocol (10X Genomics, CG000529, Rev C) and incubated at 4°C for 30 minutes. Following staining, BMMC samples were washed once with wash buffer and then fixed and quenched according to a modified protocol of the ‘Fixation of Cells and Nuclei for Chromium Fixed RNA Profiling’ protocol (CG000478, Rev C), as previously described^69^. Up to 10^6^ fixed cells per sample were barcoded for multiplexing using the Fixed RNA Feature Barcode Multiplexing Kit (10X Genomics, PN 1000628). Probe hybridization was completed according to the ‘Multiplexed samples with Feature Barcode technology for Protein using Barcode Oligo Capture’ protocol (10x Genomics, CG000673, Rev B) for 16-24 hours at 42°C. After probe hybridization, cell samples were counted, pooled and processed as described in post-hybridization steps of the *Healthy BMMC and PBMC Single-Cell CITE Sequencing*. The multiplexed pool was loaded on to Chip Q for GEM generation at a loading concentration of the number of barcoded samples multiplied by 16,000 cells/barcode. Library preparation was done in accordance with the ‘Multiplexed Samples with Feature Barcode Technology for Protein using Barcode Oligo Capture’ protocol (10x Genomics, CG000673, RevB).

Library quality control and sequencing was performed as described in *Healthy BMMC and PBMC Single-Cell CITE Sequencing*. ADT and GEX libraries were sequenced at Northwest Genomic Center at the University of Washington at manufacturer’s recommended sequencing guidelines (Read 1: 28 cycles, Read 2: 90 cycles, Index 1: 10 cycles, Index 2: 10 cycles).

#### Heavy chain isotype cytometry processing and acquisition

Fresh, overnight-shipped BMA samples from healthy IRB consenting adult donors were processed as described in *Fresh BMA Sample Processing.* Total BMMC were then processed for B cells by negative selection as described in *Total B Cell Enrichment.* Enriched B cells were seeded into a 96-well 0.5mL V-bottom plate at 10x10^6^ live cells/mL in culture media and rested for 30 minutes at 37°C and 5% CO₂. Half of the enriched B cells for each donor sample were processed for BCR isotyping via spectral flow cytometry and the other half were processed for ASC subset FACS and secreted antibody isotype analysis. For cytometry processing, cells were next washed with wash buffer, centrifuged at 400xg for 5 minutes at 4°C and incubated with Human TruStain FcX (1:100 dilution; Biolegend) and purified mouse IgG (1:40 dilution; BioRad) in wash buffer for 10 minutes at RT. After incubation, cells were washed again and incubated with 1:400 diluted Fixable Viability Stain 510 (BD Biosciences) in DPBS for 30 minutes at 4°C in the dark. After incubation, cells were washed and then stained with a custom fluorescence-tagged antibody cocktail in wash buffer for 30 minutes at RT (**Table S1**). After staining, cells were washed once and then incubated with 1.6% PFA in DPBS for 10 minutes at RT in the dark. Fixed cells were then washed twice, with centrifugation at 750xg for 5 minutes at RT. Fixed and washed cells were resuspended in Human TruStain FcX (1:100 dilution; Biolegend) and purified mouse IgG (1:40 dilution; BioRad) in Foxp3 / Transcription Factor Perm Buffer (eBioscience) and incubated for 20 minutes at RT in the dark. Permeabilized cells were centrifuged and resuspended in a custom intracellular fluorescence-tagged antibody cocktail in wash buffer (**Table S1**) and incubated for 40 minutes at RT. B cells were washed twice, immediately resuspended to 10x10^6^ live cells/mL in wash buffer and finally acquired on a Aurora 5L spectral flow cytometer (Cytek Biosciences).

#### Mitochondria activity and surface phenotype cytometry processing and data acquisition

As described above, fresh BMA samples from HDs were processed for BMMC and then for B cells by negative selection. Enriched B cells were seeded into a 96-well non-treated U-bottom plate at 10x10^6^ live cells/mL in culture media (200uL per well) supplemented with BAFF (1ug/mL, PeproTech) alone and cultured at 37°C with 5% CO₂ for one hour. Cultured B cells were then centrifuged, resuspended in 75nM MitoTracker CMX Red (Thermo Fisher Scientific) in culture media and incubated for 45 minutes at 37°C with 5% CO₂. After MitoTracker staining, cells were next washed with wash buffer, centrifuged at 400xg for 5 minutes at 4°C and incubated with Human TruStain FcX (1:100 dilution; Biolegend) in wash buffer for 10 minutes at RT. After incubation, cells were washed again and incubated with 1:400 diluted Fixable Viability Stain 510 (BD Biosciences) in DPBS for 30 minutes at 4°C in the dark. After incubation, cells were washed and then stained with a custom fluorescence-tagged antibody cocktail in wash buffer for 30 minutes at RT (**Table S1**) After staining, cells were washed once and then incubated with 1.6% PFA in DPBS for 10 minutes at RT in the dark. Fixed cells were then washed twice, with centrifugation at 750xg for 5 minutes at RT. Cells were immediately resuspended to 10x10^6^ live cells/mL in wash buffer and finally acquired on an Aurora 5L spectral flow cytometer (Cytek Biosciences).

#### Mass cytometry acquisition

Cells were suspended in TruStain FcX Fc receptor blocker (BioLegend) for 10 min at room temperature (RT) and washed in cell staining media (CSM: PBS with 0.5% BSA and 0.02% sodium azide and benzonase 25x10^8^ U/mL (Sigma)). Cells were stained in CSM for 30 min at RT, washed in CSM, and resuspended in cisplatin for 5 min to label non-viable cells (Sigma, 0.5 mM final concentration in PBS). Cells were washed in CSM and fixed with 1.6% PFA in PBS for 10 min at RT. Cells were washed in CSM and resuspended in intercalation solution (1.6% PFA in PBS, 0.02% saponin (Sigma) and 0.5 mM iridium-intercalator (Fluidigm)) overnight at 4°C. Before acquisition, cells were washed once in CSM and twice in ddH2O. Cells were filtered through a 35μm nylon mesh cell strainer, resuspended at 1 x 10^6^ cells/mL in ddH_2_O supplemented with 1x EQ four element calibration beads (Fluidigm), and acquired on a CyTOF2 mass cytometer (Fluidigm).

#### BMA processing for secreted antibody and cytokine analysis

Fresh BMA samples from healthy donors were processed for BMMC and then for B cells by negative selection. For each donor sample, half of the enriched B cells were processed for heavy chain isotyping via spectral flow cytometry (as described above) and the other half were processed for ASC subset FACS and secreted antibody isotype analysis. For secreted antibody processing, BM ASC subsets were isolated and prepared as described in *HD BM ASC subset FACS* and *HD BM ASC cultures*.

#### BMA processing for pathogen-specific antibody analysis

Cryopreserved BMMC samples from healthy adult donors were processed as described in *Cryopreserved Healthy BMMC and PBMC Sample Processing.* Total BMMC were then processed for B cells by negative selection as described in *Total B Cell Enrichment.* BM ASC subsets were isolated from total B cell pools and prepared as described above in *HD BM ASC subset FACS* and *HD BM ASC cultures*.

#### ASC culture supernatant processing and data acquisition

After 72-hour culture incubations, cell-free supernatants were collected from all sorted ASC subset and TBC culture wells and either immediately processed for analysis or rapidly frozen and stored at -80°C until further processing. ASC culture supernatants were analyzed using an electro-chemiluminescent-based multiplex immunoassay from Mesoscale Discovery (MSD) for the following analyses: 1) secreted antibody isotype, 2) cytokine secretion, 3) secreted Orthopoxvirus (Smallpox; OPXV) antigen-specific IgG levels^71^, 4) secreted respiratory virus antigen-specific IgG levels, and 5) secreted MeV antigen-specific IgG levels. Manufacturer’s recommended protocols were followed for the kit assays: 1) Isotyping Panel 1 Human/NHP Kit (IgG, IgM, IgD isotypes); 2) V-PLEX Proinflammatory Panel 1 Human Kit (IFN-γ, IL-1β, IL-2, IL-4, IL-6, IL-8, IL-10, IL-12p70, IL-13, TNFα); 3) V-PLEX Orthopoxvirus Panel 1 (IgG) Kit (VACV A27L, VACV A33R, VACV B5R, VACV D8L, VACV L1R antigens); and 4) V-PLEX Respiratory Panel 7 (IgG) Kit (Flu A/Ghana/39/2021 H5, Flu A/Massachusetts/18/2022 H3, Flu A/Wisconsin/67/2022 H1, Flu B/Austria/2021 HA, RSV Pre-Fusion F, SARS-CoV-2 N, SARS-CoV-2 Spike, SARS-CoV-2 Spike KP.2, and SARS-CoV-2 Spike XBB.1.5 v2 antigens). For the commercially available antibody quantification assays (1, 3, 4; MSD), a calibrator series (reference standard) was prepared from a pre-validated calibrator stock per manufacturer’s instructions. Blank controls were prepared using Diluent-100 (MSD) alone and culture media alone. Three orthopoxvirus and three respiratory virus serology controls were provided with the Orthopoxvirus Panel 1 and Respiratory Panel 7 IgG kits, respectively. For the custom MeV antigen-specific IgG assay (4; MSD), the 96-well plate precoated with MeV HA antigen was blocked for 30 minutes in Blocker A solution (MSD). As a manufacturer’s calibrator blend was not included with this custom-coated plate, a dilution series of serology-tested control plasma samples, described in *Custom Virus Serology Control Samples*, were included in the plate analysis to determine positivity thresholds. Culture supernatants and control samples were added to the plate at 50uL per well and assay plates were incubated for 2 hours. Following incubation, 1X SULFO-TAG anti-human IgG detection antibody (MSD) was added and the plate incubated for 2 hours. After incubation, GOLD Read Buffer B (MSD) was added to the plate, immediately followed by data acquisition on the SECTOR® S 600 plate reader (MSD). Plates were washed 3x with 300uL Wash Buffer (MSD) between all plate processing steps. All plate incubations were performed at RT and in the dark while shaking at a speed of 700rpm.

#### BM ASC confocal microscopy

ASC subsets and transitional B cells were isolated from fresh BMA samples of healthy adult donors, as described above in *HD BM ASC Subset Fluorescence-Activated Cell Sorting*. Sorted cells were immediately seeded onto 96-well glass bottom plates (CellVis) pre-coated with 0.1% gelatin (Millipore) and seeded in culture media for 16 hours at 37°C and 5% CO_2_. Cells were then stained with 100nM Mitotracker Red CMXRos (ThermoFisher Scientific) in culture media for 45 minutes at 37°C and washed with DPBS. After Mitotracker staining, cells were fixed with 1.6% PFA (Electron Microscopy Sciences) and 0.1% Triton (Sigma-Aldrich) in DPBS for 15 minutes at RT, washed with DPBS, and finally incubated with 3% BSA (Fisher Scientific) in DPBS for 16 hours at 4°C. Cells were stained with 7.5ug/mL of anti-Protein Disulfide Isomerase AF488 (PDI; ThermoFisher Scientific) for 2 hours at RT. Cells were then washed with DPBS prior to staining with NucBlue Fixed Cell Staining (Invitrogen) for 15 minutes at RT. After staining, cells were washed and mounted in SlowFade Gold antifade (Invitrogen) prior to image acquisition. Fluorescence volumetric images were acquired on a Nikon Ti2-E motorized inverted CSU-W1 SoRa Super-Resolution, 7-line Laser Unit, High QE sCMOS, Spinning Disk Confocal microscope using a Chromatic Aberration Free Infinity (CFI) Plane Apochromat Lambda 60X/1.42 NA oil objective. Super Resolution imaging was achieved with the SoRa pinhole setting and acquisition of Z stacks at 0.5 Nyquist step size. Differential Interference Contrast (DIC) images were acquired on a Nikon Ti2-E motorized inverted A1R line scanning confocal microscope equipped with a Transmission Detector using a CFI Plan Apochromat Lambda 60X/1.42 NA oil objective with prisms for DIC.

#### Formalin-fixed-paraffin-embedded (FFPE) tissue sectioning

BM tissue core FFPE blocks (n=2; Accio Biobank Online) were stored at -20°C in sealed plastic bags. Microtome (Leica Microsystems), forceps, dissecting needle, other sectioning tools, and surrounding area were cleaned with RNAseZAP (Invitrogen) and then 70% ethanol (VWR). For sectioning, blocks were rehydrated in RNAse-free water (Cytiva) in a petri dish over ice for up to 30 minutes. Blocks were then sectioned at 5μm and tissue ribbons are floated on a Milli-Q ultrapure water bath that has been heated to 38-41°C. Flattened sections without any bubbles, wrinkles, folds, or other aberrations were collected on a Superfrost plus slide (VWR). Slides were then dried for an hour or until no moisture was visible under the sections. Once dry, slides were stored with desiccant and vacuum sealed until further sample processing.

#### Tissue processing, staining and spatial proteomics image acquisition

BM tissue section slides were equilibrated to room temperature overnight to prevent cold shrinkage of the tissue that would interfere with autofluorescence subtraction. Solutions for staining and imaging were freshly prepared as follows: antibody diluent buffer (3% BSA in 1X PBS (Sigma)), Tween Wash Buffer (0.01% Tween-20 (Sigma) in PBS), DAPI (Thermo Fisher Scientific) staining solution (1 ug/mL in 1X PBS), mounting media (50% glycerol (Sigma), 0.8% propyl gallate (Sigma) in 1X PBS), and storage media (50% glycerol in 1X PBS). The pH of 0.5M sodium bicarbonate buffer (Sigma) was adjusted to a pH of 10.9–11.3 and verified weekly before use. Slides were loaded into the standard CellDIVE ClickWell chambers and followed these staining steps per round of antibody staining. ROIs encompassing all of the tissue were selected using 2X and 10X DAPI imaging. Slides were first imaged with a DAPI only stain to calibrate the autofluorescence, then stained with custom antibody cocktail (**Table S1**) to be imaged for that round of imaging, and finally the sample underwent a dye inactivation before the next round of imaging. Each slide was stained with primary antibodies and diluted in the antibody diluent buffer to a final volume of 250μL per well, per each round of imaging, for 1 hour at RT on a shaker at 250rpm. After staining incubation was completed, the slide underwent three washes with the Tween Wash Buffer, and DAPI staining solution was added for 2 minutes. The slides were washed again with the Tween Wash Buffer and then mounted with mounting media. Slides were imaged at 20X magnification on the CellDIVE system on autofluorescence and biomarker channels. The Cell DIVE performed autofluorescence subtraction and produced a biomarker image. After each imaging round, fluorophores were chemically inactivated by incubating slides for 15 minutes in freshly prepared (within 30 minutes of use) dye inactivation buffer, consisting of 0.5 M Sodium Bicarbonate, 30% Hydrogen Peroxide (Sigma). Slides were washed three times in Tween Wash Buffer and either re-stained for subsequent imaging rounds or stored in Storage Solution at 4°C in the dark. Sequential staining and imaging cycles were repeated until the full fluorescence-tagged antibody panel was completed. All image acquisition and registration were performed using the Leica CellDIVE software platform with pre-selected region-of-interest (ROI) selection between rounds.

### QUANTIFICATION AND STATISTICAL COMPUTATIONAL ANALYSIS

#### Cytometry preprocessing

Original mass cytometry data were bead normalized^72^. Published flow cytometry data from Weisel et al.^22^ and normalized mass cytometry data from Glass et al.^23^ were downloaded from public repositories. Cytometry data were compensated^73^ and uploaded to CellEngine.com for gating (**Fig. S1A-B**). Individual cell surface molecules were manually gated, and percent positive statistics were downloaded for analysis in R.

#### Cytometry Data Processing and Visualizations

Spectral cytometry data were preprocessed in Cytek SpectroFlo v4.0.3 software. Briefly, spectral unmixing was calculated using single color controls generated from cells or FSP compensation particles (Cytek Biosciences). 10,000 events were recorded for beads and 50,000-100,000 events for cells as reference controls on the 5-laser Cytek Aurora cytometer. Spectral unmixing (Ordinary Least Squares method) was calculated in the SpectroFlo software (Cytek, Version 3.3). Flow Cytometry Standard (FCS) files were exported in FCS 3.1 format. Spillover was corrected on the Unmixed data, as needed, and gated in CellEngine (CellCarta, Montreal, Canada). Graphical visualizations of processed cytometry data, aside from gating, were generated in R.

#### ASC secreted antibody processing and analyses

Sample concentrations within each assay were determined using the Discover Workbench (v4.0; MSD) software. Analyte concentrations in arbitrary units (AU per mL) or picograms (pg) per mL were calculated relative to calibrator series or Reference Standard (MSD), when available. Sample dilution factors were accounted for when calculating concentration from each assay standard curve. Graphical visualizations of concentration data were generated in R.

#### Single-cell sequencing preprocessing

Reads were demultiplexed, preprocessed, and quality filtered using Cell Ranger and BarMixer, as previously described^74^. Doublets were removed with Scrublet^75^ and ADTs were denoised using DSB^76^. Further integrated multi-modal data processing and analyses were performed using Seurat^77^. Cells were clustered and cell type labels were imputed from Azimuth bone marrow datasets^28^. Cells which were labeled as “Plasma” and found in clusters with majority “Plasma” labels were kept for downstream analyses. For analysis of MM longitudinal BM CITEseq, “Plasma” annotations from a sister manuscript^51^ analyzing the same data were used for consistency. Before removal of non-ASCs, ADT expression was compared between cell types, and any ADT which was not expressed on ASCs or which displayed non-specific staining were removed (**Fig. S2A**). For analysis of patients with MM PRETX BM fixed CITEseq data, ADTs not expressed by ASCs were kept for downstream comparisons with other immune cell types (**Fig. S15D**). Cells were assigned to the heavy chain subisotype that had the highest scaled expression of all IGH genes (**Fig. S2B**). Cells below a threshold, set for each IGH gene, were re-labeled as not determined (ND). RNA transformation^78^ and PC generation was re-run after removal of non-ASCs. Bone marrow CITEseq data were collected in two batches, so RNA PCs were aligned using Harmony^79^ for batch correction. IGH and IGV genes were excluded from PC generation. For each batch, ADTs were winsorized, min/max scaled, and mode-centered. Batches were then combined, min/max scaled, and clipped so each batch had the same total range (**Fig. S2C**). Thirteen ADTs were added in the second batch, so their expression profiles were imputed onto the first batch using XGBoost^80^ (**Fig. S2D**). ADT PCs were then aligned with Harmony. Ig proteins were excluded from PC generation. Optimal numbers of PCs were selected as the elbow point of variance explained against PC biaxials and used to generate wNN graphs^28^. Cells were clustered using the smart local moving (SLM) algorithm^81^ on a wide range of clustering resolutions, and the resolution with the highest modularity score^82^ was selected for downstream analysis (**Fig. S2E**). Cells were manually gated (**Fig. 1E**) and clusters were assigned to subsets based on the highest gate frequency of the cluster (**Fig. S3C**). Because of the highly concordant staining patterns amongst surface markers (**Fig. S3A**), the gating strategy was tailored to available markers on the panel for other datasets. For the HD BM TEAseq data, CD11a and CD31 were replaced with CD35 and CD63; for the HD matched fixed CITEseq, CD31 and CD9 were replaced with CD54 and HLA-ABC; for the MM 3’ CITEseq, CD11a, CD31, and CD9 were replaced with CD45RA, CD319, and CD47; for the MM fixed CITEseq, no markers were substituted. Light chain isotype was manually gated from surface protein staining (**Fig. S5B**) and surface Ig expression level was defined as the corresponding light chain isotype expression level.

#### ATACseq preprocessing

TEAseq cells were processed and annotated using RNA and ADT modalities as described above. Cells from the RNA module were matched to those from the ATAC module using original barcodes and well IDs. Any mismatched cells containing only RNA or only ATAC modules were excluded. Peak calling was performed using MOCHA^41^. scATACseq fragments were aggregated into subtype-by-sample pseudobulk matrices and sample-specific 500 bp tiles were identified. Consensus peaks were defined as regions accessible in at least two of six samples. Differential accessibility analysis was conducted using a zero-inflated linear mixed-effects model. The model was fit with the formula: Accessibility ∼ CellType + FragmentCounts + (1|Sample) with a zero inflated component specified as ∼ FragmentCounts + CellType. This model assessed pseudobulk chromatin accessibility as a function of ASC subset, while controlling for the total number of fragments per cell type per sample and including sample as a random effect. FDR <0.1 were considered significant. Peaks were annotated using the UCSC hg38 reference genome, and regions located within 2,000 bp upstream and 200 bp downstream of the transcription start site (TSS) were classified as promoter regions. TF activity was inferred at the pseudobulk level using chromVAR^83^. TF deviation scores for each subtype were computed by comparing the MOCHA-derived pseudobulk peak matrix against GC-content-matched background peaks. TF motif accessibility (chromVAR Z-scores) was tested using a one-way ANOVA with sample as a blocking factor to assess differences across cell types. Cell-type-specific effects were identified post hoc using estimated marginal means (EMMs), testing deviation from the overall mean. Motif footprinting plots were generated using the motifFootprint() function in MOCHA, which aggregated normalized chromatin accessibility signal across all instances of a given transcription factor motif within a 200 bp window and corrected for Tn5 insertion bias, enabling visualization of Tn5 insertion patterns around transcription factor binding sites. Normalized coverage data were exported in BigWig format using MOCHA’s exportCoverage() function and visualized as genomic tracks using Gviz^84^.

#### Differential expression analysis

To identify DEPs and DEGs, iterative one-versus-all generalized linear models were employed using MAST^85^ with FDR correction. For subset comparisons, subsets were randomly downsampled to equal cell counts; no downsampling was performed for subisotype comparisons due to low counts of some subisotypes, but subset label was included as a covariate to account for differences in subset frequencies between subisotypes. DEPs were considered significant with Q<0.1 and scaled mean difference > 0.1. DEGs were considered significant with Q<0.1 and log_2_ fold-change > 1. For comparisons between myeloma cells and normal ASC subsets, up to 1000 myeloma cells were sampled per patient, and normal ASC subsets (consisting of non-malignant ASCs from a HD and patients with MM) were equally downsampled (**Fig. S15**)

#### Gene set enrichment analysis

Canonical pathways gene sets from the Pathway Interaction Database^86^ and Hallmark gene sets^87^ from the Molecular Signature Database were filtered to remove gene sets that were irrelevant (e.g. visual signal transduction: cones) or redundant between catalogs (e.g. Hallmark: IL-2 STAT5 signaling and PID: IL-2 STAT5 pathway). Several biologically relevant gene sets from the Gene Ontology Knowledgebase^88^ and the Reactome Knowledgebase^89^ were also included. Genes rank statistics were determined from DEG analysis as -log_10_(adjusted p-value)*log_2_fold-change, and gene set enrichment scores were determined using fgsea^90^ only selecting positively expressed gene sets. Gene sets were visualized on a single-cell level using UCell^91^.

#### ASC gene scores

Multi-gene scores were generated from DEGs to identify subsets in new scRNAseq datasets (**Fig. S4E**, **Table S6**). The top 50 DEGs for each manually gated one-versus-all comparisons were used, except for CD19+9^lo^ASCr, which had too few upregulated DEGs for an accurate gene score, and CD19-ASCr, which was replaced with DEGs comparing ASCr and ASCp for improved accuracy.

#### Weighted shared nearest neighbor distance calculation

Weighted shared nearest neighbor distance graphs generated in single-cell sequencing processing were converted to adjacency matrices using igraph^92^. Edge weights were inverted to represent dissimilarity rather than similarity, and the geodesic distance between all pairs of nodes (cells) was calculated using the distances function from igraph. Infinite values (disconnected elements were replaced with values 10% greater than the maximum distance in the connected graph. For each cell, the average distance to all cells in each subset was calculated and visualized (**Fig. 2A, S6A**).

#### Cell cycle analyses

Cycling cells were identified by gating on Hallmark G2M checkpoint and PID E2F pathway gene set scores (**Fig. S7A**). Cell cycle phase probabilities and labels were calculated from gene expression using Revelio^44^. Cell cycle pseudotime was calculated by performing LDA on cell cycle phase probabilities (predictors) and cell cycle labels (responses) and fitting a principal curve on the first two discriminants (**Fig. S7B**), as previously described^45^.

#### Confocal microscopy imaging data processing, analysis and visualization

Mitochondria and ER channels were processed using MegaSeg, which created 3D probability maps to assign voxels to each organelle. To localize structures in the XY plane, maximum-intensity projections (MIPs) of the 3D probability maps along the z-axis were computed. ER channel MIPs were used to initialize cell outline detection because the ER spans a large cytoplasmic volume and forms extensive ER–mitochondria contact sites^93^. Initial cell masks were generated by applying global Otsu thresholding to ER-channel MIPs^94,95^. These masks were refined using alpha-shape reconstruction (alpha parameter set to 0.1) to obtain concave cell contours that more faithfully captured true morphology. Connected component labeling followed by size filtering removed small, spurious objects; the size threshold corresponded to the mean projected object area across the dataset (n = 941 pixels or 1.41μm). The remaining components were relabeled to produce instance-level masks for individual ASCs. All masks underwent visual quality control to ensure exclusion of cells touching image boundaries, insufficiently separated neighboring cells, and cells below 3μm^2^ in projected area. Approximately 11% (n =157) of masks were excluded, yielding a final set of 1,257 high-quality single-cell masks. The corresponding 3D mitochondrial and ER probability maps were restricted to cellular regions by extruding the 2D outline through the z-dimension. Voxels with probability ≥0.995 (range 0 to 1) were binarized as organelle-positive; all others were set to background (zero). Connected components within each 3D single-cell volume were identified as candidate organelles. A volume filter was applied using the median segmented-object volume computed across all cells (minimum volume = 28 voxels or 0.0084 μm^3^), chosen for robustness to outliers and validated by visual inspection. Final 3D segmentation masks were stored as 8-bit TIFF files.

Morphometric features were extracted from segmented mitochondrial and ER objects within individual ASCs. For each object, volume was computed from voxel occupancy, fluorescence intensity from raw image intensities, and surface area was estimated using marching cubes reconstruction^96,97^ to calculate sphericity, compactness, and shape factor. Network architecture was quantified by 3D skeletonization using the Lee thinning algorithm^98^, followed by graph-based analysis to compute junction density, tubulation index, and endpoint ratio, which were summarized at the single-cell level. One was added to mitochondrial and ER counts to account for cells with zero counts. Likewise, mean mitochondrial intensity, mitochondrial volume, and intersection volume zero values were replaced with 0.9*minimum non-zero value. Mean intensity, volume, and count morphometrics were log_10_ transformed. Network scores and sphericity metrics were left in linear scales. All morphometrics were then z-scored. Violin plots were winsorized at 0.01 and 0.99 percentiles for improved visualization (**Fig. 3H, S8H**).

#### Mean Ig monomer secretion per cell calculations

The concentration of Ig isotypes in culture supernatants was quantified by MSD (**Fig. 4B**), but this measurement does not account for differences in isotype composition between sorted subsets. To correct for this, the frequency of each Ig isotype within each subset from each HD was quantified by flow cytometry and multiplied by the total number of cells deposited per well (**Fig. S9B**). Supernatant concentrations (ng/mL) were converted to total mass (kDa) per well. Molecular weights for were assigned as 180kDa, 150kDa, and 160kDa for IgM, IgG, and IgA monomers, respectively^99^. The calculated mass of each isotype in the culture supernatant (IgH*_sup.mass_*) was divided by the product of the corresponding isotype’s molecular mass (IgH*_mol.mass_*) and the number of cells for that isotype in that well (N*_cells_*):

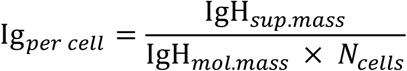

#### Spatial proteomics processing

Multiplexed images were stitched, flat-field corrected, registered, and aligned by the commercial Cell DIVE software (version 4.0.1.15). Cell segmentation was performed in Aivia (version 15.0) with the protocol/recipe ‘Multiplex Cell Detection v1’, which employs nuclear cell detection and user-assigned membrane expansion with modified CellPose segmentation^100^. Membrane expansion markers CD3d, CD3e, CD14, CD16, CD19, CD20, CD31, CD34, CD38, CD44, CD45RB, CD54, HLA-DR, and NaK-ATPase were utilized for cell boundary segmentation. After segmentation, whole cell mean intensities were measured for all channels. Aggregate dye spot removal was performed by excluding cells that had a median fluorescence intensity of 60,000 units or greater in any marker channel. Dye spot removal resulted in a reduction of 1.2% (420 of 34790 total; bone marrow 1) and 6.6% (1527 of 23029 total; bone marrow 2) of total segmented cells.

#### Spatial proteomics cell table processing and clustering

Each image was processed independently in parallel to account for technical variation. Mean cell fluorescence intensities were arcsinh-transformed with custom cofactors selected to maximize separation of positive and negative modes and scaled to the 99.9^th^ percentile. A supervised hierarchical clustering approach was employed, as previously described^101,102^. Cells were overclustered with FlowSOM using COL1A1, COL3A1, CD27, CD31, CD34, CD45, IRF4, and SMA, and clusters were annotated as immune cells or MSCs. Immune cells were subclustered using CD3d, CD14, CD16, CD19, CD20, CD34, CD38, CD39, CD45, CD45RB, CD71, HLADR, IRF4, and Ki67 and annotated into immune cell types (**Fig. S12B**). ASCs were further subclustered using CD3d, CD19, CD38, CD44, CD45, CD45RB, CD71, HLADR, and IRF4 (HD30) or CD3d, CD19, CD31, CD38, CD39, CD44, CD45, CD45RB, CD54, CD71, HLADR, and IRF4 (HD31) and annotated into ASC subsets.

#### Permutation analysis of ASC niches

Each cell’s niche was defined as all cells with centroids within 50μm of the query cell’s centroid. An ASC niche was considered homogenous if all other ASCs in its niche matched the query cell’s phenotype. ASCs with no other ASCs in their niche were removed from the analysis. The observed frequency of homogenous niches was calculated and then ASC subset labels were randomly permuted 1,000 times and the homogenous niche frequency was quantified for each permutation to generate a null distribution (**Fig. 5H**). Empirical p-values were calculated as the number of permuted homogenous niche frequencies greater or equal to the observed homogenous niche frequency, divided by the number of permutations plus one (**Fig. 5I**). A similar, previously published analysis^103^ was applied to identify cell type enrichments in ASC niches. For each ASC niche, the cell type composition was quantified and summed for each ASC subset across all niches as the observed count. Cell type labels were randomly permuted 100 times, and the cell type counts in ASC subset niches were quantified for each permutation to generate a null distribution (**Fig. 5J**). Z-scores of observed counts were calculated using the permuted count distributions (**Fig. 5K**). FDR-corrected empirical Q-values were calculated as the number of permuted cell type counts greater or equal to the observed counts (if enriched) or lesser or equal to the observed counts (if excluded), divided by the number of permutations plus one.

#### Identification of malignant cells

For each patient with MM (**Fig. 7, S14-15**), total ASCs across timepoints were merged into a single object, and a patient-specific wNN graph was calculated and clustered, as described above. Clusters with homogenous heavy chain and light isotype usage, aberrant expression profiles, and an enrichment for cells from pre-treatment timepoints were annotated as malignant. Non-malignant clusters were typically the minority of ASCs in patients and separated within UMAP space (**Fig. S14A**).

## Notes

https://apps.allenimmunology.org/aifi/resources/antibody-secreting-cells/

https://doi.org/10.5281/zenodo.19499360

