## Supplemental Materials for "Multi-omic profiling of human antibody-secreting cells reveals diverse subsets sustain durable humoral immunity"

#### SUPPLEMENTAL TABLE LEGENDS

**Table S1: Antibody panels – related to Fig. 1**

**Table S2: HD cohort characteristics – related to Fig. 1**

**Table S3: HD DEPs – related to Fig. 2**

**Table S4: HD DEGs – related to Fig. 2**

**Table S5: HD DATF motifs – related to Fig. 2**

**Table S6: ASC subset gene sets – related to Fig. 2**

**Table S7: HD GSEA – related to Fig. 2**

**Table S8: Patients with MM cohort characteristics – related to Fig. 7**

**Table S9: MM DEPs – related to Fig. S15**

**Table S10: MM DEGs – related to Fig. S15**

### SUPPLEMENTAL FIGURES

### Figure S1

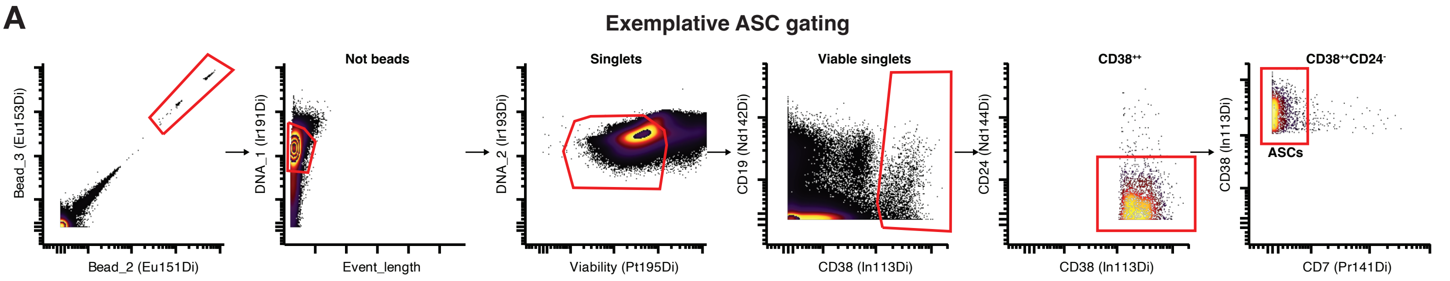


**B**

##### ASC gating schemes by dataset

| Dataset | ASC gating scheme |
|---|---|
| Weisel et al. flow cytometry screen data on human spleen | Singlet+Viable+CD38++ |
| Original mass cytometry screen data on human bone marrow | Singlet+Viable+CD7-CD24-CD38++ |
| Glass et al. mass cytometry data on multiple human tissues | Singlet+Viable+CD24-CD38++ |
| Original mass cytometry data on matched human bone marrow and peripheral blood | Singlet+Viable+Lin-CD24-CD38++ |

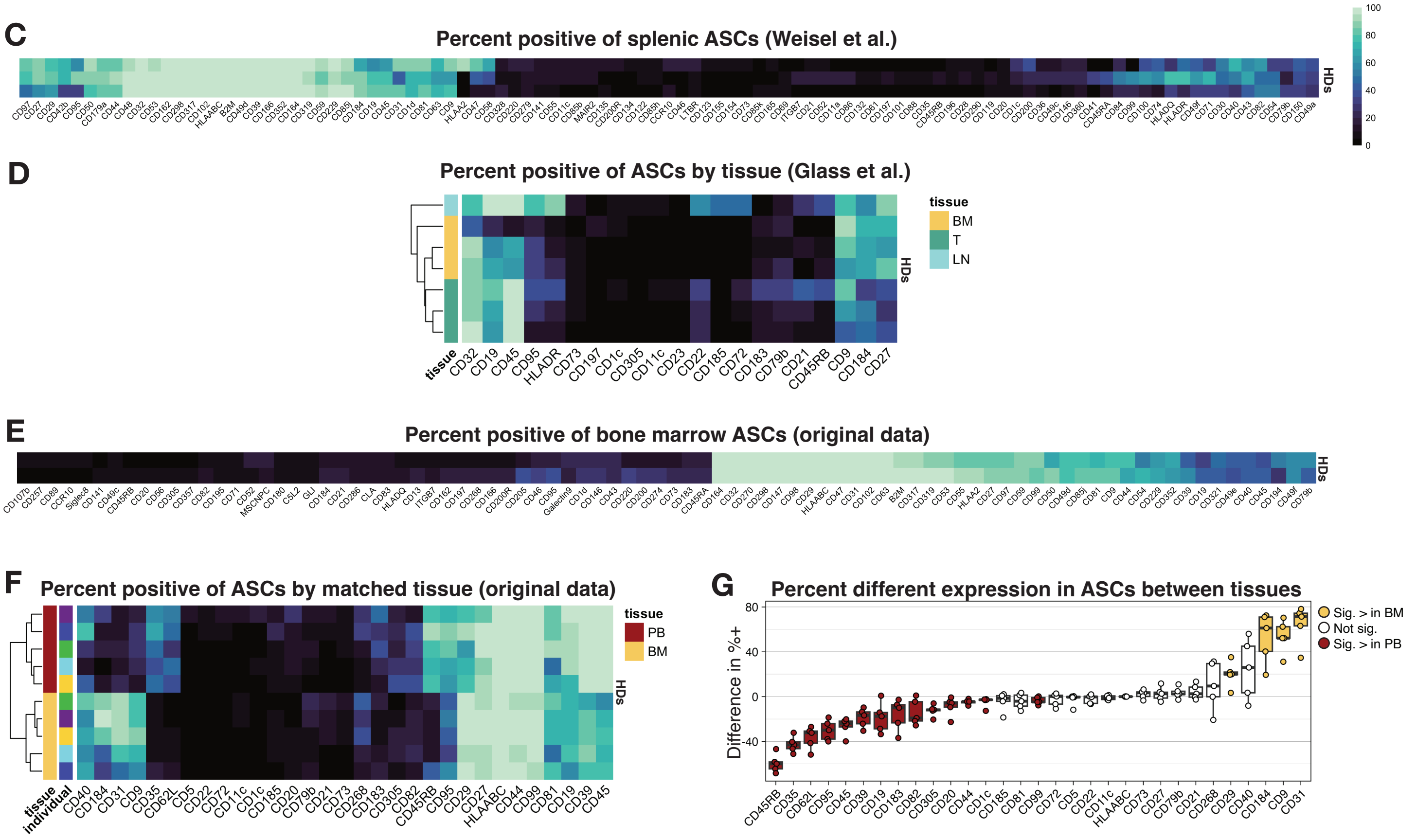

**Figure S1: Cytometry meta-analysis – related to Fig. 1.**

A) Exemplative ASC gating scheme for cytometry meta-analysis. Original data from HD BM mass cytometry screen.
B) Table of gating schemes used for the four cytometry datasets included in the meta-analysis.
C) Percent positive of cell surface proteins on splenic ASCs by individual. Data from publicly available HD flow cytometry screen.
D) Percent positive of cell surface proteins on ASCs by individual and tissue. Data from publicly available HD mass cytometry.
E) Percent positive of cell surface proteins on bone marrow ASCs by individual. Original data from HD mass cytometry screen.
F) Percent positive of cell surface proteins on ASCs by individual and tissue. Original data from HD matched BM and PB mass cytometry.
G) Different in percent positive between BM and PB ASCs by individual (dots). Crossbar indicates median, boxes represent interquartile range (IQR), and whiskers represent IQR +/ 1.5*IQR. Significance defined as $Q<0.1$ using Wilcoxon rank sum test with FDR correction. Data from HD matched BM and PB mass cytometry.

### Figure S2

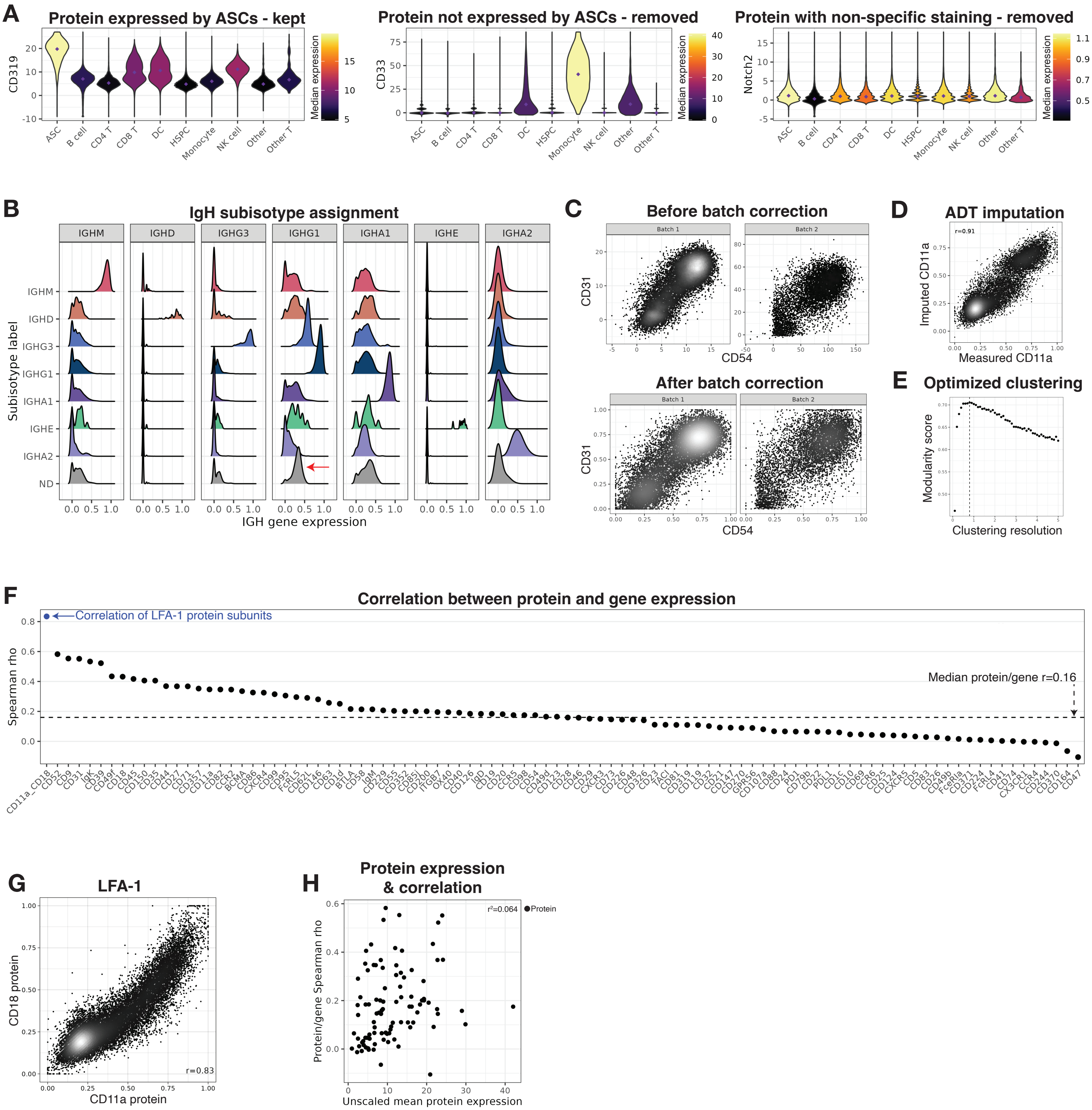
A
Protein expressed by ASCs - kept
CD319
Median expression
Protein not expressed by ASCs - removed
CD33
Protein with non-specific staining - removed
Notch2
ASC
B cell
CD4 T
CD8 T
DC
HSPC
Monocyte
NK cell
Other
Other T
B
IgH subisotype assignment
IGHM
IGHD
IGHG3
IGHG1
IGHA1
IGHE
IGHA2
ND
Subisotype label
IGH gene expression
C
Before batch correction
After batch correction
Batch 1
Batch 2
CD31
CD54
D
ADT imputation
r=0.91
Imputed CD11a
Measured CD11a
E
Optimized clustering
Modularity score
Clustering resolution
F
Correlation between protein and gene expression
Correlation of LFA-1 protein subunits
Spearman rho
Median protein/gene r=0.16
G
LFA-1
CD18 protein
CD11a protein
r=0.83
H
Protein expression
& correlation
Protein/gene Spearman rho
Unscaled mean protein expression
Protein

**Figure S2: Single-cell sequencing preprocessing and subset discovery – related to Fig. 1.**

A) Exemplative plots showing unscaled expression of ADTs by Azimuth cell type. These plots informed filtering of ADTs before ASC-specific analysis. Data from HD BM fixed CITEseq.

B) Exemplative plots showing IGH gene expression by ASC subisotype. ND denotes subisotype not determined. Arrow indicates increased expression of IGHG1 in ND cells (as in IGHG3 labeled cells), suggesting they are enriched in IGHG2 and IGHG4 subisotypes, which were not included in the gene probe set. Data from HD BM fixed CITEseq.

C) Exemplative plots showing ADT batch correction. Data from HD BM fixed CITEseq.

D) Exemplative plot showing accuracy of imputation method used for ADTs absent in batch 1. Plot shows CD11a expression in batch 2 (x-axis) and predicted expression of CD11a in batch 2, trained on batch 1 data (y-axis). Data from HD BM fixed CITEseq.

E) Exemplative plot showing selection of optimal clustering resolution. Cells were clustered at a wide range of resolutions (x-axis) and the resolution with highest modularity score (dashed line) was selected for downstream analysis. Modularity score derived calculated with the R function igraph::modularity. Data from HD BM fixed CITEseq.

F) Spearman correlation between ADT expression and the expression of the corresponding gene. Blue dot identifies the correlation of two ADTs that from a single complex (CD11a & CD18) as a positive control. Dashed line indicates median of protein/gene correlations. Data from HD BM fixed CITEseq.

G) Biaxial plot surface protein expression of LFA protein subunits. R indicates Spearman correlation highlighted in F. Data from HD BM fixed CITEseq.

H) Biaxial of unscaled mean expression of ADTs and correlation coefficient between ADT and underlying gene expression as in F. Adjusted r-squared, calculated using linear regression, shows protein expression level does not inform correlation strength. Data from HD BM fixed CITEseq.

### Figure S3

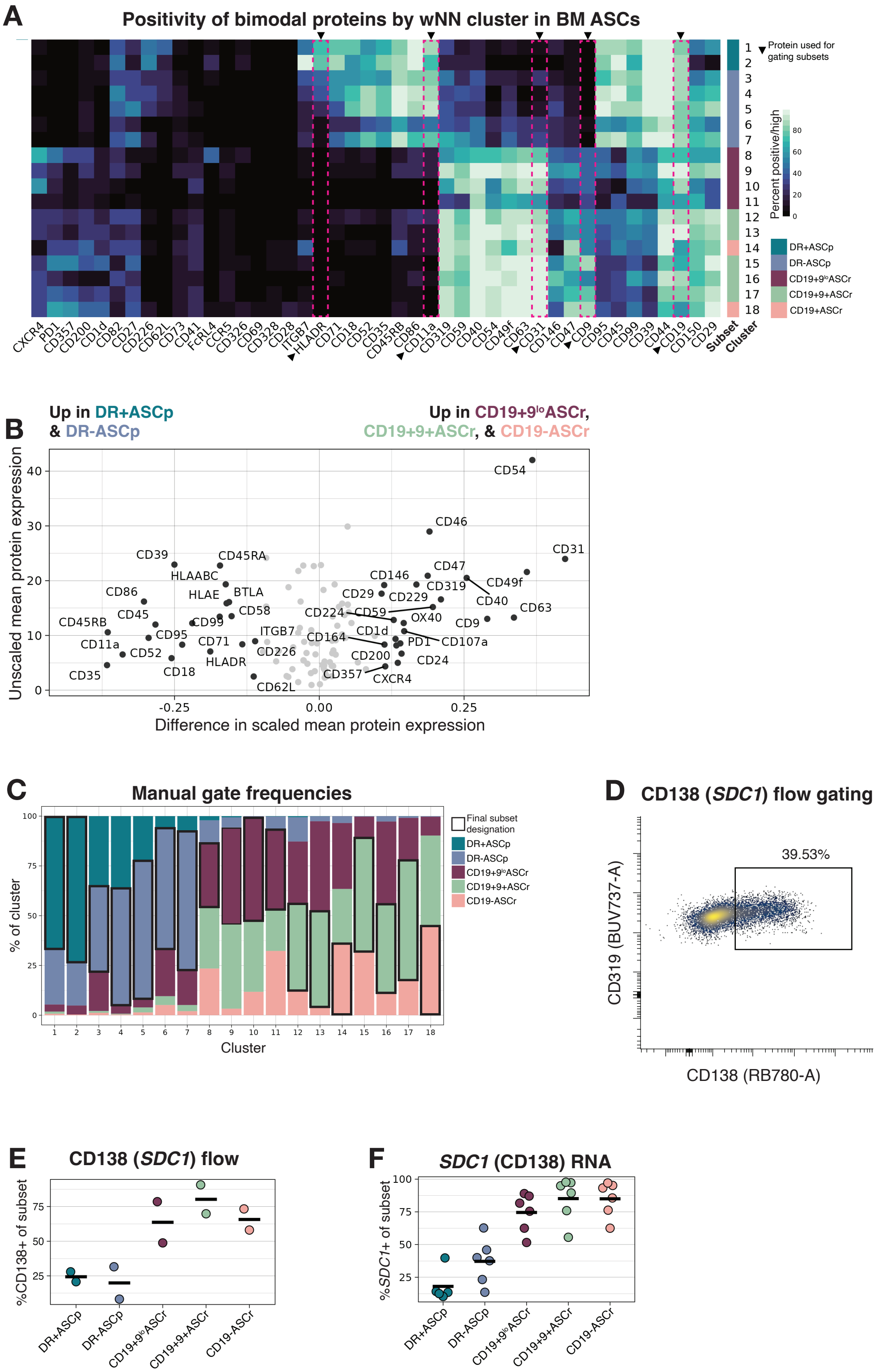
A
Positivity of bimodal proteins by wNN cluster in BM ASCs
Protein used for gating subsets
Percent positive/high
DR+ASCp
DR-ASCp
CD19+9loASCr
CD19+9+ASCr
CD19+ASCr
CXCR4 PD1 CD357 CD200 CD1d CD82 CD27 CD226 CD62L CD73 CD41 FcRL4 CCR5 CD326 CD69 CD328 CD28 ITGB7 HLADR CD71 CD18 CD52 CD35 CD45RB CD86 CD11a CD319 CD59 CD40 CD54 CD49f CD63 CD31 CD146 CD47 CD9 CD95 CD45 CD99 CD39 CD44 CD19 CD150 CD29
Subset
Cluster
B
Up in DR+ASCp & DR-ASCp
Up in CD19+9loASCr, CD19+9+ASCr, & CD19-ASCr
Unscaled mean protein expression
Difference in scaled mean protein expression
CD54 CD46 CD31 CD39 CD45RA HLAABC CD146 CD47 CD49f CD86 HLAE BTLA CD29 CD229 CD319 CD40 CD45 CD58 CD224 CD59 CD63 CD45RB CD99 OX40 CD9 CD95 ITGB7 CD1d CD11a CD71 CD164 PD1 CD107a CD52 CD226 CD18 HLADR CD200 CD24 CD35 CD357 CXCR4 CD62L
C
Manual gate frequencies
Final subset designation
DR+ASCp
DR-ASCp
CD19+9loASCr
CD19+9+ASCr
CD19-ASCr
% of cluster
Cluster
D
CD138 (SDC1) flow gating
39.53%
CD319 (BUV737-A)
CD138 (RB780-A)
E
CD138 (SDC1) flow
%CD138+ of subset
DR+ASCp
DR-ASCp
CD19+9loASCr
CD19+9+ASCr
CD19-ASCr
F
SDC1 (CD138) RNA
%SDC1+ of subset
DR+ASCp
DR-ASCp
CD19+9loASCr
CD19+9+ASCr
CD19-ASCr

**Figure S3: ASC surface protein characterization– related to Fig. 1.**

A) Percent positive/high of ADTs with clear bimodal expression in ASCs by cluster. Clusters are colored by subset. Data from HD BM fixed CITEseq.
B) Differences in scaled mean ADT expression (x-axis) between ASCp against ASCr, and unscaled mean expression of ADTs (y-axis) of total ADTs expressed by ASCs. Labeled proteins are significantly differentially expressed. Data from HD BM fixed CITEseq.
C) Percent of each manual gate by Seurat cluster. Black boxes indicate the highest prevalence gate in each cluster, which was used to assign subsets. Data from HD BM fixed CITEseq.
D) Exemplative CD138 gate on total $CD38^{hi}CD319+$ ASCs. Data from HD BM flow cytometry.
E) Percent *SDC1*+ of ASC subsets. Crossbar indicates mean. Data from HD BM fixed CITEseq.
F) Percent CD138+ of ASC subsets. Crossbar indicates mean. Data from HD BM flow cytometry.

### Figure S4A-B

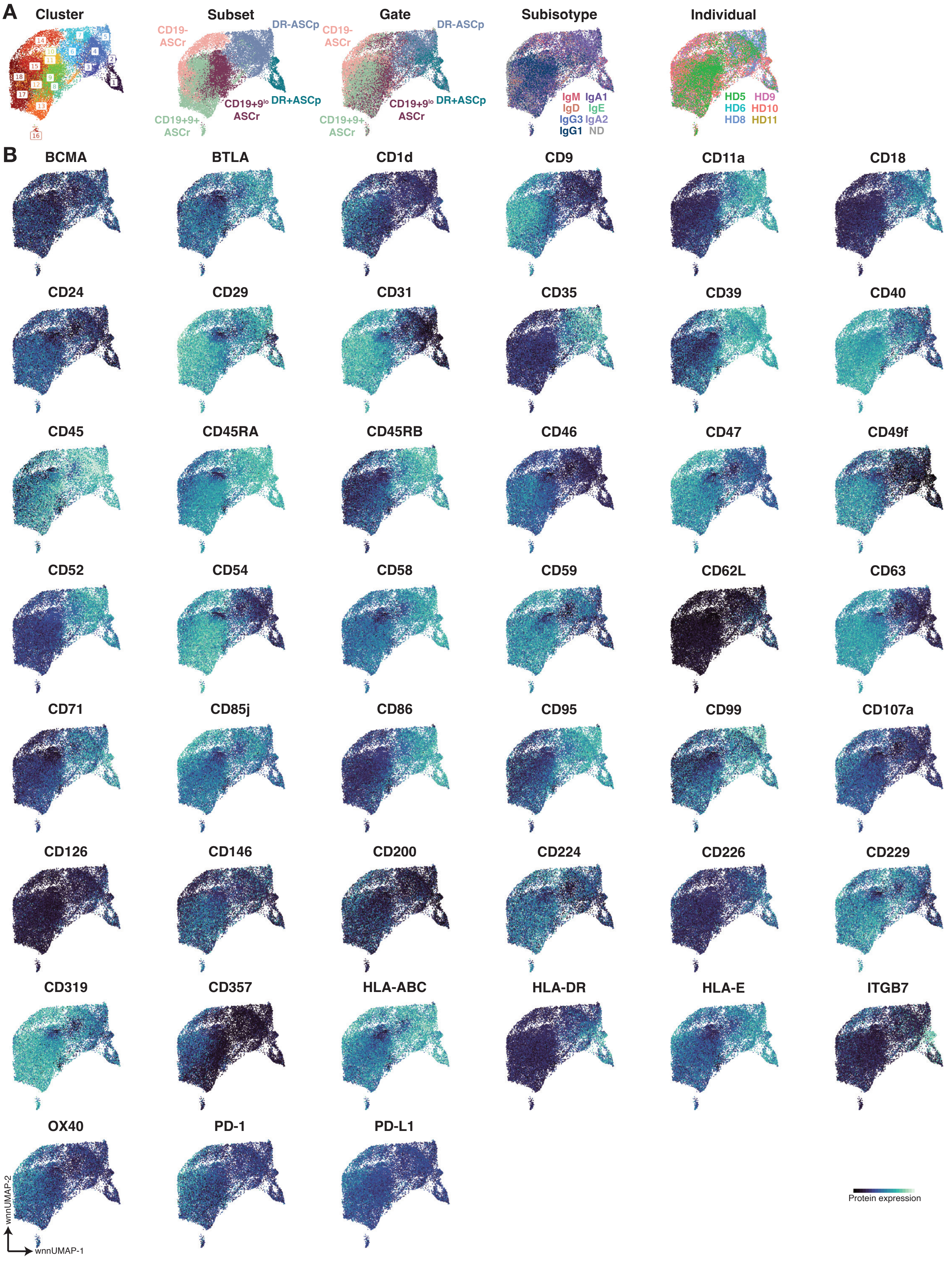

A
Cluster
Subset
Gate
Subisotype
Individual
CD19-
ASCr
DR-ASCp
CD19+9lo
ASCr
DR+ASCp
CD19+9+
ASCr
IgM
IgA1
IgD
IgE
IgG3
IgA2
IgG1
ND
HD5
HD9
HD6
HD10
HD8
HD11
B
BCMA
BTLA
CD1d
CD9
CD11a
CD18
CD24
CD29
CD31
CD35
CD39
CD40
CD45
CD45RA
CD45RB
CD46
CD47
CD49f
CD52
CD54
CD58
CD59
CD62L
CD63
CD71
CD85j
CD86
CD95
CD99
CD107a
CD126
CD146
CD200
CD224
CD226
CD229
CD319
CD357
HLA-ABC
HLA-DR
HLA-E
ITGB7
OX40
PD-1
PD-L1
wnnUMAP-2
wnnUMAP-1
Protein expression

### Figure S4C

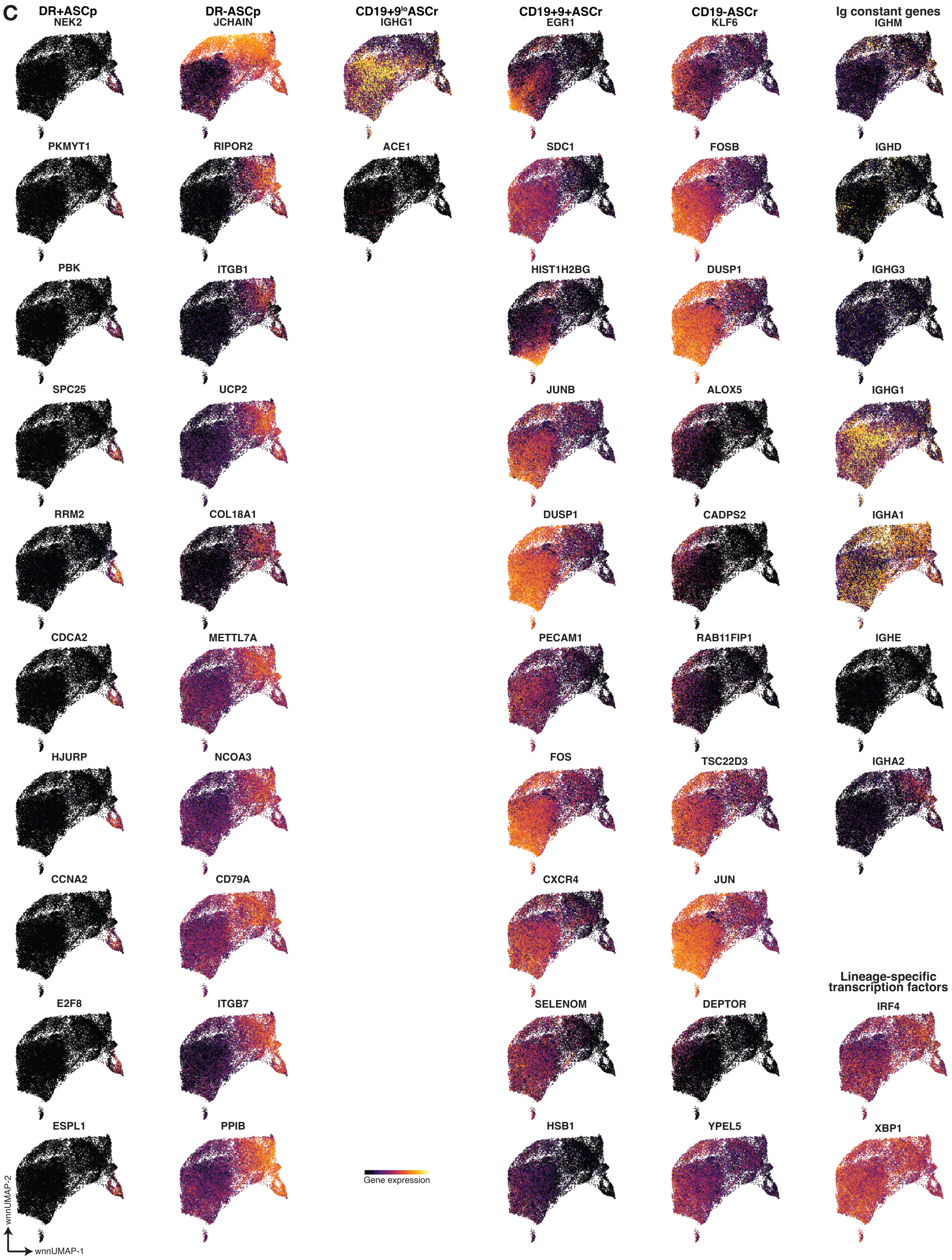
C
DR+ASCp
NEK2
PKMYT1
PBK
SPC25
RRM2
CDCA2
HJURP
CCNA2
E2F8
ESPL1
DR-ASCp
JCHAIN
RIPOR2
ITGB1
UCP2
COL18A1
METTL7A
NCOA3
CD79A
ITGB7
PPIB
CD19+9$^{lo}$ASCr
IGHG1
ACE1
Gene expression
CD19+9+ASCr
EGR1
SDC1
HIST1H2BG
JUNB
DUSP1
PECAM1
FOS
CXCR4
SELENOM
HSB1
CD19-ASCr
KLF6
FOSB
DUSP1
ALOX5
CADPS2
RAB11FIP1
TSC22D3
JUN
DEPTOR
YPEL5
Ig constant genes
IGHM
IGHD
IGHG3
IGHG1
IGHA1
IGHE
IGHA2
Lineage-specific transcription factors
IRF4
XBP1
wnnUMAP-2
wnnUMAP-1

### Figure S4D-E

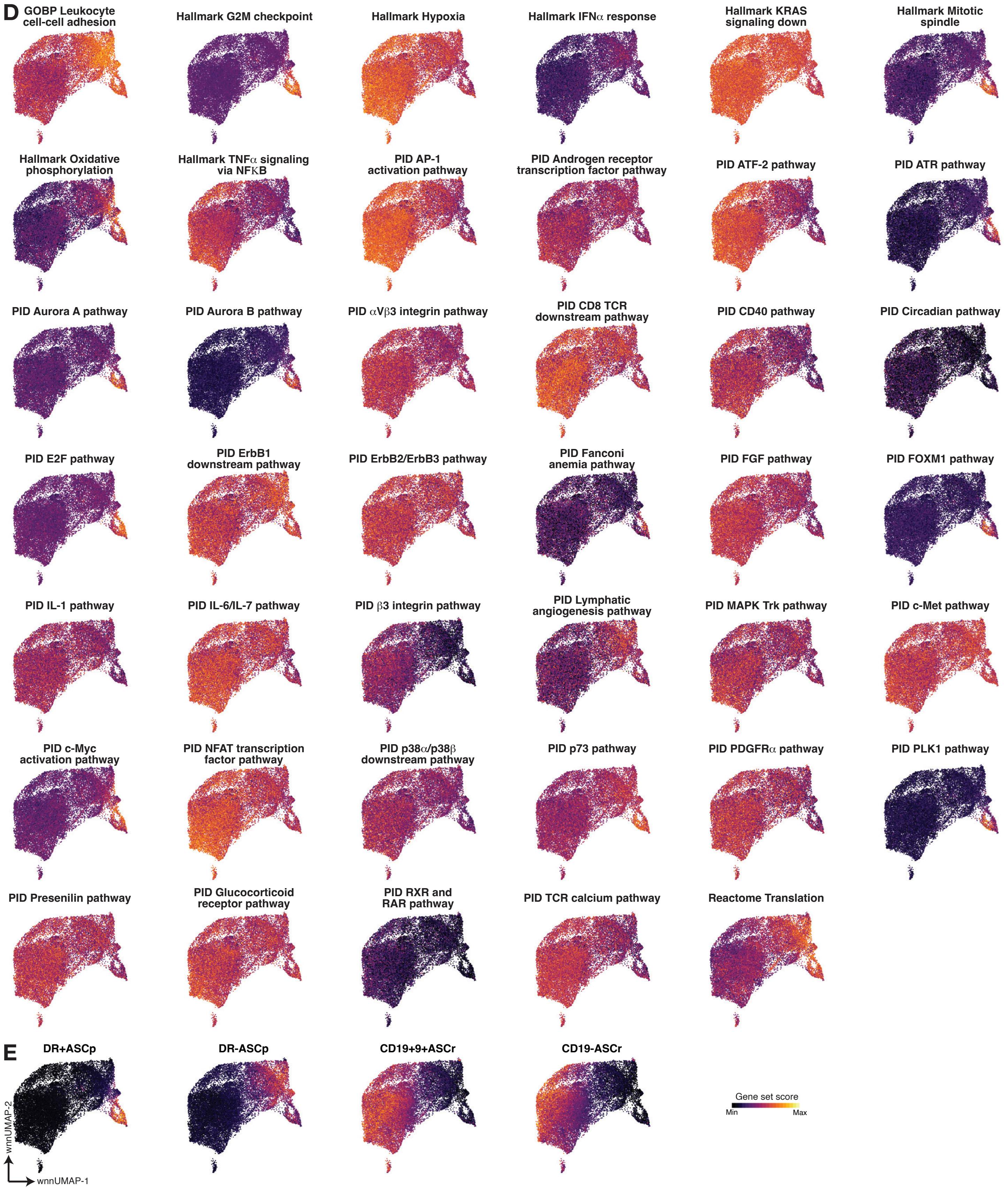
D
GOBP Leukocyte cell-cell adhesion
Hallmark G2M checkpoint
Hallmark Hypoxia
Hallmark IFNα response
Hallmark KRAS signaling down
Hallmark Mitotic spindle
Hallmark Oxidative phosphorylation
Hallmark TNFα signaling via NFKB
PID AP-1 activation pathway
PID Androgen receptor transcription factor pathway
PID ATF-2 pathway
PID ATR pathway
PID Aurora A pathway
PID Aurora B pathway
PID αVβ3 integrin pathway
PID CD8 TCR downstream pathway
PID CD40 pathway
PID Circadian pathway
PID E2F pathway
PID ErbB1 downstream pathway
PID ErbB2/ErbB3 pathway
PID Fanconi anemia pathway
PID FGF pathway
PID FOXM1 pathway
PID IL-1 pathway
PID IL-6/IL-7 pathway
PID β3 integrin pathway
PID Lymphatic angiogenesis pathway
PID MAPK Trk pathway
PID c-Met pathway
PID c-Myc activation pathway
PID NFAT transcription factor pathway
PID p38α/p38β downstream pathway
PID p73 pathway
PID PDGFRα pathway
PID PLK1 pathway
PID Presenilin pathway
PID Glucocorticoid receptor pathway
PID RXR and RAR pathway
PID TCR calcium pathway
Reactome Translation
E
DR+ASCp
DR-ASCp
CD19+9+ASCr
CD19-ASCr
Gene set score
Min
Max
wnnUMAP-2
wnnUMAP-1

**Figure S4: wnnUMAPs – related to Fig. 1.**

A) wnnUMAPs colored by various identities. Data from HD BM fixed CITEseq.
B) wnnUMAPs colored by ADT expression, showing only DEPs, as in Fig. S6D, Table S3. Data from HD BM fixed CITEseq.
C) wnnUMAPs colored by GEX expression, showing up to top ten upregulated DEGs by subset, as in Fig. S6D, Table S4. IgH genes and lineage-specific transcription factor overlays are also shown. Data from HD BM fixed CITEseq.
D) wnnUMAPs colored by gene set score, showing only DEGSs, as in Fig. 2F, Table S7. Data from HD BM fixed CITEseq.
E) wnnUMAPs colored by gene set score, showing scores generated to identify ASC subsets in scRNAseq data, as in Table S6. Data from HD BM fixed CITEseq.

### Figure S5

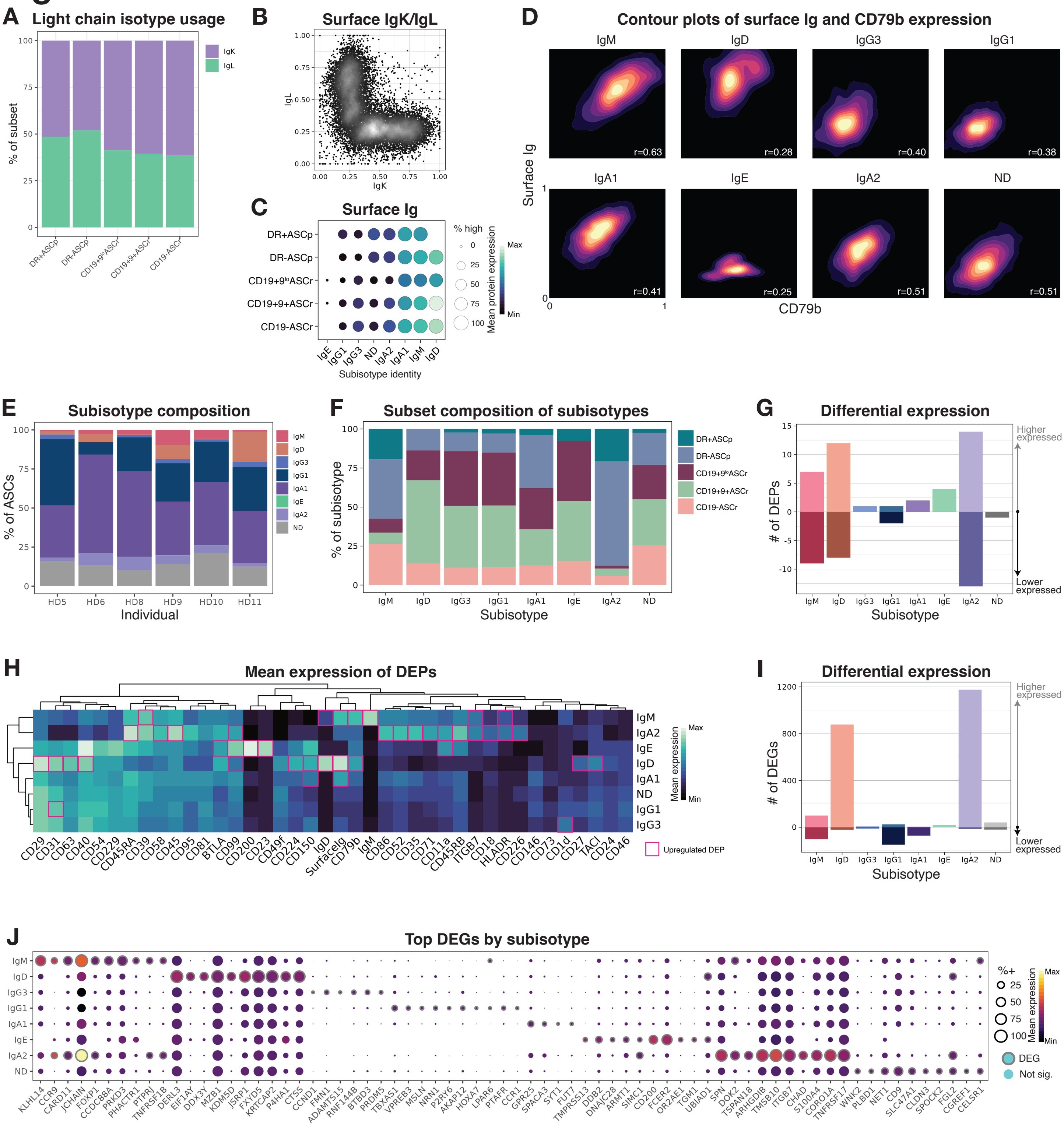
A
Light chain isotype usage
% of subset
IgK
IgL
DR+ASCp
DR-ASCp
CD19+9loASCr
CD19+9+ASCr
CD19-ASCr
B
Surface IgK/IgL
IgK
IgL
C
Surface Ig
% high
Mean protein expression
Max
Min
Subisotype identity
D
Contour plots of surface Ig and CD79b expression
IgM
IgD
IgG3
IgG1
IgA1
IgE
IgA2
ND
r=0.63
r=0.28
r=0.40
r=0.38
r=0.41
r=0.25
r=0.51
r=0.51
Surface Ig
CD79b
E
Subisotype composition
% of ASCs
Individual
HD5
HD6
HD8
HD9
HD10
HD11
F
Subset composition of subisotypes
% of subisotype
Subisotype
G
Differential expression
### of DEPs
Higher expressed
Lower expressed
Subisotype
H
Mean expression of DEPs
Mean expression
Upregulated DEP
I
Differential expression
### of DEGs
Higher expressed
Lower expressed
Subisotype
J
Top DEGs by subisotype
%+
Mean expression
DEG
Not sig.

**Figure S5: Subisotype usage captures a facet of ASC identity – related to Fig. 1.**

A) Percent of light chain isotype used by subset. Data from HD BM fixed CITEseq.
B) Biaxial plot of ADT surface light chain staining. Data from HD BM fixed CITEseq, first batch.
C) Surface Ig (derived from ADT surface light chain staining, as in B) mean expression (color) and percent positive (size) by subset (y-axis) and subisotype identity (x-axis). Absent elements indicate less than five cells present. Data from HD BM fixed CITEseq, first batch.
D) Contour plots of surface Ig and CD79b by subisotype. Data from HD BM fixed CITEseq, first batch.
E) Percent IgH subisotype usage by individual. ND denotes subisotype not determined. Data from HD BM fixed CITEseq.
F) Percent subset identity by subisotype. Data from HD BM fixed CITEseq.
G) Number of DEPs by subisotype. Data from HD BM fixed CITEseq.
H) Mean expression of DEPs by subisotype. Significantly upregulated DEPs are boxed. Data from HD BM fixed CITEseq.
I) Number of DEGs by subisotype. Data from HD BM fixed CITEseq.
J) Mean expression (color) and percent positive (size) of up to top ten DEGs by subisotype. Outline indicates statistical significance. IgH genes removed from analysis. Data from HD BM fixed CITEseq.

### Figure S6

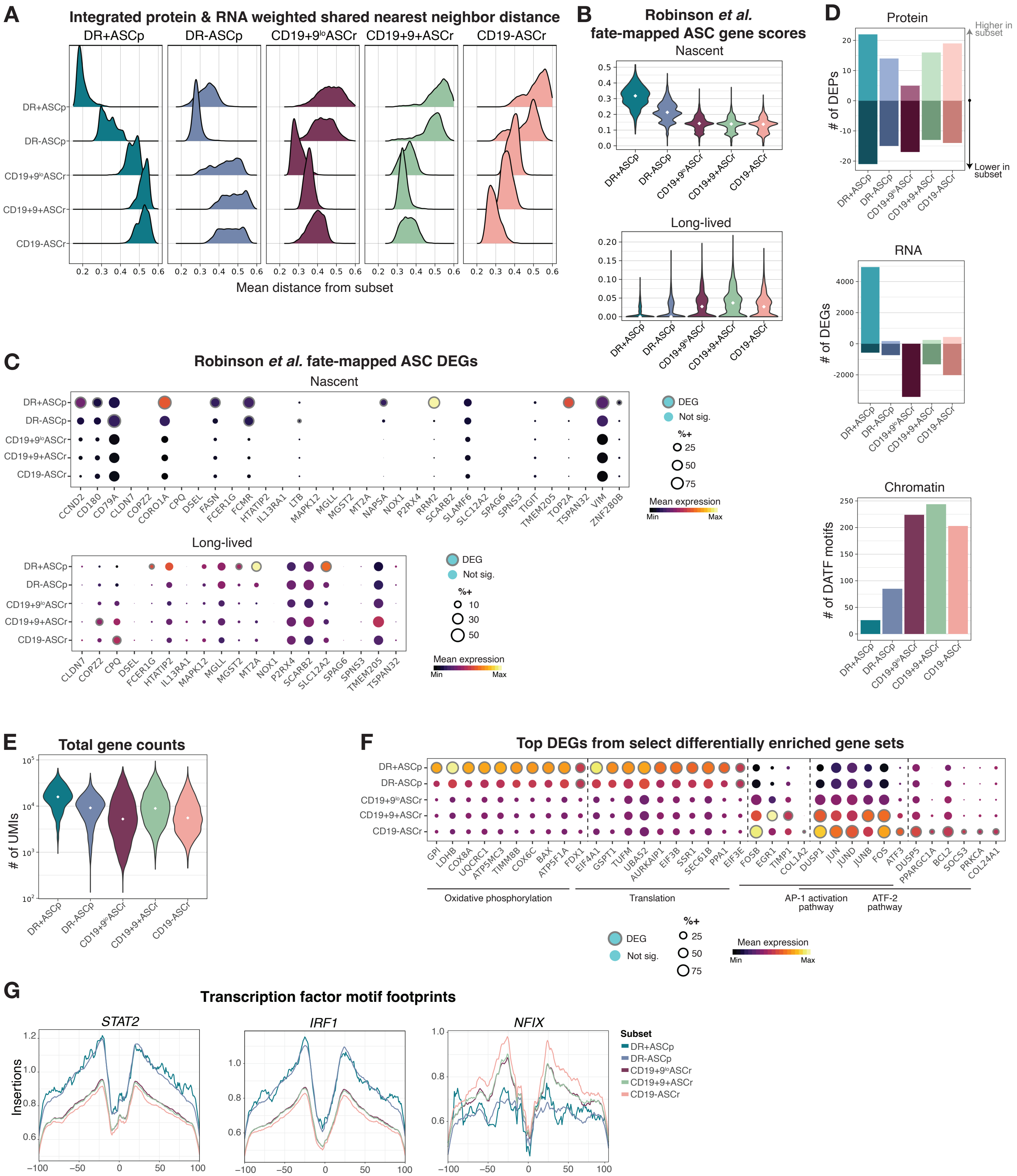
A
Integrated protein & RNA weighted shared nearest neighbor distance
DR+ASCp
DR-ASCp
CD19+9loASCr
CD19+9+ASCr
CD19-ASCr
Mean distance from subset
B
Robinson et al. fate-mapped ASC gene scores
Nascent
Long-lived
C
Robinson et al. fate-mapped ASC DEGs
Nascent
Long-lived
DEG
Not sig.
%+
Mean expression
Min
Max
D
Protein
### of DEPs
Higher in subset
Lower in subset
RNA
### of DEGs
Chromatin
### of DATF motifs
E
Total gene counts
### of UMIs
F
Top DEGs from select differentially enriched gene sets
Oxidative phosphorylation
Translation
AP-1 activation pathway
ATF-2 pathway
G
Transcription factor motif footprints
STAT2
IRF1
NFIX
Insertions
Subset

**Figure S6: Differential expression analyses– related to Fig. 2.**

A) Mean pairwise geodesic distances between all cells through the weighted shared nearest neighbor graph (x-axis) to each subset (y-axis), faceted by subset (panels). Data from HD BM fixed CITEseq.
B) Nascent and long-lived gene set scores by subset, as in Fig. 2B. Data from HD BM fixed CITEseq.
C) Genes underlying nascent and long-lived gene set scores by subset. Size indicates percent positive, color indicates mean expression, and outline indicates statistical significance. Data from HD BM fixed CITEseq
D) Quantification of DEPs (left), DEGs (center), and DATF motifs (right) by subset (see methods). ADT and GEX data from HD BM fixed CITEseq; ATAC data from HD BM TEAseq.
E) Unique molecular identifier (UMI) counts by subset. Diamond indicates median. Data from HD BM fixed CITEseq.
F) Top 10 DEGs from the indicated gene sets by subset. Size indicates percent positive, color indicates mean expression, and outline indicates statistical significance. Data from HD BM fixed CITEseq
G) Normalized insertions summed across the indicated transcription factor motif loci by subset (color). Data from HD BM TEAseq.

### Figure S7

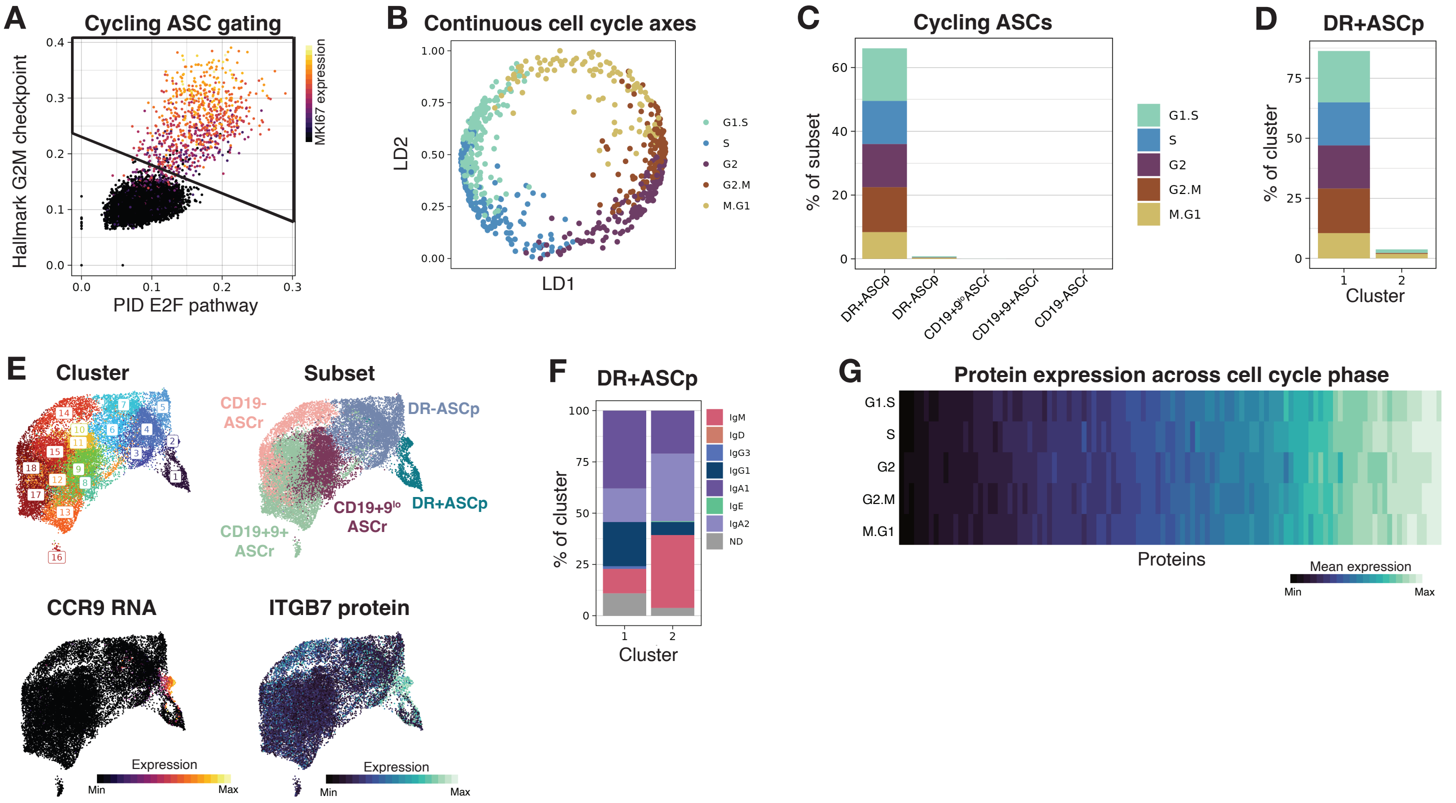
A
Cycling ASC gating
Hallmark G2M checkpoint
PID E2F pathway
MKI67 expression
B
Continuous cell cycle axes
LD1
LD2
G1.S
S
G2
G2.M
M.G1
C
Cycling ASCs
% of subset
DR+ASCp
DR-ASCp
CD19+9loASCr
CD19+9+ASCr
CD19-ASCr
D
DR+ASCp
% of cluster
Cluster
E
Cluster
Subset
CD19-ASCr
DR-ASCp
CD19+9lo ASCr
DR+ASCp
CD19+9+ ASCr
CCR9 RNA
ITGB7 protein
Expression
Min
Max
F
DR+ASCp
% of cluster
Cluster
IgM
IgD
IgG3
IgG1
IgA1
IgE
IgA2
ND
G
Protein expression across cell cycle phase
Proteins
Mean expression

**Figure S7: Only ASCp cycle and their phenotype is stable through cell cycle phases – related to Fig. 2.**

A) Gating scheme utilizing gene set scores to identify cycling cells. Data from HD BM fixed CITEseq.
B) Linear discriminant (LD) analysis derived from cell cycle phase probabilities (predictors) and cell cycle phase labels (response). Data from HD BM fixed CITEseq.
C) Percent cycling cells by subset, colored by cell cycle phase. Data from HD BM fixed CITEseq.
D) Percent cycling cells for DR+ASCp clusters, colored by cell cycle phase. Data from HD BM fixed CITEseq.
E) wnnUMAPs of ASCs colored by subset, cluster, ADT expression, and GEX expression. Data from HD BM fixed CITEseq.
F) Frequency of subisotype usage by Seurat cluster for DR+ASCp clusters. ND denotes subisotype not determined. Data from HD BM fixed CITEseq.
G) Mean expression of all surface proteins (x-axis) in cycling cells, by cell cycle phase. Data from HD BM fixed CITEseq. No statistically significant differences were observed between cell cycle phases.

### Figure S8

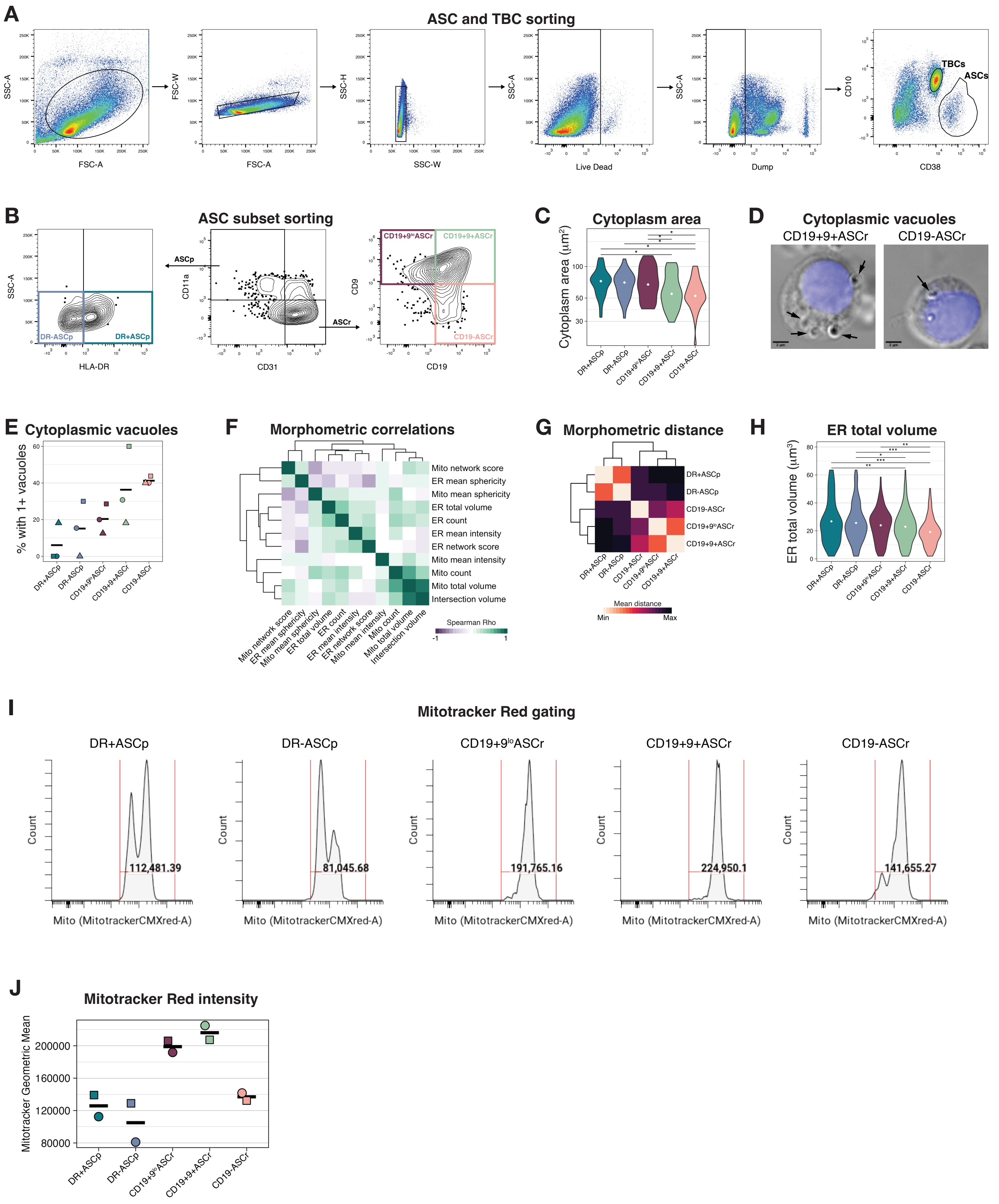
A
ASC and TBC sorting
SSC-A
FSC-A
FSC-W
SSC-H
SSC-W
Live Dead
Dump
CD10
CD38
TBCs
ASCs
B
ASC subset sorting
HLA-DR
DR-ASCp
DR+ASCp
ASCp
ASCr
CD11a
CD31
CD9
CD19
CD19+9loASCr
CD19+9+ASCr
CD19-ASCr
C
Cytoplasm area
Cytoplasm area (µm2)
D
Cytoplasmic vacuoles
CD19+9+ASCr
CD19-ASCr
E
Cytoplasmic vacuoles
% with 1+ vacuoles
F
Morphometric correlations
Mito network score
ER mean sphericity
Mito mean sphericity
ER total volume
ER count
ER mean intensity
ER network score
Mito mean intensity
Mito count
Mito total volume
Intersection volume
Spearman Rho
G
Morphometric distance
Mean distance
Min
Max
H
ER total volume
ER total volume (µm3)
I
Mitotracker Red gating
DR+ASCp
DR-ASCp
CD19+9loASCr
CD19+9+ASCr
CD19-ASCr
Count
Mito (MitotrackerCMXred-A)
112,481.39
81,045.68
191,765.16
224,950.1
141,655.27
J
Mitotracker Red intensity
Mitotracker Geometric Mean
200000
160000
120000
80000

**Figure S8: Morphometric analyses – related to Fig. 3.**

A) Exemplative gating strategy used to sort ASC and TBCs. Data from HD BM FACS.
B) Exemplative gating strategy used for sort ASC subsets from ASC gate in A. Data from HD BM FACS.
C) Cytoplasm area by subset. White dot indicates median. Q-values calculated by FDR-corrected Wilcoxon rank sum test. *Q<0.1. Data from HD BM DIC imaging.
D) Representative DIC images (60x magnification) of ASC subsets with vacuoles (arrows). Data from HD BM DIC imaging.
E) Percent of cells with 1+ vacuoles by subset. Shape indicates donor; crossbar indicates mean. Data from HD BM DIC imaging.
F) Spearman correlations of morphometric features. Data from HD BM volumetric imaging.
G) Euclidean distance between subsets calculated from morphometric feature z-scores. Data from HD BM volumetric imaging.
H) ER total volume by subset. White dot indicates median. Q-values calculated using pairwise FDR-corrected linear mixed effects model to account for donor bias. *Q<0.1; **Q<0.01; ***Q<0.001. Data from HD BM volumetric imaging.
I) Representative gating of Mitotracker Red staining in ASC subsets. Data from HD BM flow cytometry. Number indicates geometric mean of gated population.
J) Geometric mean intensity of Mitotracker Red by subset. Shapes indicate donors and crossbar indicates mean.

### Figure S9

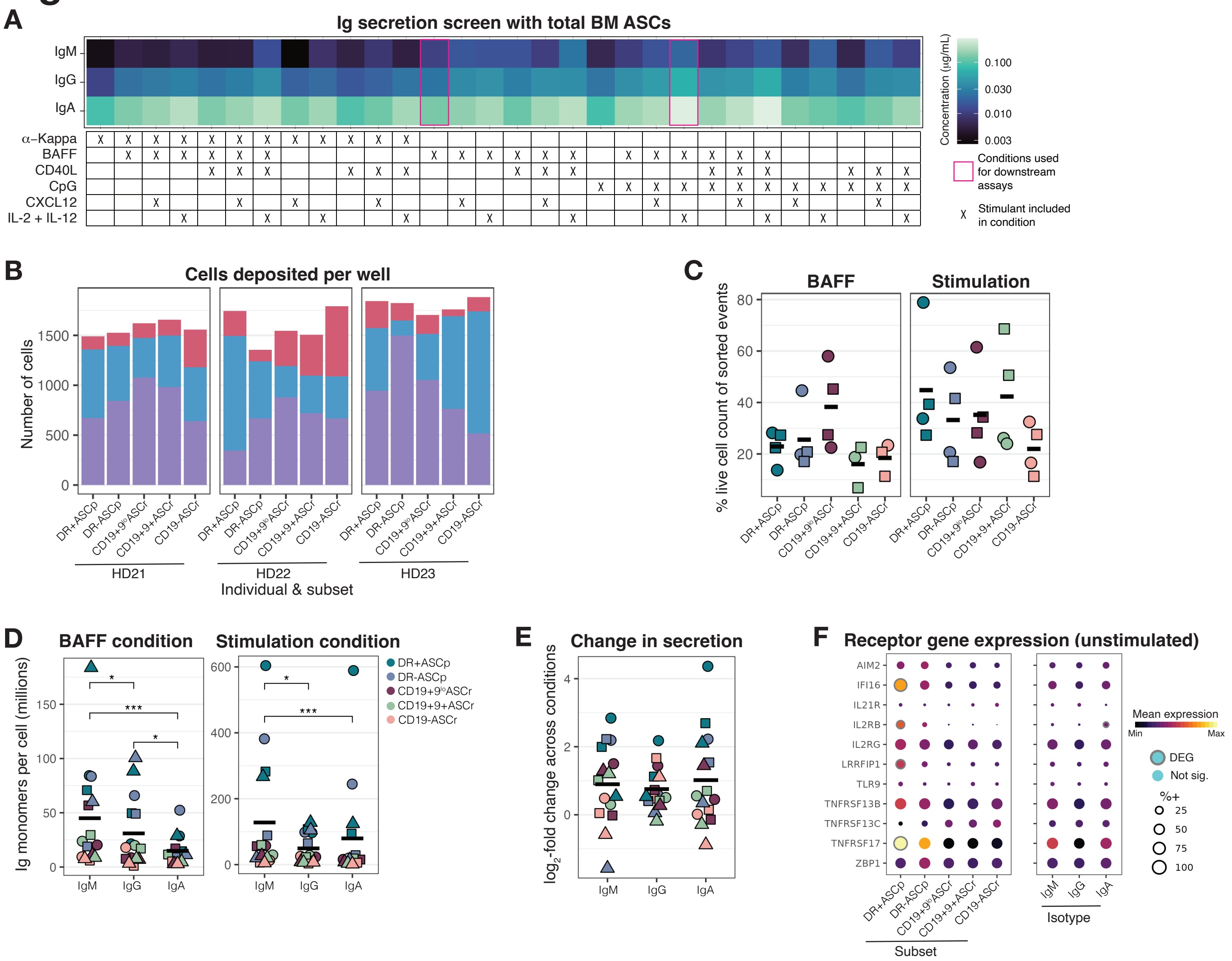

A
Ig secretion screen with total BM ASCs
IgM
IgG
IgA
α–Kappa
BAFF
CD40L
CpG
CXCL12
IL-2 + IL-12
Concentration (μg/mL)
0.100
0.030
0.010
0.003
Conditions used for downstream assays
Stimulant included in condition
B
Cells deposited per well
Number of cells
Individual & subset
HD21
HD22
HD23
C
BAFF
Stimulation
% live cell count of sorted events
D
BAFF condition
Stimulation condition
Ig monomers per cell (millions)
DR+ASCp
DR-ASCp
CD19+9loASCr
CD19+9+ASCr
CD19-ASCr
E
Change in secretion
log2-fold change across conditions
F
Receptor gene expression (unstimulated)
AIM2
IFI16
IL21R
IL2RB
IL2RG
LRRFIP1
TLR9
TNFRSF13B
TNFRSF13C
TNFRSF17
ZBP1
Mean expression
Min
Max
DEG
Not sig.
%+
25
50
75
100
Subset
Isotype

**Figure S9: Antibody secretion analyses – related to Fig. 4.**

A) Ig concentration by isotype (y-axis) from 72-hour HD BM total ASC culture supernatants in the indicated culture conditions (x-axis). Magenta boxes indicate culture conditions used in downstream assays. Data from one HD BM using the MSD isotyping panel.
B) Cells deposited by isotype (color), subset (x-axis), and individual (facet). Calculated as 2,000 sorted cells per well multiplied by the frequency of each isotype within each subset for each donor. Data from HD BM flow cytometry.
C) Live cell counts after 72 hours of cell culture as a percent of sorted events, by subset (x-axis), individual (shape), and culture condition (facet). Crossbar indicates mean. No significant differences observed (Q>0.1). Q-values calculated by Wilcoxon signed rank test with FDR correction. Two biological replicates per HD are included. Data from HD BM FACS and cell counter.
D) Mean Ig monomers secreted per cell by isotype (x-axis), subset (color), individual (shape), and culture condition (panel). Crossbar indicates mean. Q-values calculated by Wilcoxon signed rank test with FDR correction. Data from HD BM MSD isotyping. *Q<0.1; ***Q<0.001.
E) $Log_2$-fold change in Ig monomers secreted per cell from BAFF condition to stimulation condition by isotype (x-axis), subset (color), and donor (shape). Crossbar indicates mean. No significant differences observed (Q>0.1). Q-values calculated by Wilcoxon signed rank test with FDR correction. Data from HD BM MSD isotyping.
F) Mean expression (color) of receptors to molecules added to the stimulation condition, in unstimulated cells, by subset (left panel) and isotype (right panel). Size indicates percent positive and outline indicates significance. Data from HD BM fixed CITEseq.

### Figure S10

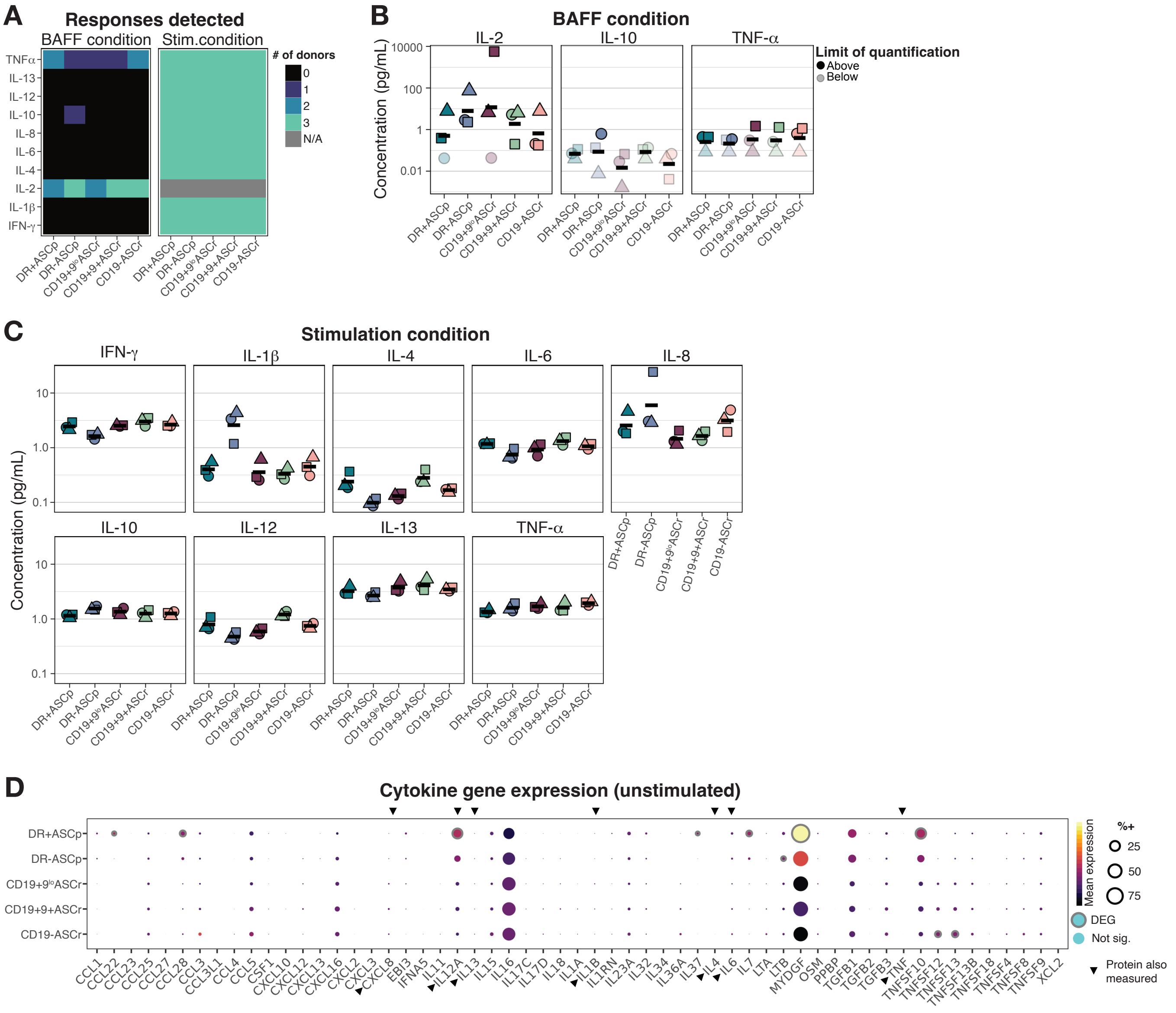
A
Responses detected
BAFF condition
Stim.condition
### of donors
0
1
2
3
N/A
TNFα
IL-13
IL-12
IL-10
IL-8
IL-6
IL-4
IL-2
IL-1β
IFN-γ
DR+ASCp
DR-ASCp
CD19+9loASCr
CD19+9+ASCr
CD19-ASCr
B
BAFF condition
IL-2
IL-10
TNF-α
Concentration (pg/mL)
10000
100
1
0.01
Limit of quantification
Above
Below
C
Stimulation condition
IFN-γ
IL-1β
IL-4
IL-6
IL-8
IL-10
IL-12
IL-13
TNF-α
Concentration (pg/mL)
10
1.0
0.1
D
Cytokine gene expression (unstimulated)
DR+ASCp
DR-ASCp
CD19+9loASCr
CD19+9+ASCr
CD19-ASCr
Mean expression
%+
25
50
75
DEG
Not sig.
Protein also measured
CCL1 CCL22 CCL23 CCL25 CCL27 CCL28 CCL3 CCL3L1 CCL4 CCL5 CSF1 CXCL10 CXCL12 CXCL13 CXCL16 CXCL2 CXCL3 CXCL8 EBI3 IFNA5 IL11 IL12A IL13 IL15 IL16 IL17C IL17D IL18 IL1A IL1B IL1RN IL23A IL32 IL34 IL36A IL37 IL4 IL6 IL7 LTA LTB MYDGF OSM PPBP TGFB1 TGFB2 TGFB3 TNF TNFSF10 TNFSF12 TNFSF13 TNFSF13B TNFSF18 TNFSF4 TNFSF8 TNFSF9 XCL2

**Figure S10: Proinflammatory cytokine expression is largely invariant across ASC subsets – related to Fig. 4.**

A) Number of individuals with cytokines (rows) detected above the lower limit of detection by subset (columns) after 72 hours of culture with BAFF (left) or a stimulation cocktail (right), consisting of BAFF, CpG, IL-2, and IL-21. Recombinant IL-2 was included in the stimulation cocktail and therefore removed for the analysis of stimulated cells. Data from HD BM MSD proinflammatory cytokine panel.

B) Supernatant cytokine concentration after BAFF culture by subset. Shapes indicate individual; crossbar indicates mean; transparency indicates below the lower limit of detection. Data were underpowered for statistical comparisons. Data from HD BM MSD proinflammatory cytokine panel.

C) Supernatant cytokine concentration after stimulation by subset. Shapes indicate individual; crossbar indicates mean; transparency indicates below the lower limit of detection. Data were underpowered for statistical comparisons. Data from HD BM MSD proinflammatory cytokine panel.

D) Mean (color) and percent positive (size) gene expression of cytokines by subisotype. Outline indicates statistical significance. Data from HD BM fixed CITEseq.

### Figure S11

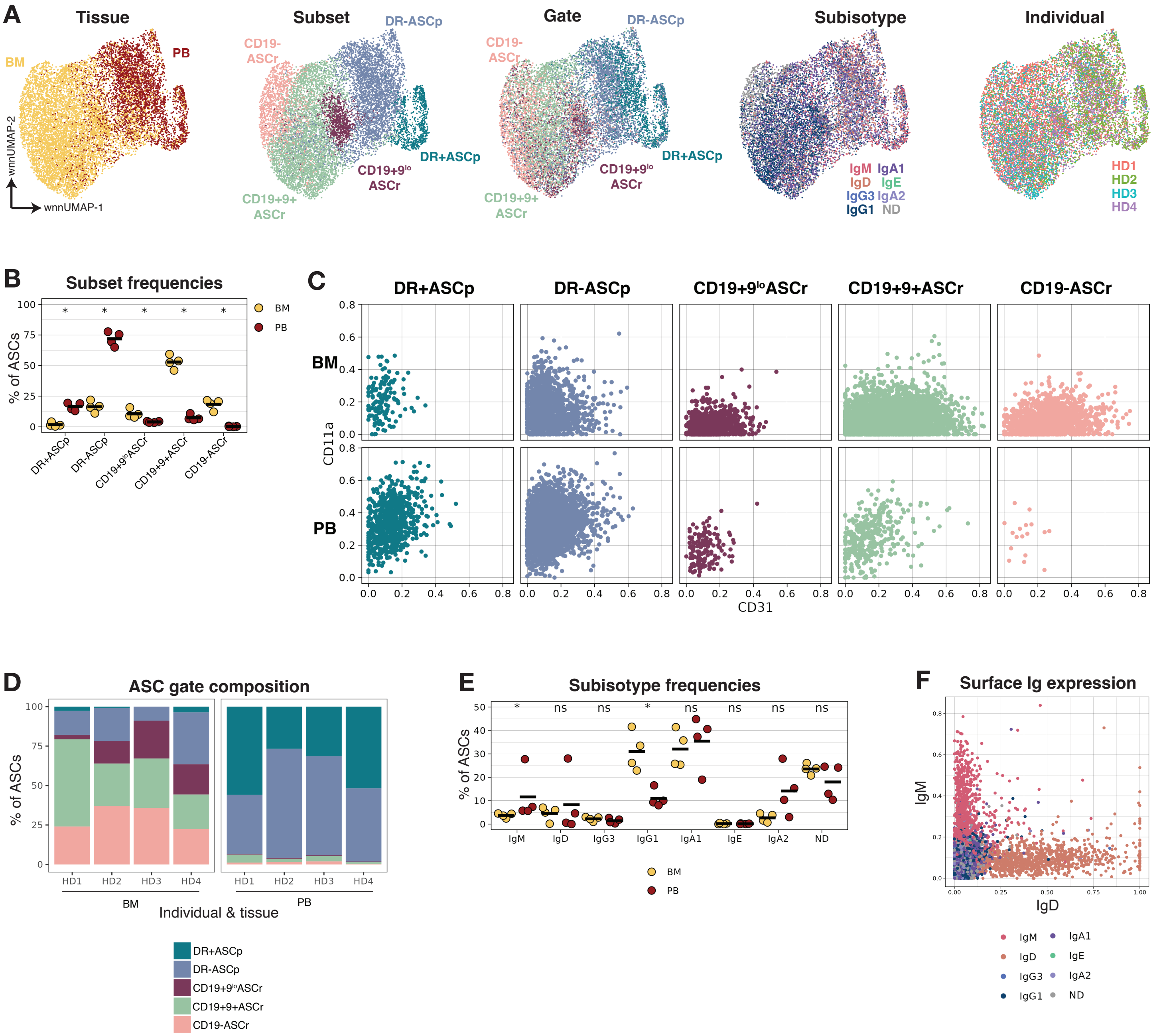

A
Tissue
BM
PB
wnnUMAP-2
wnnUMAP-1
Subset
DR-ASCp
CD19- ASCr
DR+ASCp
CD19+9lo ASCr
CD19+9+ ASCr
Gate
Subisotype
IgM IgA1
IgD IgE
IgG3 IgA2
IgG1 ND
Individual
HD1
HD2
HD3
HD4
B
Subset frequencies
% of ASCs
BM
PB
C
DR+ASCp
DR-ASCp
CD19+9loASCr
CD19+9+ASCr
CD19-ASCr
CD11a
CD31
D
ASC gate composition
Individual & tissue
E
Subisotype frequencies
F
Surface Ig expression
IgM
IgD

**Figure S11: Analysis of donor-matched tissues – related to Fig. 5**

A) wnnUMAPs of ASCs colored by various identities. Data from HD matched fixed CITEseq.
B) Percent ASC subset of total ASCs by tissue. Crossbar indicates mean. P-values calculated using Wilcoxon rank sum test. Data from HD matched fixed CITEseq. *P<0.05
C) Biaxial plots of total ASCs by tissue (rows) and subset (columns). Data from HD matched fixed CITEseq.
D) Percent ASC manual gate of total ASCs by sample. Data from HD matched fixed CITEseq.
E) Percent subisotype usage of total ASCs by tissue. Crossbar indicates mean. P-values calculated using Wilcoxon rank sum test. Data from HD matched fixed CITEseq. *P<0.05
F) Surface IgM and IgD protein expression of total ASCs colored by subisotype label, as determined by gene expression. Data from HD matched fixed CITEseq.

### Figure S12

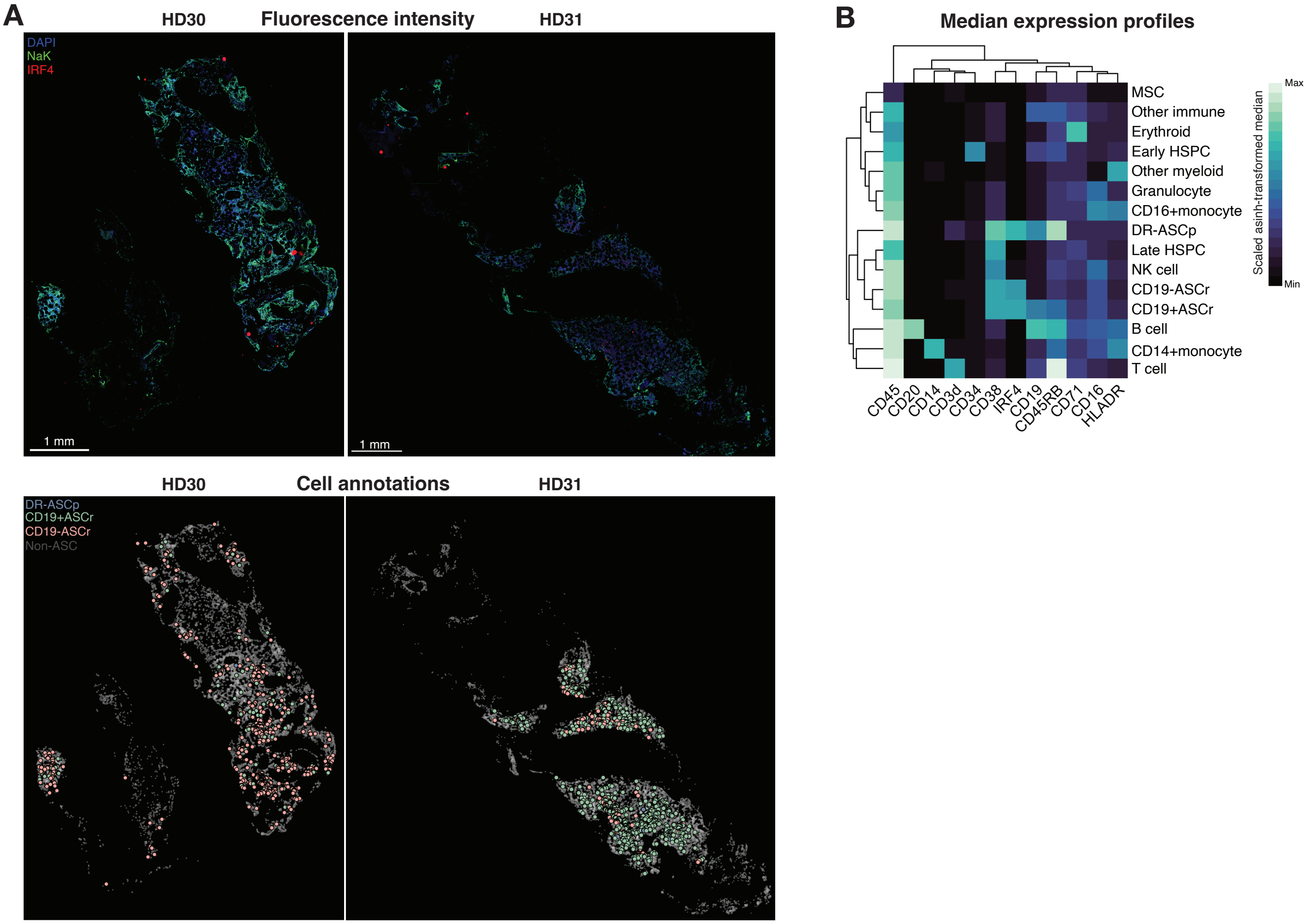

A
HD30
Fluorescence intensity
HD31
DAPI
NaK
IRF4
1 mm
1 mm
HD30
Cell annotations
HD31
DR-ASCp
CD19+ASCr
CD19-ASCr
Non-ASC
B
Median expression profiles
MSC
Other immune
Erythroid
Early HSPC
Other myeloid
Granulocyte
CD16+monocyte
DR-ASCp
Late HSPC
NK cell
CD19-ASCr
CD19+ASCr
B cell
CD14+monocyte
T cell
CD45
CD20
CD14
CD3d
CD34
CD38
IRF4
CD19
CD45RB
CD71
CD16
HLADR
Scaled asinh-transformed median
Max
Min

**Figure S12: Analysis of spatial proteomics – related to Fig. 5**

A) Full bone marrow core images colored by fluorescence signal (top) or cell subset identity localization (bottom). In lower images, ASC subset dots are outlined and expanded relative to non-ASC cells for visualization. Data from HD BM spatial proteomics.

B) Median expression of cell types. Data from HD BM spatial proteomics.

### Figure S13

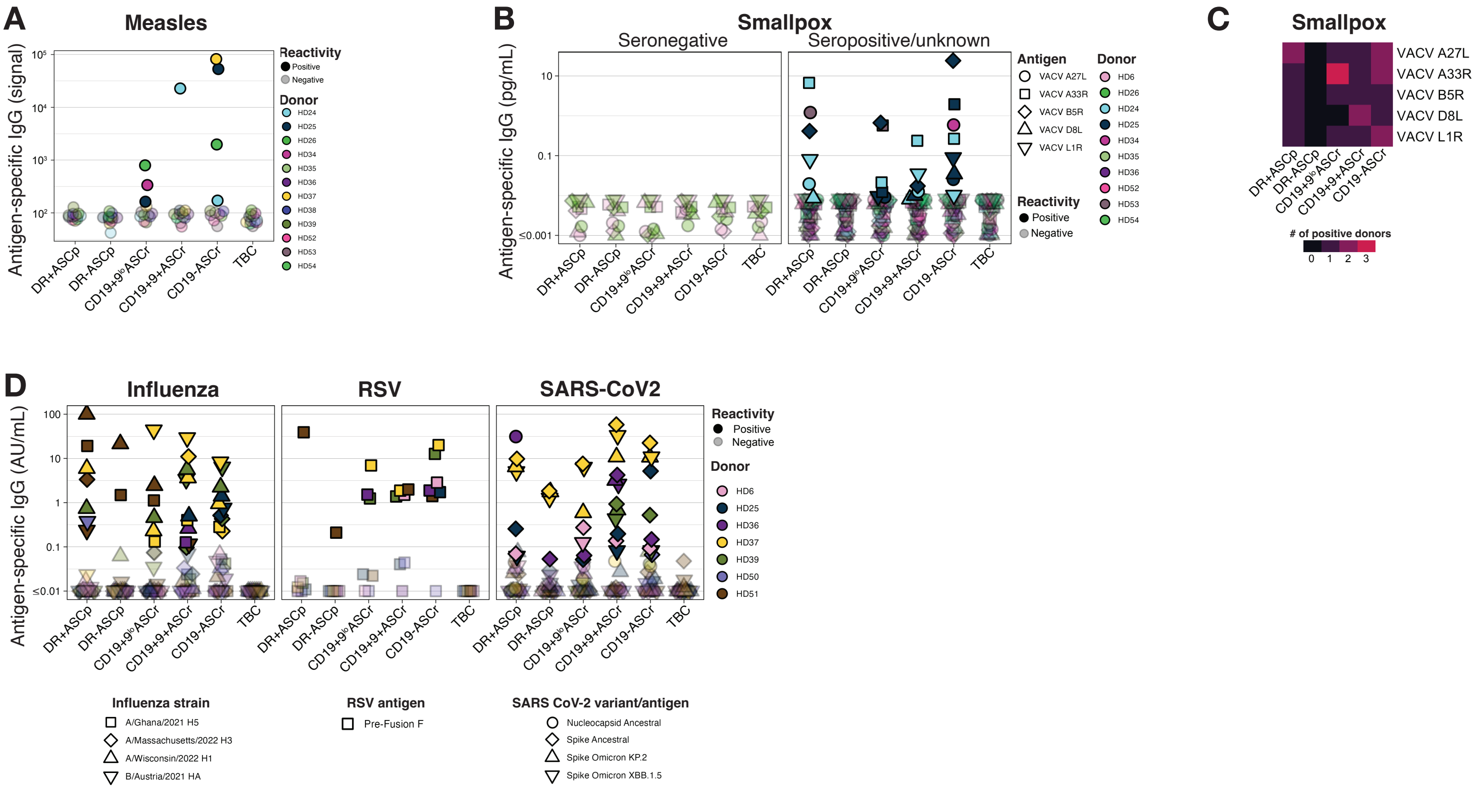
A
Measles
Antigen-specific IgG (signal)
DR+ASCp
DR-ASCp
CD19+9$^{lo}$ASCr
CD19+9+ASCr
CD19-ASCr
TBC
Reactivity
Positive
Negative
Donor
HD24
HD25
HD26
HD34
HD35
HD36
HD37
HD38
HD39
HD52
HD53
HD54
B
Smallpox
Seronegative
Seropositive/unknown
Antigen-specific IgG (pg/mL)
Antigen
VACV A27L
VACV A33R
VACV B5R
VACV D8L
VACV L1R
Donor
HD6
HD26
HD24
HD25
HD34
HD35
HD36
HD52
HD53
HD54
Reactivity
Positive
Negative
C
Smallpox
VACV A27L
VACV A33R
VACV B5R
VACV D8L
VACV L1R
### of positive donors
D
Influenza
RSV
SARS-CoV2
Antigen-specific IgG (AU/mL)
Reactivity
Positive
Negative
Donor
HD6
HD25
HD36
HD37
HD39
HD50
HD51
Influenza strain
A/Ghana/2021 H5
A/Massachusetts/2022 H3
A/Wisconsin/2022 H1
B/Austria/2021 HA
RSV antigen
Pre-Fusion F
SARS CoV-2 variant/antigen
Nucleocapsid Ancestral
Spike Ancestral
Spike Omicron KP.2
Spike Omicron XBB.1.5

**Figure S13: Analysis of antigen-specific secretion – related to Fig. 6**

A) Antigen-specific IgG signal from culture supernatants by individual (color) and subset (x-axis). TBC indicates transitional B cells, used as negative controls. Transparent dots indicate values below positivity threshold. Data from HD BM MSD measles panel.

B) Antigen-specific IgG concentration from culture supernatants of seronegative (left) and non-seronegative donors (right) by VCV antigen (shape), individual (color), and subset (x-axis). Transparent dots indicate values below positivity threshold. Data from HD BM MSD orthopoxvirus panel.

C) Number of HDs with culture supernatants positive for specificity against VCV antigens (y-axis) by subset (x-axis). Data from HD BM MSD orthopoxvirus panel.

D) Antigen-specific IgG concentration from culture supernatants by individual (color), strain/variant/antigen (shape), subset (x-axis), and virus (panels). Transparent dots indicate values below positivity threshold. Data from HD BM Respiratory panel 7.

### Figure S14

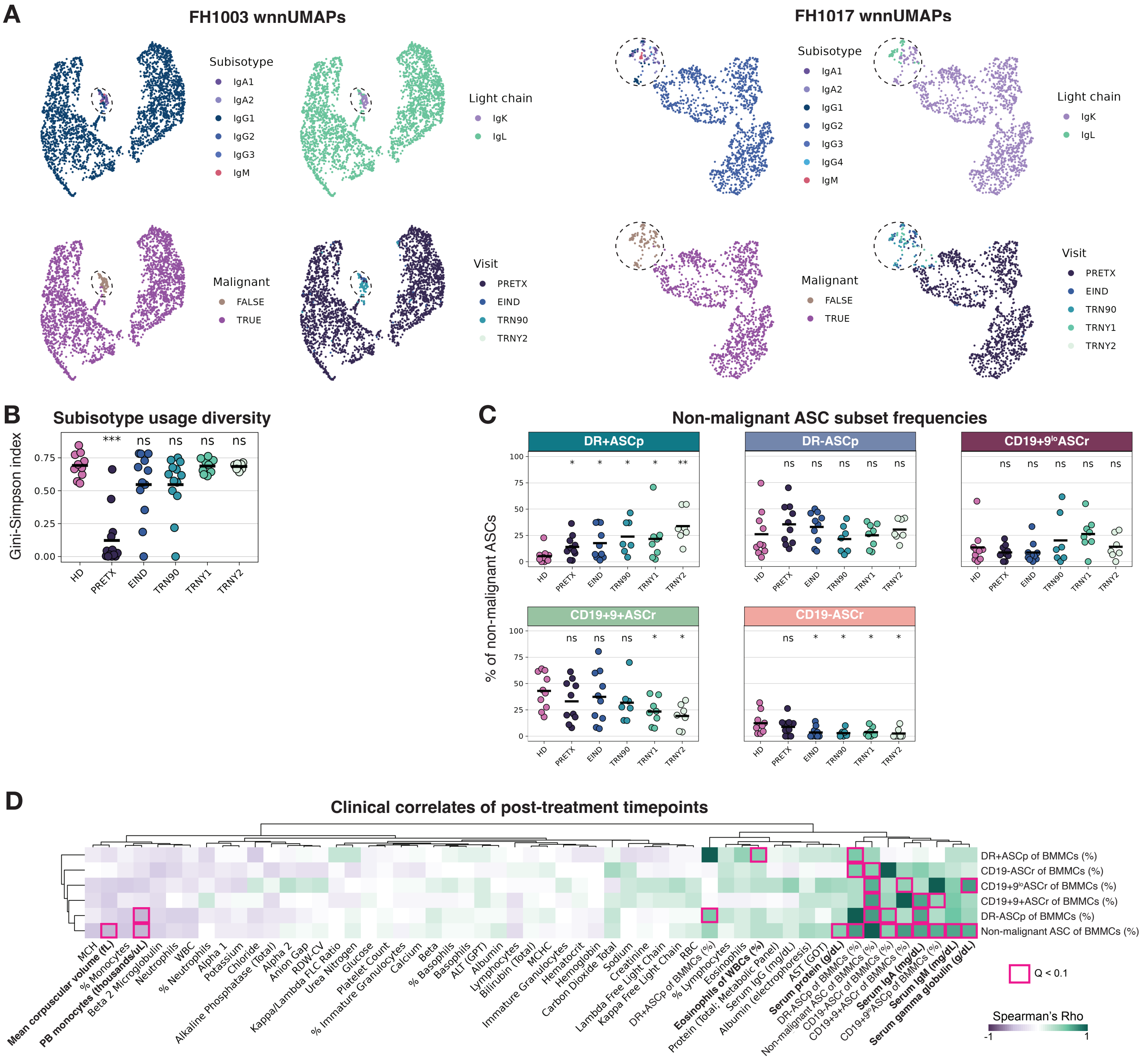

A
FH1003 wnnUMAPs
FH1017 wnnUMAPs
Subisotype
IgA1
IgA2
IgG1
IgG2
IgG3
IgG4
IgM
Light chain
IgK
IgL
Malignant
FALSE
TRUE
Visit
PRETX
EIND
TRN90
TRNY1
TRNY2
B
Subisotype usage diversity
Gini-Simpson index
HD
PRETX
EIND
TRN90
TRNY1
TRNY2
C
Non-malignant ASC subset frequencies
% of non-malignant ASCs
DR+ASCp
DR-ASCp
CD19+9loASCr
CD19+9+ASCr
CD19-ASCr
D
Clinical correlates of post-treatment timepoints
DR+ASCp of BMMCs (%)
CD19-ASCr of BMMCs (%)
CD19+9loASCr of BMMCs (%)
CD19+9+ASCr of BMMCs (%)
DR-ASCp of BMMCs (%)
Non-malignant ASC of BMMCs (%)
Q < 0.1
Spearman's Rho

**Figure S14: Analysis of longitudinal MM cohort – related to Fig. 7**

A) wnnUMAPs from two exemplative patients with MM, inclusive of all patient timepoints, colored by indicated label. Dashed circles indicate the location of non-malignant cells. Each patient's wnnUMAPs were generated independently and represent different manifolds. Data from MM longitudinal BM CITEseq.
B) Gini-Simpson index (diversity) of subisotypes used by total ASCs for HDs (left) and patients with MM, by timepoint (x-axis). Low diversity at PRETX timepoints is due to the high abundance of malignant clonotype. Crossbar indicates mean. Q-values calculated by Wilcoxon rank sum test with FDR correction, comparing to HDs. Data from MM longitudinal BM CITEseq. ***Q<0.001
C) Percent subset of non-malignant ASCs for HDs (left) and patients with MM, by timepoint (x-axis) and subset (panels). Crossbar indicates mean. Q-values calculated by Wilcoxon rank sum test with FDR correction, comparing to HDs. Data from MM longitudinal BM CITEseq. *Q<0.1; **Q<0.01.
D) Correlation between non-malignant ASC total and subset frequencies (y-axis) and various clinical tests taken at any post-treatment timepoint (x-axis). Outlines indicated statistical significance. Correlated clinical features are in bold. Correlation was calculated using the Spearman method and p-values were corrected by FDR. Data from MM longitudinal BM CITEseq and clinical tests.

### Figure S15

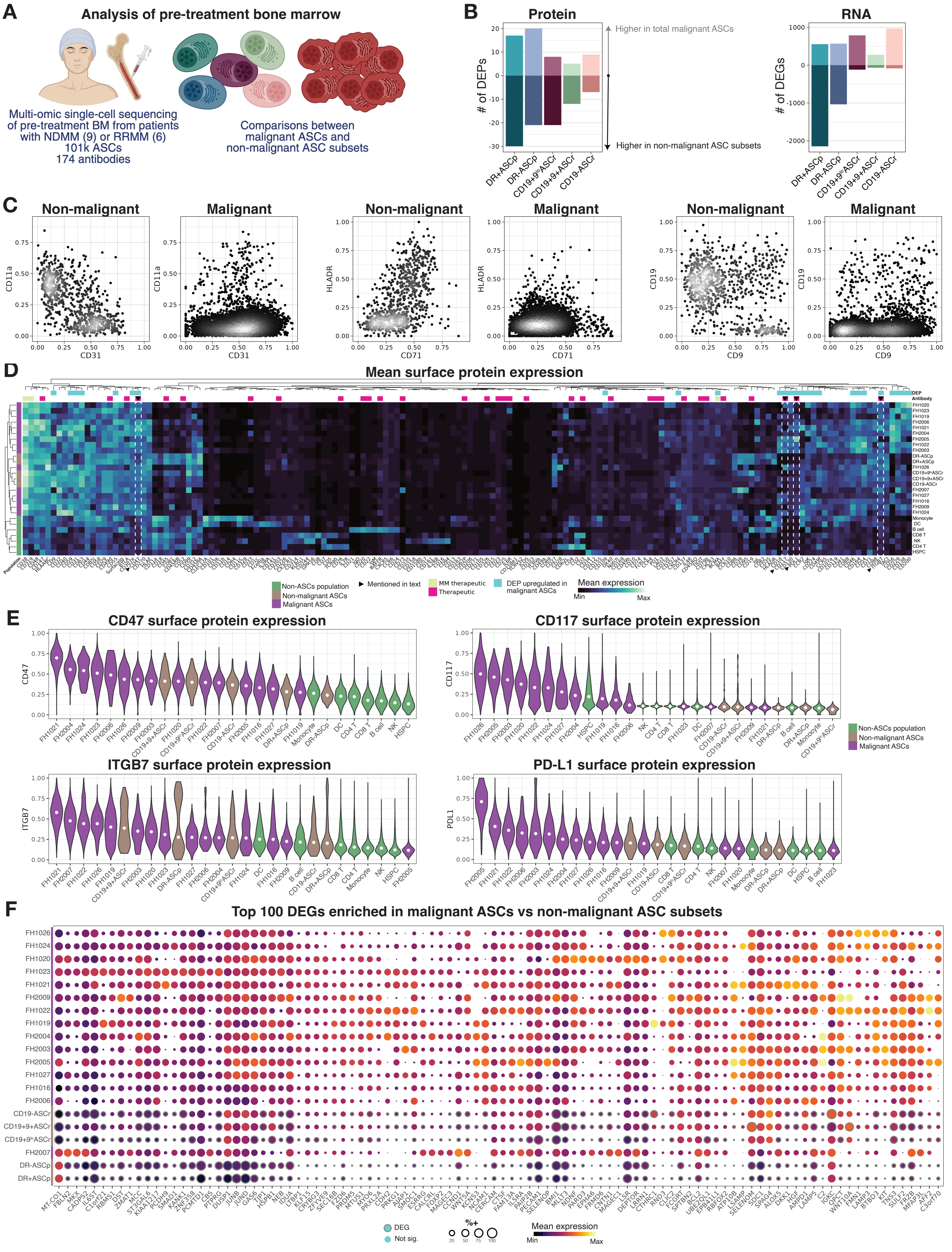
A
Analysis of pre-treatment bone marrow
Multi-omic single-cell sequencing of pre-treatment BM from patients with NDMM (9) or RRMM (6)
101k ASCs
174 antibodies
Comparisons between malignant ASCs and non-malignant ASC subsets
B
Protein
RNA
### of DEPs
### of DEGs
Higher in total malignant ASCs
Higher in non-malignant ASC subsets
DR+ASCp
DR-ASCp
CD19+9loASCr
CD19+9+ASCr
CD19-ASCr
C
Non-malignant
Malignant
CD11a
CD31
HLADR
CD71
CD19
CD9
D
Mean surface protein expression
DEP
Antibody
Population
Non-ASCs population
Non-malignant ASCs
Malignant ASCs
Mentioned in text
MM therapeutic
Therapeutic
DEP upregulated in malignant ASCs
Mean expression
Min
Max
E
CD47 surface protein expression
CD117 surface protein expression
ITGB7 surface protein expression
PD-L1 surface protein expression
CD47
CD117
ITGB7
PDL1
Non-ASCs population
Non-malignant ASCs
Malignant ASCs
F
Top 100 DEGs enriched in malignant ASCs vs non-malignant ASC subsets
DEG
Not sig.
%+
25
50
75
100
Mean expression
Min
Max

**Figure S15: Malignant ASCs display aberrant expression profiles – related to Fig. 7**

A) Experimental diagram.
B) Quantification of DEPs (left) and DEGs (right) comparing total malignant cells to non-malignant ASC subsets (x-axis). Data from MM PRETX BM fixed CITEseq.
C) Biaxial plots of surface proteins used for subset gating in non-malignant and malignant ASCs. Data from MM PRETX BM fixed CITEseq.
D) Mean surface protein expression of total ADTs in patient myeloma cells (annotated with patient IDs; purple sidebar), other immune cell types (green sidebar), and non-malignant ASC subsets (brown sidebar). DEPs indicated with blue boxes above heatmap. Proteins that are currently clinical therapeutic targets are indicated above in magenta or light yellow if indicated for myeloma patients. Data from MM PRETX BM fixed CITEseq.
E) Surface protein expression of indicated ADTs in patient myeloma cells (annotated with patient IDs; purple), other immune cell types (green), and non-malignant ASC subsets (brown). Data from MM PRETX BM fixed CITEseq.
F) Mean expression (color) and percent positive (size) of top 100 DEGs enriched in malignant cells compared to non-malignant ASC subsets. Outline on normal ASC subset bubbles indicates statistical significance. Myeloma cells are indicated by patient ID and purple sidebar. Data from MM PRETX BM fixed CITEseq.
